# D-Cycloserine as Adjunct to Brief Computerised CBT for Spider Fear: Effects on Fear, Behaviour, and Cognitive Biases

**DOI:** 10.1101/602276

**Authors:** Nils Kappelmann, Mareike Suesse, Susann Steudte-Schmiedgen, Reinoud Kaldewaij, Michael Browning, Tanja Michael, Mike Rinck, Andrea Reinecke

**Author notes:** Correspondence: Andrea Reinecke, Department of Psychiatry, Warneford Hospital, Oxford OX37JX, UK.

## Abstract

In anxiety disorders, cognitive behavioural therapy (CBT) improves information-processing biases such as implicit fear evaluations and avoidance tendencies, which predicts treatment response, so they might constitute important treatment targets. This study investigated (i) whether information-processing biases changed following single-session computerised CBT for spider fear, and (ii) whether this effect could be augmented by administration of D-cycloserine (DCS). Spider-fearful individuals were randomized to receiving 250mg of DCS (n=21) or placebo (n=17) and spider fear was assessed using self-report, behavioural, and information-processing (Extrinsic Affective Simon Task & Approach Avoidance Task) measures. Linear mixed-effects analyses indicated improvements on self-report and behavioural spider fear following CBT, but not on cognitive bias measures. There was no evidence of an augmentation effect of DCS on any outcome. Cognitive biases at 1-day were not predictive of 1-month follow-up spider fear. These findings provide no evidence for information-processing biases relating to CBT response or augmentation with DCS.

## 1. Introduction

Anxiety disorders are effectively treated with exposure therapy (Hofmann & Smits, 2008; Stewart & Chambless, 2009). However, treatments are costly, difficult to access, half of the patients do not respond, and the anxiety-provoking nature of exposure techniques can result in patients terminating treatment prematurely (Choy, Fyer, & Lipsitz, 2007; Emmelkamp et al., 2002; Gunter & Whittal, 2010; Wang et al., 2006). This calls for better treatment protocols offering faster, yet still effective, care. To arrive at such treatment protocols, researchers need to better understand which processes underlie successful therapy and integrate approaches from psychology and neuroscience to arrive at an effective, contemporary therapy that synergistically bridges disciplines.

Recent work has suggested that one mechanistic driver of improvement in clinical symptoms might be changes in information-processing biases to threat. In a waiting-list controlled study investigating single-session CBT for panic disorder, attentional bias towards threat stimuli improved one day post-treatment and prior to any symptomatic changes. These early reductions in attentional bias towards threat were found to predict better symptomatic improvement at one-month follow-up (FU), explaining about 50% of the variance in symptom change (Reinecke, Waldenmaier, Cooper, & Harmer, 2013). Other findings also include changes in attentional biases following treatment in specific phobia (van den Hout, Tenney, Huygens, & De Jong, 1997), generalised anxiety disorder (Mogg, Bradley, Millar, & White, 1995), and social anxiety disorder (Calamaras, Tone, & Anderson, 2012).

Beyond attentional bias towards threat, other information-processing biases such as implicit threat evaluation and threat avoidance could be similarly relevant to treatment change. These biases can be quantified in computerised tasks, for instance, by assessing differences in individuals’ reaction time for congruent (unpleasant word & fear stimulus) versus incongruent (pleasant word & fear stimulus) stimuli pairings or speed of approaching versus avoiding threatening versus neutral stimuli, respectively. Exaggerated implicit threat evaluation was found to decrease following CBT in generalised anxiety disorder (Reinecke, Rinck, Becker, & Hoyer, 2013; Teachman & Woody, 2003), in panic disorder (Teachman, Marker, & Smith-Janik, 2008), and, together with reductions in avoidance tendencies, in specific phobia (Reinecke, Soltau, Hoyer, Becker, & Rinck, 2012). These findings correspond to cognitive theories of treatment action, which propose that specific anxiety is characterised by initial hypervigilance towards threat stimuli (e.g., via attentional allocation to the feared object or implicit fear evaluation) and subsequent attentional avoidance of the phobic object or situation (Amir, Foa, & Coles, 1998; Cisler & Koster, 2010; Mogg & Bradley, 2006). Following this reasoning, these information-processing biases need to be successfully targeted early on during treatment as they may precede meaningful symptomatic improvements. Two studies on implicit threat evaluation have already demonstrated that improvements in this processing bias are correlated with symptomatic improvements (Teachman et al., 2008) and predicted additional symptom change during treatment FU (Reinecke, Rinck, et al., 2013). Thus, improvements in information-processing biases might provide an important, early constituent of treatment change, so that research needs to establish whether and how these biases are changed following interventions and whether they predict clinical improvements.

One promising method for facilitating the alteration of information-processing biases could be pharmacological augmentation of CBT with D-cycloserine (DCS). DCS is a partial agonist of the *N*-methyl-D-aspartate (NMDA) receptor and, based on the NMDA receptor’s role in learning and memory (Zorumski & Izumi, 2012), has been proposed to enhance extinction learning of the fear association in exposure therapy. While results from RCTs have been mixed, meta-analyses suggest that DCS offers an effective augmentation strategy for anxiety, obsessive-compulsive, and post-traumatic stress disorders if used correctly (Mataix-Cols et al., 2017; McGuire, Wu, Piacentini, McCracken, & Storch, 2017). Here, factors such as exposure duration seem relevant in that shorter CBT protocols around 2-5 sessions (Guastella et al., 2008; Hofmann et al., 2006; Otto et al., 2010; Ressler et al., 2004) were more effective than longer protocols (de Kleine, Hendriks, Kusters, Broekman, & van Minnen, 2012; Hofmann et al., 2013; Kushner et al., 2007; Storch et al., 2007; Wilhelm et al., 2008). Some studies using longer protocols also report DCS benefits in early sessions only (Hofmann et al., 2013; Kushner et al., 2007; Wilhelm et al., 2008). This suggests that drug enhancement of extinction learning might be short-lived and confined to an initial acceleration of therapy. To make the most out of intensive and drop out-prone therapy, DCS might thus best be combined with brief CBT designs such as the one-session computer-based exposure therapy developed by Muller and colleagues (2011). If studies demonstrated DCS effects in such cost-effective CBT designs, this could pave the way for less expensive and more accessible anxiety treatment.

The purpose of the present study was to investigate (i) whether implicit fear evaluations and avoidance tendencies were indeed changed following CBT, and (ii) whether DCS could augment any CBT-induced modification of these information-processing biases, compared with placebo augmentation. We also wanted to explore whether changes in information-processing biases could predict spider fear at FU. Spider-fearful individuals were treated with one-session CBT while being randomised to treatment augmentation with single-dose DCS or placebo. We measured implicit fear evaluations, avoidance, and spider fear at baseline, one-day, and one-month FU. Our hypotheses were that (i) implicit fear evaluation and avoidance tendencies would be reduced at one-day and one-month FU following CBT, and (ii) DCS would facilitate CBT-induced changes in information-processing biases.

## 2. Methods and Materials

### 2.1. Participants

Formal sample size calculations were limited by the lack of evidence regarding the effect of DCS-augmented CBT on cognitive outcome measures. We estimated sample sizes based on a medium-sized (f=.25) 2×2 mixed-factors interaction of DCS-group and time, p=.05, correlation among repeated measures of r=.50, and power of 1-ß=.80, the power analysis with G*Power (Faul, Erdfelder, Buchner, & Lang, 2009) yielded a necessary total sample size of 34 participants.

To allow for dropouts, 40 spider-fearful individuals were recruited through advertisements on posters, flyers, community websites, and in newspapers and compensated for participation. Inclusion criteria were age 18-80 years, non-smoker or smoking less than 5 cigarettes per day, no use of psychoactive medication in the previous six weeks, a body mass index (BMI) between 18 and 30 kg/m^2^, and a score of 14 or higher on the Spider Anxiety Screening (SAS; Rinck et al., 2002) at pre-screening and baseline. Exclusion criteria were pregnancy or breast-feeding; current use of DCS, ethionamide or isoniazid; lifetime history of bipolar disorder, psychosis, alcohol, medication or drug abuse or dependence, or a current primary depressive disorder as assessed using the Structured Clinical Interview for DSM-IV (SCID); a first-degree family member with a history of severe psychiatric disease; lifetime history of severe physical illness; previous exposure-based CBT for spider fear; and inadequate English skills. It was also assessed whether participants qualified for a diagnosis of Specific Phobia using the SCID for DSM-IV. This study received approval from the Oxford University research ethics committee and all participants provided written consent for participation in the study.

### 2.2. Study Design

This single-centre study used a double-blind parallel design, where participants were randomised to receiving a single dose of 250 mg cycloserine (King’s Pharmaceuticals) or placebo (microcrystalline cellulose, Rayonex GmbH). Participants and the researcher responsible for treatment and data collection remained naïve to drug group allocation. Placebo was encapsulated in shells identical to DCS, and capsules were administered to participants in identical containers. Generation of the randomisation sequence, treatment allocation and drug dispense in identical capsule shells and bottles were executed by a researcher not in direct contact with study participants. The randomization sequence was generated using a random number generator, and was based on blocked randomization (blocks of four) while stratifying for gender. Of the total sample of 38 participants, 21 were randomized to DCS and 17 to placebo, of which 13 and 7 qualified for a diagnosis of SP (*χ*^2^ (1)=1.62, p=.20), respectively (see Table 1).

After an initial screening visit when inclusion and exclusion criteria were assessed, participants were invited for the treatment day. Here, they completed clinical anxiety and bias measures at baseline, which led to exclusion of two participants who were below the SAS<14 cut-off. Afterwards, they received the exposure-based CBT treatment, which was followed by another questionnaire-based assessment of their spider fear. All participants were then invited for one-day and one-month FU assessments when they completed the whole assessment battery again. Study flow including three missing data points due to technical errors as well as 1 induced missing data point due to high error rate on AAT is described in Figure 1.

**Figure 1.**
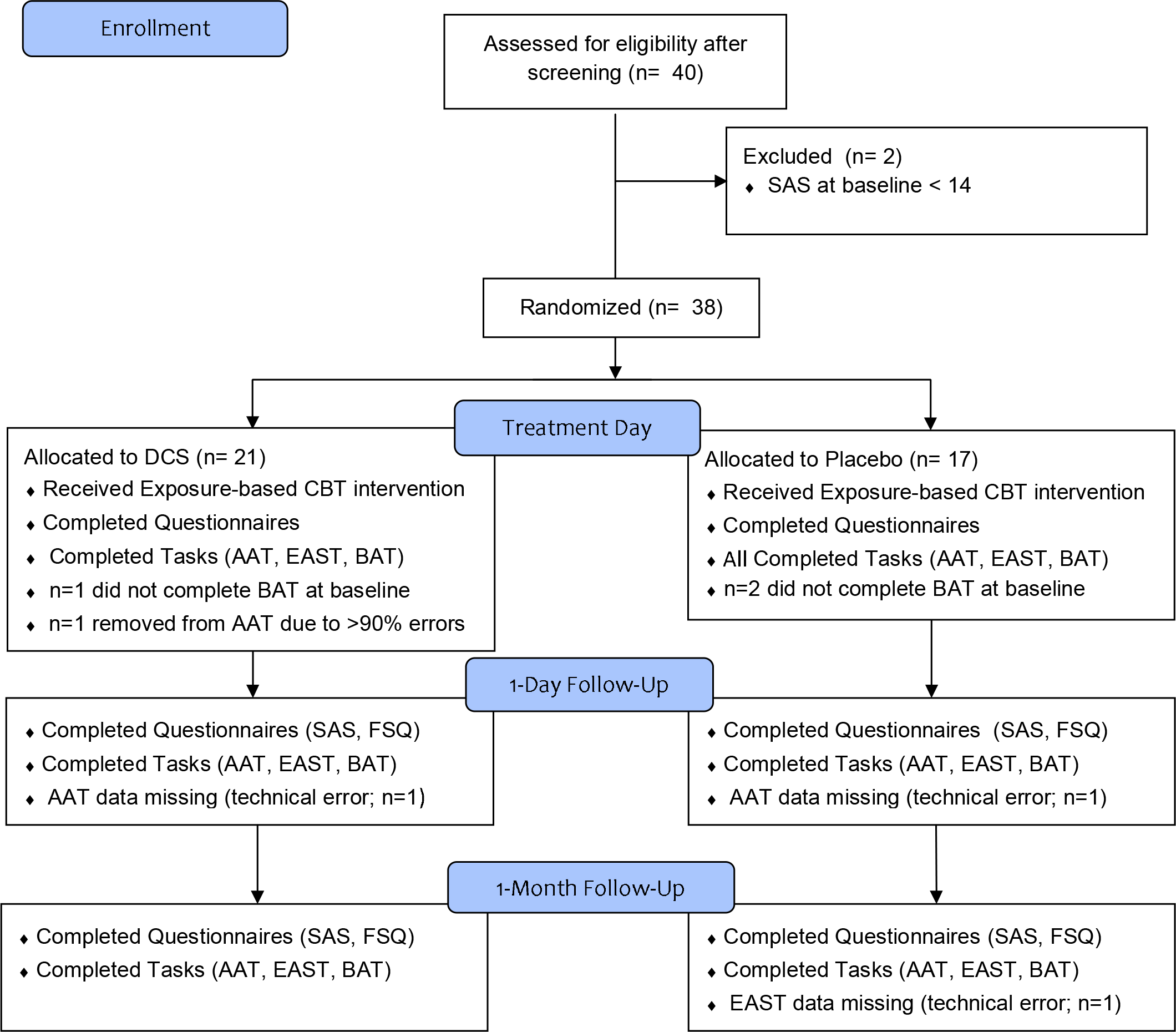
Adapted Consort Study Flow Diagram.

### 2.3. Clinical Anxiety Measures

Spider fear was measured using the questionnaires Fear of Spiders Questionnaire (FSQ; Szymanski & O’Donohue, 1995) and Spider Anxiety Screening (SAS; Rinck et al., 2002). Additionally, we assessed spider fear in a behavioural approach test (BAT) in which participants were asked to quickly approach a cage with two tarantula carapaces on a windowsill. We measured speed of approach as distance divided by time (m/s). This fear-relevant task has been used in previous studies and demonstrated good 1-week test-retest reliability (r=0.84, p<.001; Reinecke, Becker, & Rinck, 2010; Reinecke et al., 2012).

### 2.4. Threat Bias Measures

Threat bias was assessed using the Extrinsic Affective Simon Task (EAST) and the Approach Avoidance Task (AAT) with fear-related stimuli that were used in previous research (Reinecke et al., 2010, 2012; Rinck & Becker, 2007) and demonstrated sufficient reliability (1-week test–retest reliability: EAST: r = .42, p < .001; AAT: r = .35, p < .01; internal consistency: EAST: Cronbach’s α = .44, AAT: Cronbach’s α = .66) and validity (correlation with FSQ: EAST: r = −.32, p < .01, AAT: r = −.59, p < .001).

#### 2.4.1. EAST

Stimuli for the EAST were either words in font size 24 or pictures (300×400 pixels). There were 20 word stimuli with pleasant (e.g., happiness, pleasure) or unpleasant valence (fear, dangerous) and 10 picture stimuli with spider or butterfly photographs (300×400 pixels). Pleasant and unpleasant as well as spider and butterfly stimuli were equally represented among words and pictures, respectively, and all picture stimuli were present in original and mirrored form (the animal’s gaze points left and right once). The EAST was divided into 2 practice (i.e., valence practice and target practice) and 5 experimental blocks and feedback on mistakes was provided throughout. During the valence practice block, participants were instructed to categorise words based on valence with a left and a right response key in a counterbalanced order. In a total of 80 valence practice trials with each word presented 4 times, participants learned to associate both sides with a specific valence. In a subsequent target practice block, participants categorised 5 pictures of dragonflies (twice in original and twice in mirrored form) based on their gaze direction as left or right. In the five experimental blocks, participants had to categorise 40 words and 40 pictures based on valence and gaze direction, respectively, in a pseudorandom order. This resulted in compatible trials in which butterflies were associated with positive valence and spiders with negative valence but also in incompatible trials in which spiders were paired with positive valence and butterflies with negative valence. The fear evaluation effect was computed based on the difference in reaction times (RTs) between compatible and incompatible trials of spider pictures.

#### 2.4.2. AAT

Stimuli for the AAT consisted of butterfly and spider pictures (8 each), presented in portrait (400×300 pixels) and landscape format (300×400 pixels), and portrait- or landscape-shaped rectangles as control stimuli. Participants were asked to respond to stimuli based on shape only (i.e., portrait or landscape) by either pushing or pulling the joystick. Response direction and format were counterbalanced between participants. Pushing and pulling of the joystick resulted in images gradually shrinking or enlarging in size, respectively. After 18 practice trials consisting of control stimuli only, participants completed two experimental blocks with 160 trials divided into 64 spider trials, 64 butterfly trials, and 32 control trials with equal distribution of picture formats and pseudorandom order. Adding to this, experimental blocks included 32 further control trials each at beginning and end needed for calibration of joystick reactions. Outcome was the subtracted RT of fear-incompatible (pulling spiders) from fear-compatible (pushing spiders) trials corrected for general response tendencies with the joystick (pulling butterflies subtracted from pushing butterflies).

### 2.5. Treatment

#### 2.5.1. Exposure-based CBT

45-minute treatment involved a combination of the following components: i) Psychoeducation, including written information about the anxiety response, the relevance of avoidance and escape behaviour, and overwriting the fear association by exposure. ii) Computer-based exposure, involving participants visually exploring seven large spider pictures that were presented for 3 minutes each. To facilitate visual attention towards the spider, participants had to click on star symbols superimposed on the images. This approach was validated in previous research and had shown that it decreased self-report (i.e., FSQ) and behavioural (i.e., BAT) spider fear post-treatment and until 1-month FU when compared to non-treatment controls (Müller et al., 2011). iii) Exposure to a dead spider, guided by the therapist and with the goal of reducing initial fear levels to at least 50% of baseline fear. All diagnostic assessments and treatments were supervised by an experienced clinical psychologist (AR).

### 2.6. Statistical Analyses

To answer our hypotheses on efficacy of CBT and augmentation with DCS, respectively, we conducted linear mixed-effects analyses, which allow missing data in outcome variables (EAST, AAT, SAS, FSQ, & BAT) instead of list-wise deletion. To statistically test predictors of linear mixed-effects models, two statistical models with and without predictor of interest are created and formally tested against each other using likelihood-ratios (*χ*^2^) tests. Initially, a basic model with effects for time and a random intercept was tested against the same model with random slope for time. Likelihood-ratio testing of the random slope determined whether it was taken forward for analyses of fixed effects. CBT efficacy was tested using likelihood-ratio test of time variables. DCS efficacy was tested by including group and group*time effects in the model and testing inclusion of the interaction with likelihood-ratio test.

We also explored whether cognitive bias measures could predict spider fear indices. To this end, we computed correlations of measures at 1-day and 1-month FU and further used multiple linear regression analyses of spider fear levels (i.e., SAS, FSQ, BAT) at 1-month FU on 1-day cognitive bias measures (EAST, AAT). Importantly, regression analyses were adjusted for respective spider fear levels at 1-day FU.

Statistical analyses were conducted in *R* (R Core Team, 2017) and the *lme4* package was used for linear mixed-effects modelling (Bates, Maechler, Bolker, & Walker, 2013). Assumptions were evaluated by visual inspection of normality and residuals. The impact of outliers on linear mixed-effects analyses was further evaluated using the *influence.ME* package (Nieuwenhuis, te Grotenhuis, & Pelzer, 2012), which indicated that the pattern of results was stable against outlying observations. Scripts and results of analyses are available as Online Supplementary Material.

## 3. Results

### 3.1. Group Matching and Drug Side Effects

Comparison of baseline characteristics in DCS and placebo groups (see Table 1) did not reveal any differences in demographics, clinical characteristics, or subjective and behavioural measures of spider fear. There was also no difference in avoidance as measured by the AAT, but participants of the DCS group showed significantly greater levels of implicit fear evaluation of spiders as measured by the EAST. As is visible from Figure 2, however, this baseline difference seemed to result from an outlying observation as confirmed by sensitivity analysis without this outlier (p=.15).

**Table 1.** Baseline characteristics of DCS and placebo groups.

|  | DCS (n=21) | Placebo (N=17) | p value<sup>a</sup> |
| --- | --- | --- | --- |
| <u>Demographics</u> |  |  |  |
| Females, *N* (%) | 16 (76.19) | 14 (82.35) | .643 |
| Age in years, *M* (*SD*) | 26.67 (5.97) | 26.88 (7.79) | .924 |
| Education in years, *M* (*SD*) | 16.76 (2.07) | 16.65 (4.85) | .922 |
| Verbal intelligence (NART), *M* (*SD*) | 116.40 (5.27) | 113.00 (5.77) | .091 |
| Native English, *N* (%) | 18 (85.71) | 14 (82.35) | .731 |
| Ethnicity (Caucasian), *N* (%) | 18 (85.71) | 16 (94.12) | .154 |
| Smoking, *N* (%) | 2 (11.76) | 2 (9.52) | .823 |
| Drinking (units/week), *M* (*SD*) | 5.60 (6.46) | 5.85 (4.52) | .890 |
| Comorbid lifetime diagnoses, *N* (%) |  |  |  |
| Major Depressive Disorder | 1 (4.76) | 1 (5.88) |  |
| Specific Phobia (dogs) | - | 1 (5.88) |  |
| Social Phobia | 1 (4.76) | - | .558 |
| <u>Anxiety Measures</u> |  |  |  |
| Spider Phobia diagnosis, *N* (%) | 13 (61.90) | 7 (41.18) | .203 |
| SAS, *M* (*SD*) | 20.14 (2.76) | 19.47 (2.94) | .473 |
| FSQ, *M* (*SD*) | 70.86 (13.18) | 66.88 (16.32) | .412 |
| BAT (n=35), *M* (*SD*) | 0.27 (0.20) | 0.18 (0.14) | .190 |
| STAIT, *M* (*SD*) | 35.57 (9.78) | 33.65 (7.52) | .509 |
| <u>Fear Information Processing</u> |  |  |  |
| EAST spider evaluation, *M* (*SD*) | -17.36 (39.10) | -2.53 (39.59) | .022* |
| AAT spider avoidance (n=37), *M* (*SD*) | -15.12 (123.84) | -7.27 (81.02) | .818 |

*Note:* <sup>a</sup> p values are based on $\chi^2$ -test for categorical and independent samples, and on t-test for continuous data. <sup>b</sup> These physiological measures reflect the difference score in standing versus sitting position. \* $p < .05$ .

When asked for their beliefs about the allocated group, 16 participants thought they received DCS (7 were correct), 15 participants thought they received placebo (5 were correct), and 7 indicated they had no idea. Consequently, there was no evidence for participants correctly inferring their treatment group as supported by a *χ*^2^-test (*χ*^2^(2)=1.66, p=.437). In a similar vein, experimenters correctly classified 8 out of 12 participants as receiving DCS, 2 out of 8 as receiving placebo, and indicated ‘no idea’ about the allocated group for 18 participants, suggesting the experimenters were not able to correctly identify group allocation (*χ*^2^(2)=3.84, p=.146).

Physiological measures and side effects, as indexed by visual analog scale (VAS) ratings, were assessed at baseline and drug peak levels (Table 2). Repeated measures ANOVA revealed no significant differences in change between groups on physiological measures and VAS ratings of alertness, anxiousness, depressiveness, dizziness, flushness, hopelessness, nausea, sadness, sleepiness, tachycardia, or tearfulness (all p>.1). Participants in the DCS compared to placebo groups, however, became significantly dizzier (p=.011), more flushed (p=.014), and reported greater tremor (p=.035) at drug peak levels.

**Table 2.**
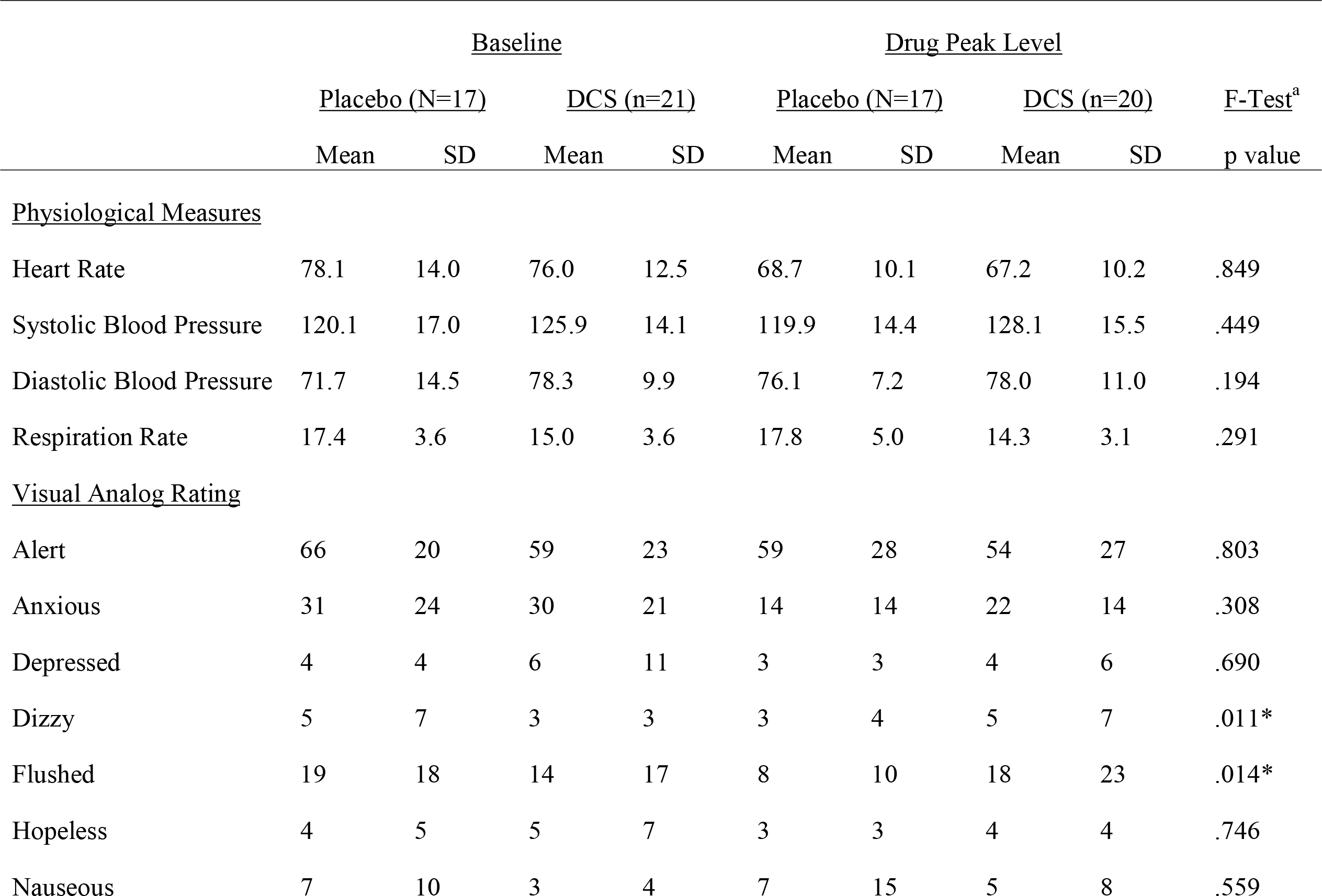
Heart rate, blood pressure and visual analogue scale ratings in the two groups before drug intake and at drug peak-level.

**Table 2.**
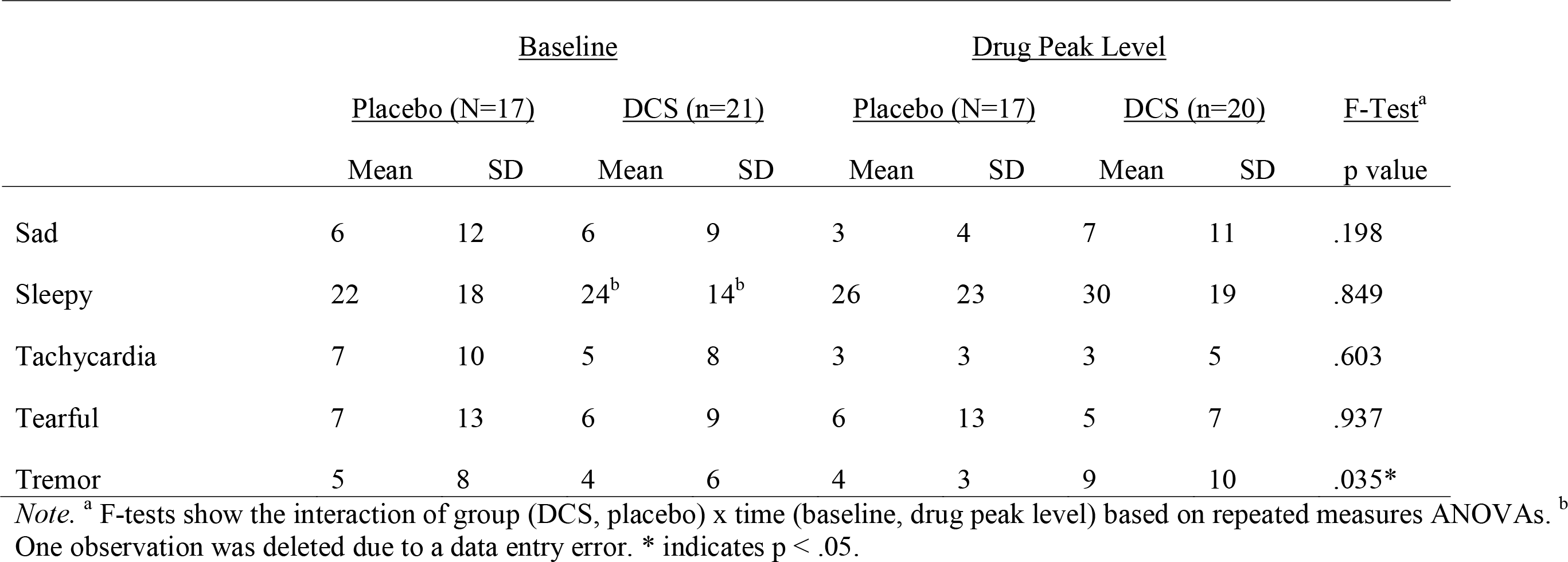

### 3.2. Effects of CBT and DCS Augmentation on Spider Fear

Linear mixed-effects modelling was used to test the improvements in fear anxiety following CBT (across groups) and DCS augmentation (between groups) (Figure 2). Subjective spider fear levels showed significant improvements from baseline to post-treatment and FU both on the SAS (*χ*^2^(3)=92.17, p<.001) and the FSQ (*χ*^2^(3)=122.80, p<.001). Significant behavioural reduction of spider fear from baseline to FU was also apparent as indicated by the BAT showing faster approach of the spider carapace following treatment (*χ*^2^(2)=31.72, p<.001). To test the efficacy of DCS augmentation to CBT on spider fear, identified linear mixed-effects regression models were used with the addition of group and group*time effects. Likelihood-ratio testing of the model with group*time effects versus the same model without the interaction effect revealed no significant effects for spider fear outcome variables SAS (*χ*^2^(3)=1.54, p=.67), FSQ (*χ*^2^(3)=0.68, p=.88), or the BAT (*χ*^2^(3)=4.47, p=.21). In sum, there was longitudinal evidence for effects of CBT, but not of DCS augmentation, on subjective and behavioural spider fear measures.

**Figure 2.**
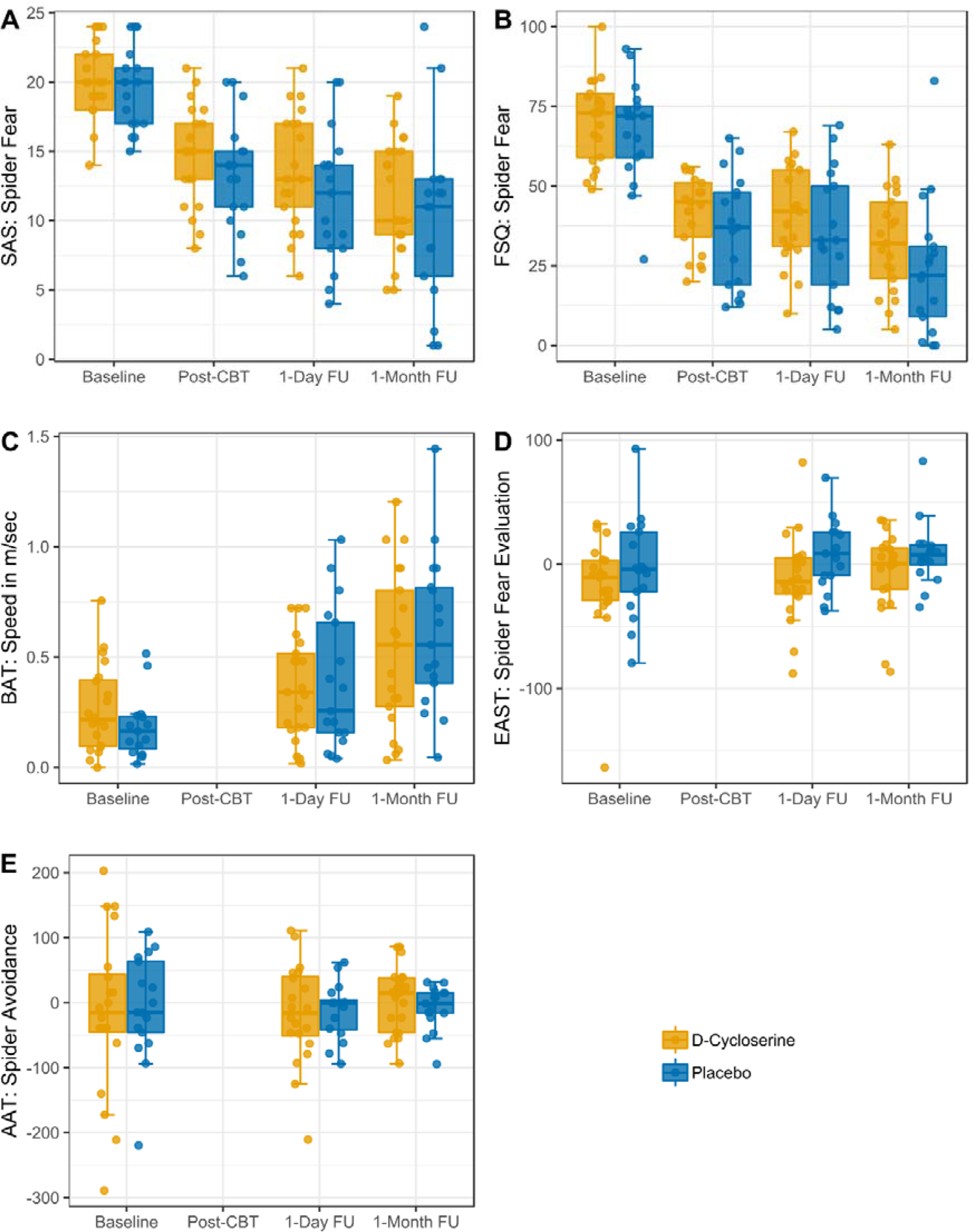
Subjective, Behavioural, and Cognitive Bias Outcome by Time and Group.

### 3.3. Effects of CBT and DCS Augmentation on Cognitive Bias Measures

Before analyses, error percentages were checked for cognitive bias measures. For the EAST, errors were generally low (averaging <1%) and only one participant had noticeable 17% errors at the baseline session. This was not deemed sufficient for exclusion from analyses, however, as only correct trials were included in analyses. For the AAT, one participant had more than 90% errors at the first session, so data were excluded from analyses. Apart from this outlier, errors were also low on the AAT (averaging ~1%).

Using the same sequential linear mixed-effects modelling approach as in 3.2, we tested whether there were significant changes in cognitive bias measures following CBT. However, neither implicit fear evaluation, as indexed by the EAST, nor avoidance, as indexed by the AAT, showed significant changes across groups (EAST: *χ*^2^(2)=3.31, p=.19; AAT: *χ*^2^(2)=0.87, p=.65). When including group and group*time effects, DCS had no significant augmentation effect on implicit fear evaluation, as indicated by the EAST (*χ*^2^(2)=0.17, p=.92), or on avoidance, as indicated by the AAT (*χ*^2^(2)=0.39, p=.82).

### 3.4. Prediction of Spider Fear at 1-month Follow-up by Cognitive Bias Levels

We explored whether cognitive bias measures at 1-day FU were related to spider fear indices at 1-month FU. Table 3 highlights a correlation matrix, which showed significant small associations (i) of implicit fear evaluation (EAST) with subjective (SAS & FSQ) and behavioural (BAT) spider fear, and (ii) of avoidance tendencies with spider fear on SAS and BAT (all Pearson’s r>.30, p<.05), but not on the FSQ. All correlations were in the predicted direction, that is, greater cognitive biases towards threat were correlated with greater spider fear (i.e., greater self-reported fear and slower behavioural approach). We next ran multiple linear regression analyses adjusted for subjective and behavioural spider fear indices at 1-day FU, to further explore whether performance on cognitive bias measures at 1-day FU predicted improvements in spider fear at 1-month FU. Results are depicted in Table 4 and showed that neither EAST nor AAT significantly predicted subjective and behavioural spider fear levels at 1-month FU after adjustment for spider fear indices.

**Table 3.**
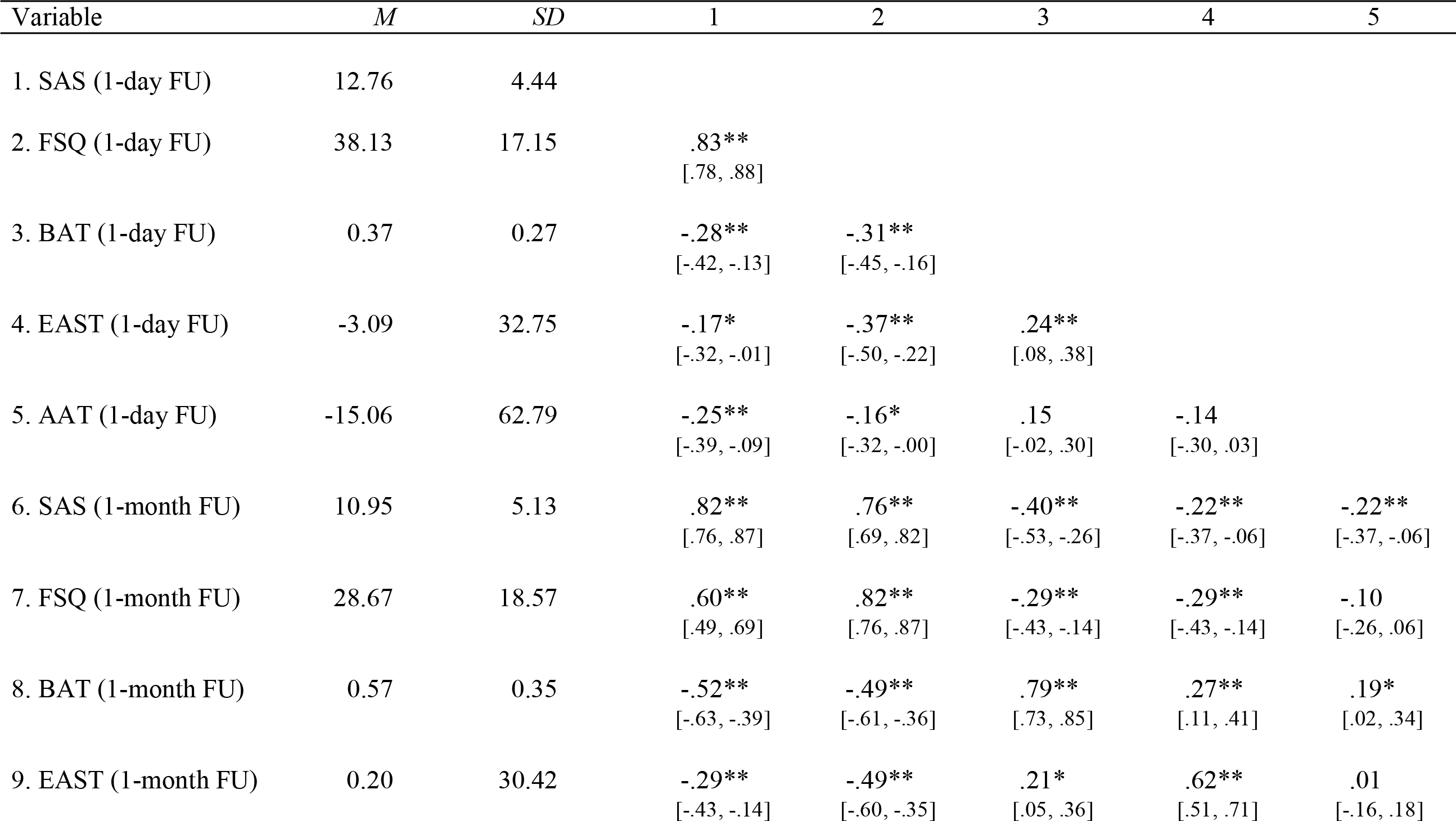
Means, standard deviations, and correlations with confidence intervals of spider fear measures at 1-day with 1-month follow-up.

**Table 3.**
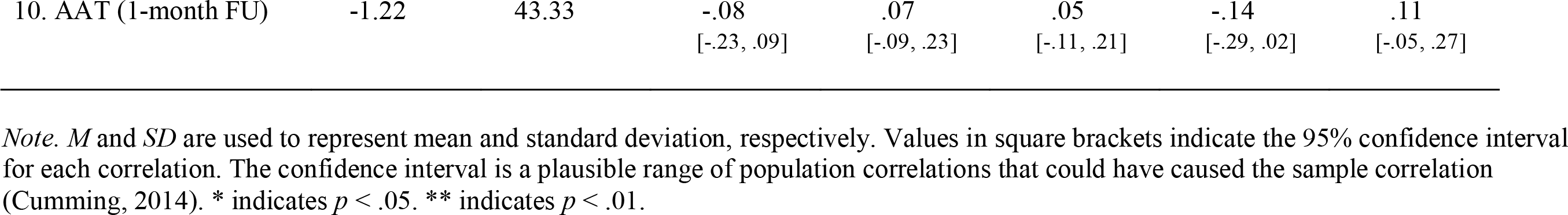

**Table 4.** Adjusted linear regression of spider fear at 1-month follow-up on cognitive bias at 1-day follow-up.

|  | 1-Month Follow-up Spider Fear Outcome |  |  |  |  |  |
| --- | --- | --- | --- | --- | --- | --- |
|  | SAS |  | FSQ |  | BAT |  |
| | $\beta$ (SE) | p value | $\beta$ (SE) | p value | $\beta$ (SE) | p value |
| Intercept | -0.813 (1.722) | 0.640 | -7.107 (5.159) | 0.178 | 0.460 (0.093) | <0.001* |
| AAT | -0.003 (0.009) | 0.697 | 0.013 (0.032) | 0.687 | 0.001 (0.001) | 0.237 |
| EAST | -0.015 (0.016) | 0.366 | 0.022 (0.064) | 0.730 | 0.003 (0.002) | 0.082 |
| Spider Fear Covariate<sup>a</sup> | 0.915 (0.129) | <0.001* | 0.936 (0.127) | <0.001* | 0.670 (0.321) | 0.046* |

*Note.* <sup>a</sup> Spider fear covariate was the respective spider fear outcome at 1-day follow-up. \* $p < .05$ .

## 4. Discussion

This randomised, controlled study for the first time investigated whether implicit fear bias is altered in response to one-session computerised CBT for spider fear and augmentation with DCS. Computerised CBT was combined with DCS and concurrent multimodal spider fear assessment from subjective, behavioural, and cognitive sides. Longitudinally, we could replicate effectiveness of computerised, one-session CBT for reducing subjective and behavioural measures of spider fear immediately following one-session treatment and further reduced until 1-month FU (Müller et al., 2011). Cognitive biases of avoidance and implicit fear evaluation were not affected by short computerised CBT treatment, however, which contrasts earlier work where a longer in-vivo setting was used (Reinecke et al., 2012). Results from exploratory analyses further showed that these cognitive biases at 1-day FU did not predict subjective and behavioural spider fear at 1-month FU when adjusted for respective measures at 1-day FU. Finally, there was no evidence for DCS augmenting any clinical effects throughout spider fear outcome measures. These findings thus add to the existing literature on CBT and DCS augmentation by assessing their respective impact on multimodal spider fear, including cognitive bias measures.

In earlier work, both implicit fear evaluation and avoidance tendencies improved following short CBT (versus waiting control) in spider fearful individuals (Reinecke et al., 2012), so interpretation requires careful comparisons of similarities and differences. In terms of similarities, spider fear was relatively similar across studies in terms of subjective (AAT & EAST) and behavioural (BAT) fear levels as well as the number of individuals qualifying for diagnosis of spider phobia (here: 20/38; previous study: 15/29). Cognitive biases, however, were more pronounced before treatment in the earlier study both on the EAST and the AAT. Additionally, the CBT design differed in that the present investigation predominantly used an easy to administer computerised exposure exercise while the earlier investigation involved ~3 hours of graded therapist-led exposure with live spiders and additional therapist contact on two further occasions. Importantly, both treatment protocols involved a single treatment session only.

Another investigation of implicit fear evaluation in Generalised Anxiety Disorder (GAD) following 15 weekly CBT sessions also found evidence for improvements in cognitive bias (Reinecke, Rinck, et al., 2013). Albeit the EAST in that study was not capturing spider fear but more generalised worry (e.g., accidents or wars), it is noteworthy that bias levels were somewhat higher as well. These mentioned differences to earlier work might account for non-replication of cognitive bias measures predicting symptom improvements, which were shown in GAD and panic disorder (Reinecke, Rinck, et al., 2013; Reinecke, Waldenmaier, et al., 2013), although the latter study assessed emotion bias using the dot-probe task, which differs from avoidance and implicit fear evaluation.

Additionally, brief computerised CBT might not have the same effects on cognitive bias as previous studies with higher therapist-contact. In sum, a combination of less severe initial bias levels and a less intense, computerised treatment protocol in the present study might have prevented alteration of cognitive bias levels. However, it is also possible that change in implicit fear bias is not relevant to symptom change in response to brief computerised CBT for spider fear. Future studies should test this using more standard CBT protocols. Such replication would be especially relevant (i) as both attentional bias and implicit fear bias predicted treatment changes in GAD and panic disorder (Reinecke, Rinck, et al., 2013; Reinecke, Waldenmaier, et al., 2013) and (ii) since our results are unlikely to result from invalid cognitive bias measures as all correlations of cognitive bias measures at 1-day FU with subjective and behavioural fear indices at 1-month FU were in the predicted direction (see again Table 3).

Lack of efficacy of DCS augmentation in the present study might have been caused by use of a subclinical sample, or by the partly computerised application of CBT. As only half of the sample qualified for a diagnosis, room for improvement on subjective and behavioural spider fear indices might have been small and sufficiently targeted by the effective one-session CBT protocol. Lastly, it is not clear whether the computerised exposure element of our treatment protocol induced sufficient new learning of non-fearful associations for DCS to act upon. This latter interpretation requires future work, however, as the present investigation constitutes the first partly computerised CBT application augmented by DCS.

While strengths of this study include randomised design for DCS investigation, multimodal assessment of spider fear, and innovative, partly computerised CBT protocol, several limitations are worth mentioning. First, replication of CBT effectiveness is restricted by lack of a CBT control group as this study was set up to primarily investigate DCS augmentation. Second, the small sample precludes firm conclusion about DCS effectiveness following the partly computerised treatment protocol in spider-fearful individuals. Third, the sample might have been unrepresentative of the general population in terms of education and intelligence (both high), largely Caucasian ethnicity, and young age. Future work is required that addresses these shortcomings in a larger and more representative cohort.

To conclude, one-session, computerised CBT shows promise for treatment of spider fear as it reduced subjective and behavioural spider fear indices with improvements until 1-month FU. However, there was no evidence for improvements in avoidance and implicit fear evaluations of spider stimuli, for unique predictive value of cognitive bias measures for subjective and behavioural treatment improvements, or for augmentation effect of DCS.

## Supporting information

Supplementary Script for Analysis and Results

## Acknowledgements

This research was funded by a grant from the Oxfordshire Health Services Research Committee awarded to AR. AR is funded by a fellowship from MQ: Transforming Mental Health. NK is supported by the International Max Planck Research School for Translational Psychiatry (IMPRS-TP).

