## Supplementary Script for Analysis and Results for "D-Cycloserine as Adjunct to Brief Computerised CBT for Spider Fear: Effects on Fear, Behaviour, and Cognitive Biases"

*Supplementary Analysis Script and Results*

*2019-04-07*

**References** **99**

### Preparation

#### Loading Packages

Relevant R packages are loaded:

```
library("tidyverse")
```

```
## -- Attaching packages ------------------------------------------------------- tidyverse 1.2.1 --
```

```
## v ggplot2 3.1.0     v purrr   0.3.0
## v tibble  2.0.1     v dplyr   0.7.8
## v tidyr   0.8.2     v stringr 1.3.1
## v readr   1.3.1     v forcats 0.3.0
```

```
## -- Conflicts ---------------------------------------------------------- tidyverse_conflicts() --
## x dplyr::filter() masks stats::filter()
## x dplyr::lag()    masks stats::lag()
```

```
library("lme4")
```

```
## Loading required package: Matrix
```

```
##
## Attaching package: 'Matrix'
```

```
## The following object is masked from 'package:tidyr':
##
##     expand
```

```
library("influence.ME")
```

```
##
## Attaching package: 'influence.ME'
## The following object is masked from 'package:stats':
##
##     influence
```

```
library("apaTables")
```

#### Loading Data

Data is loaded into R:

```
load("./Data/SpiderAnxietyData.RData")
```

#### Coding

##### Time

Dummy variables are created to index study time points:

```
dat$post <- ifelse(dat$Time == 2, 1, 0)
dat$fu1 <- ifelse(dat$Time == 3, 1, 0)
dat$fu2 <- ifelse(dat$Time == 4, 1, 0)
dat$fu <- ifelse(dat$Time >= 3, 1, 0)
dat$postfu <- ifelse(dat$Time >= 2, 1, 0)
dat$Timefac <- factor(dat$Time)
dat$Sessionfac <- factor(dat$Session)
```

Centred versions of the time variables are also created:

```
dat$Timecen <- -3
dat[dat$Time == 2,]$Timecen <- -1
dat[dat$Time == 3,]$Timecen <- 1
dat[dat$Time == 4,]$Timecen <- 3
dat$Sessioncen <- dat$Session - 2
```

##### Group

A centred Group variable is created:

```
dat$Groupcen <- ifelse(dat$Group == 0, -1, 1)
```

##### Subject ID

For analysis purposes, the subject variable will be recoded from class *integer* to class *factor*:

```
dat$Subject <- factor(dat$Subject)
```

##### Outlier Removal (high error rate)

One participant had errors >90% at pre-treatment session in the AAT, so that a missing value will be introduced here for analyses:

```r
dat[dat$Subject == "843" & dat$Time == 1, "AAT"] <- NA
dat[dat$Subject == "843", "AAT_1"] <- NA
```

### Replication: CBT effect

#### SAS

##### Testing the Random Model part

The linear mixed-effects model is created with spider anxiety as outcome and fixed effects of time as independent predictor. Random effect is a random intercept by subject.

```r
model <- lmer(SAS ~ Timefac + (1 | Subject),
              data = dat, REML = FALSE)
```

Next, it is tested whether the model requires a random slope for Time (centred, linear variable) by testing the initial model against a more complex model.

```r
model2 <- lmer(SAS ~ Timefac + (1 + Timecen | Subject),
               data = dat, REML = FALSE)
```

This is the result, which shows the random slope is significant, so will be retained in the model.

```r
anova(model, model2)
```

```
## Data: dat
## Models:
## model: SAS ~ Timefac + (1 | Subject)
## model2: SAS ~ Timefac + (1 + Timecen | Subject)
##        Df    AIC    BIC logLik deviance  Chisq Chi Df Pr(>Chisq)
## model   6 807.21 825.35 -397.6   795.21
## model2  8 775.00 799.19 -379.5   759.00 36.211      2   1.37e-08 ***
## ---
## Signif. codes:  0 '***' 0.001 '**' 0.01 '*' 0.05 '.' 0.1 ' ' 1
```

##### Testing the effect of interest

The base model is now tested against the same models without the effects of time:

```r
model_null <- lmer(SAS ~  (1 + Timecen | Subject),
              data = dat, REML = FALSE)
```

```
## Warning in checkConv(attr(opt, "derivs"), opt$par, ctrl =
## control$checkConv, : Model failed to converge with max|grad| = 0.00450667
## (tol = 0.002, component 1)
```

These are the results, which support rejection of null hypothesis:

```r
anova(model2, model_null)
```

```
## Data: dat
## Models:
## model_null: SAS ~ (1 + Timecen | Subject)
## model2: SAS ~ Timefac + (1 + Timecen | Subject)
##            Df    AIC    BIC  logLik deviance  Chisq Chi Df Pr(>Chisq)
```

```
## model_null  5 861.17 876.29 -425.58   851.17
## model2      8 775.00 799.19 -379.50   759.00 92.172      3  < 2.2e-16 ***
## ---
## Signif. codes:  0 '***' 0.001 '**' 0.01 '*' 0.05 '.' 0.1 ' ' 1
```

This is the final model (re-estimated with REML=TRUE):

```
model <- lmer(SAS ~ Timefac + (1 + Timecen | Subject),
              data = dat, REML = TRUE)
```

```
## Warning in checkConv(attr(opt, "derivs"), opt$par, ctrl =
## control$checkConv, : Model failed to converge with max|grad| = 0.00278769
## (tol = 0.002, component 1)
```

```
summary(model)
```

```
## Linear mixed model fit by REML ['lmerMod']
## Formula: SAS ~ Timefac + (1 + Timecen | Subject)
##    Data: dat
##
## REML criterion at convergence: 757.5
##
## Scaled residuals:
##      Min       1Q   Median       3Q      Max
## -2.01655 -0.50018  0.07909  0.49121  2.74180
##
## Random effects:
##  Groups   Name        Variance Std.Dev. Corr
##  Subject  (Intercept) 11.2440  3.3532
##           Timecen      0.4301  0.6558   0.78
##  Residual              4.0238  2.0060
## Number of obs: 152, groups:  Subject, 38
##
## Fixed effects:
##             Estimate Std. Error t value
## (Intercept)  19.8421     0.4815   41.21
## Timefac2     -5.7368     0.5070  -11.31
## Timefac3     -7.0789     0.6268  -11.29
## Timefac4     -8.8947     0.7869  -11.30
##
## Correlation of Fixed Effects:
##          (Intr) Timfc2 Timfc3
## Timefac2 -0.341
## Timefac3 -0.200  0.618
## Timefac4 -0.100  0.606  0.765
## convergence code: 0
## Model failed to converge with max|grad| = 0.00278769 (tol = 0.002, component 1)
```

##### Assumptions

Here are the residuals plotted against fitted values to investigate underlying heteroskedasticity and linearity assumptions:

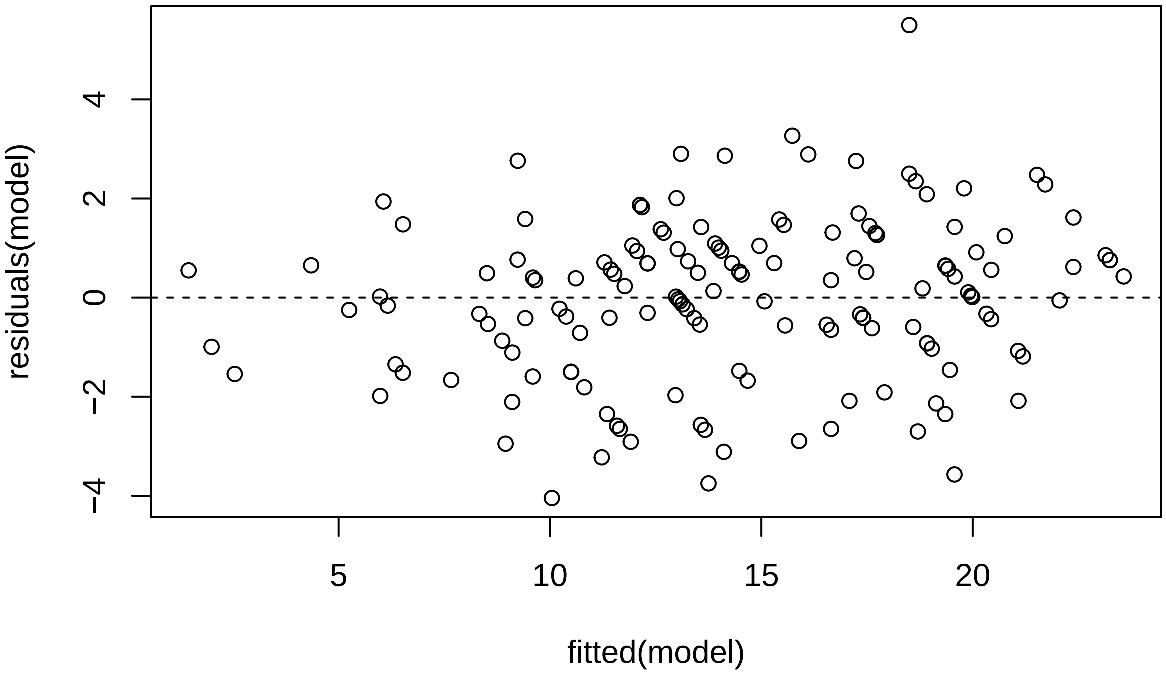

And the assumption of normality of residuals with histogram and qqplot:

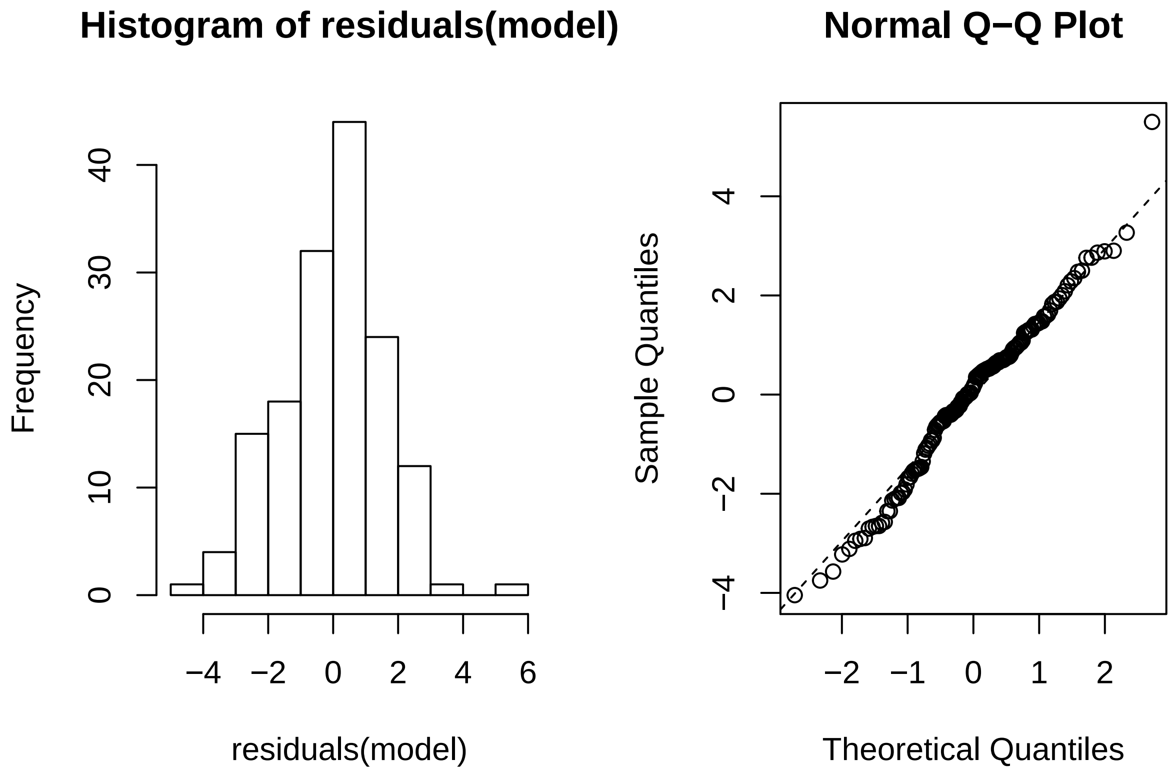

Using *Influence.ME*, outlying subjects are checked:

```
outlier <- influence(model, group = "Subject")
cks.d <- cooks.distance(outlier)
dfbetas <- dfbetas(outlier)
```

Now the Cook's distance values will be plotted. Outlier Subjects are highlighted in red:

```
plot(outlier,
     which = "cook",
     sort = TRUE,
     cutoff = 4/length(unique(dat$Subject)),
     xlab = "Cooks Distance")
```

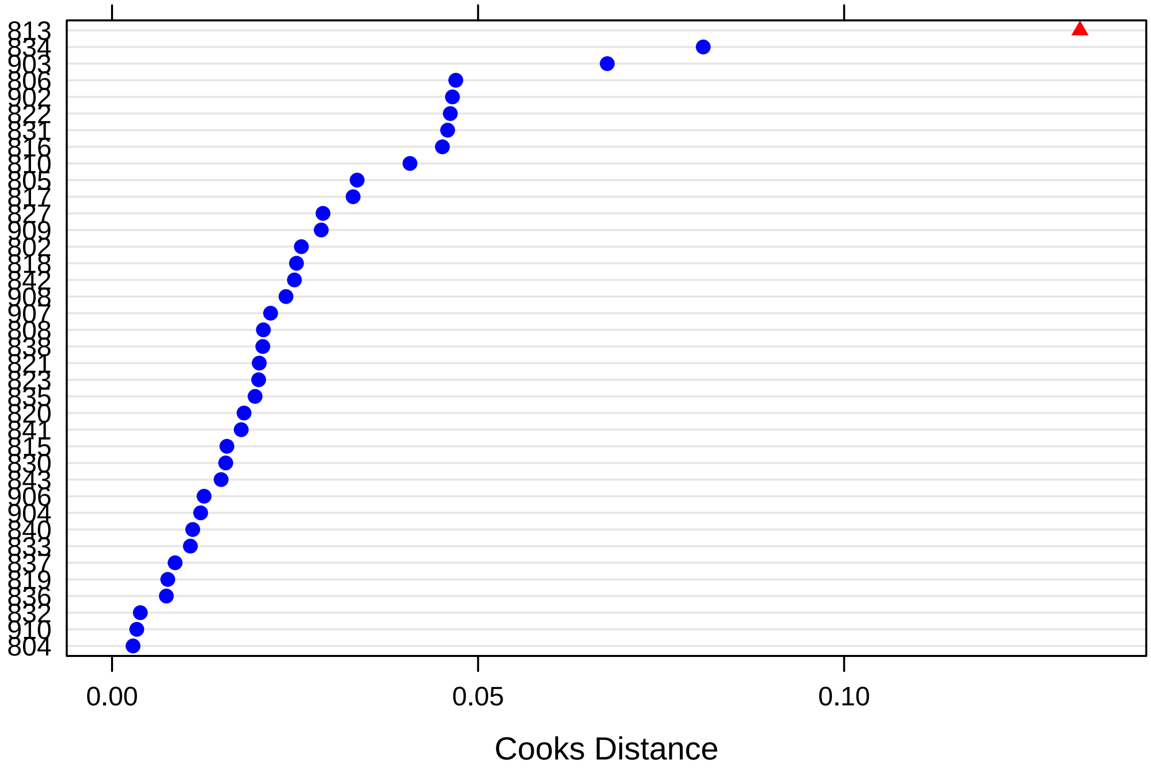

There is one outlying subject based on Cook's distance (i.e., *813*).

Finally, these are the p-values for dfbeta values, that indicate whether exclusion of a specific subject suffices to change significance values:

```
sigtest(outlier, test = -1.96)
```

```
## $Intercept
##     Altered.Teststat Altered.Sig Changed.Sig
## 802         40.89316       FALSE       FALSE
## 804         40.15010       FALSE       FALSE
## 805         40.84633       FALSE       FALSE
## 806         40.67311       FALSE       FALSE
## 808         40.15738       FALSE       FALSE
## 810         41.48042       FALSE       FALSE
## 813         40.31400       FALSE       FALSE
## 815         40.51140       FALSE       FALSE
## 816         41.82236       FALSE       FALSE
## 817         40.43779       FALSE       FALSE
## 818         40.83592       FALSE       FALSE
## 819         40.33965       FALSE       FALSE
## 820         40.20097       FALSE       FALSE
## 821         40.81998       FALSE       FALSE
## 822         42.01527       FALSE       FALSE
## 823         40.39687       FALSE       FALSE
## 827         40.91940       FALSE       FALSE
## 830         40.47993       FALSE       FALSE
```

```
## 831         40.07573       FALSE       FALSE
## 832         40.10470       FALSE       FALSE
## 833         40.27306       FALSE       FALSE
## 834         42.67482       FALSE       FALSE
## 835         41.51919       FALSE       FALSE
## 836         40.30577       FALSE       FALSE
## 837         40.37853       FALSE       FALSE
## 838         40.13570       FALSE       FALSE
## 840         40.13836       FALSE       FALSE
## 841         40.83446       FALSE       FALSE
## 842         40.13116       FALSE       FALSE
## 843         40.68620       FALSE       FALSE
## 902         40.32638       FALSE       FALSE
## 903         40.70533       FALSE       FALSE
## 904         40.51161       FALSE       FALSE
## 906         40.13117       FALSE       FALSE
## 907         40.32577       FALSE       FALSE
## 908         40.50290       FALSE       FALSE
## 909         42.23358       FALSE       FALSE
## 910         40.34559       FALSE       FALSE
##
## $Timefac2
##     Altered.Teststat Altered.Sig Changed.Sig
## 802        -11.07732        TRUE       FALSE
## 804        -11.12316        TRUE       FALSE
## 805        -10.94792        TRUE       FALSE
## 806        -11.30867        TRUE       FALSE
## 808        -11.06660        TRUE       FALSE
## 810        -11.02010        TRUE       FALSE
## 813        -11.28528        TRUE       FALSE
## 815        -11.30511        TRUE       FALSE
## 816        -11.10239        TRUE       FALSE
## 817        -11.57429        TRUE       FALSE
## 818        -10.93014        TRUE       FALSE
## 819        -11.20234        TRUE       FALSE
## 820        -11.20263        TRUE       FALSE
## 821        -11.32281        TRUE       FALSE
## 822        -11.29234        TRUE       FALSE
## 823        -11.31696        TRUE       FALSE
## 827        -11.16781        TRUE       FALSE
## 830        -11.16563        TRUE       FALSE
## 831        -10.98962        TRUE       FALSE
## 832        -10.97691        TRUE       FALSE
## 833        -11.05923        TRUE       FALSE
## 834        -11.46542        TRUE       FALSE
## 835        -11.13396        TRUE       FALSE
## 836        -11.15602        TRUE       FALSE
## 837        -10.94728        TRUE       FALSE
## 838        -11.33519        TRUE       FALSE
## 840        -11.19277        TRUE       FALSE
## 841        -10.98440        TRUE       FALSE
## 842        -11.04032        TRUE       FALSE
## 843        -11.13061        TRUE       FALSE
## 902        -11.18710        TRUE       FALSE
```

```
## 903        -11.92449        TRUE       FALSE
## 904        -11.28260        TRUE       FALSE
## 906        -11.02432        TRUE       FALSE
## 907        -11.00678        TRUE       FALSE
## 908        -10.92802        TRUE       FALSE
## 909        -11.13492        TRUE       FALSE
## 910        -11.06558        TRUE       FALSE
##
## $Timefac3
##     Altered.Teststat Altered.Sig Changed.Sig
## 802        -11.25293        TRUE       FALSE
## 804        -11.04920        TRUE       FALSE
## 805        -10.92754        TRUE       FALSE
## 806        -11.49966        TRUE       FALSE
## 808        -10.87255        TRUE       FALSE
## 810        -11.12163        TRUE       FALSE
## 813        -11.44758        TRUE       FALSE
## 815        -11.34085        TRUE       FALSE
## 816        -11.10361        TRUE       FALSE
## 817        -11.34174        TRUE       FALSE
## 818        -11.02048        TRUE       FALSE
## 819        -11.14328        TRUE       FALSE
## 820        -10.99017        TRUE       FALSE
## 821        -11.31743        TRUE       FALSE
## 822        -11.23039        TRUE       FALSE
## 823        -10.98192        TRUE       FALSE
## 827        -11.27911        TRUE       FALSE
## 830        -11.23480        TRUE       FALSE
## 831        -11.22168        TRUE       FALSE
## 832        -10.99342        TRUE       FALSE
## 833        -11.03888        TRUE       FALSE
## 834        -11.31550        TRUE       FALSE
## 835        -11.01172        TRUE       FALSE
## 836        -10.99005        TRUE       FALSE
## 837        -11.02087        TRUE       FALSE
## 838        -11.42876        TRUE       FALSE
## 840        -11.06976        TRUE       FALSE
## 841        -11.05690        TRUE       FALSE
## 842        -11.20931        TRUE       FALSE
## 843        -11.07321        TRUE       FALSE
## 902        -11.08399        TRUE       FALSE
## 903        -11.53831        TRUE       FALSE
## 904        -11.19821        TRUE       FALSE
## 906        -11.04245        TRUE       FALSE
## 907        -11.14584        TRUE       FALSE
## 908        -10.90244        TRUE       FALSE
## 909        -11.09785        TRUE       FALSE
## 910        -11.04537        TRUE       FALSE
##
## $Timefac4
##     Altered.Teststat Altered.Sig Changed.Sig
## 802        -11.36935        TRUE       FALSE
## 804        -11.03727        TRUE       FALSE
## 805        -10.95701        TRUE       FALSE
```

```
## 806         -11.80883       TRUE       FALSE
## 808         -10.99580       TRUE       FALSE
## 810         -10.95603       TRUE       FALSE
## 813         -11.63060       TRUE       FALSE
## 815         -11.32585       TRUE       FALSE
## 816         -11.24387       TRUE       FALSE
## 817         -11.14997       TRUE       FALSE
## 818         -10.99189       TRUE       FALSE
## 819         -11.20737       TRUE       FALSE
## 820         -11.12892       TRUE       FALSE
## 821         -11.24021       TRUE       FALSE
## 822         -11.05004       TRUE       FALSE
## 823         -11.04194       TRUE       FALSE
## 827         -11.42323       TRUE       FALSE
## 830         -11.17497       TRUE       FALSE
## 831         -11.50362       TRUE       FALSE
## 832         -10.94436       TRUE       FALSE
## 833         -10.96645       TRUE       FALSE
## 834         -10.97886       TRUE       FALSE
## 835         -11.00410       TRUE       FALSE
## 836         -10.99008       TRUE       FALSE
## 837         -11.00470       TRUE       FALSE
## 838         -11.63209       TRUE       FALSE
## 840         -11.19651       TRUE       FALSE
## 841         -11.00490       TRUE       FALSE
## 842         -11.14602       TRUE       FALSE
## 843         -11.13984       TRUE       FALSE
## 902         -11.33434       TRUE       FALSE
## 903         -11.11236       TRUE       FALSE
## 904         -11.20951       TRUE       FALSE
## 906         -11.03675       TRUE       FALSE
## 907         -11.13382       TRUE       FALSE
## 908         -11.00689       TRUE       FALSE
## 909         -11.04038       TRUE       FALSE
## 910         -11.00121       TRUE       FALSE
```

There is no indication that results are changed by an outlying subject.

#### FSQ

##### Testing the Random Model part

The linear mixed-effects model is created with spider anxiety as outcome and fixed effects of time as independent predictor. Random effect is a random intercept by subject.

```
model <- lmer(FSQ ~ Timefac + (1 | Subject),
              data = dat, REML = FALSE)
```

Next, it is tested whether the model requires a random slope for Time (centred, linear variable) by testing the initial model against a more complex model.

```
model2 <- lmer(FSQ ~ Timefac + (1 + Timecen | Subject),
               data = dat, REML = FALSE)
```

This is the result, which shows the random slope is significant, so will be retained in the model.

```r
anova(model, model2)
```

```
## Data: dat
## Models:
## model: FSQ ~ Timefac + (1 | Subject)
## model2: FSQ ~ Timefac + (1 + Timecen | Subject)
##        Df    AIC    BIC  logLik deviance  Chisq Chi Df Pr(>Chisq)
## model   6 1227.2 1245.4 -607.62   1215.2
## model2  8 1211.7 1235.9 -597.84   1195.7 19.566      2   5.64e-05 ***
## ---
## Signif. codes:  0 '***' 0.001 '**' 0.01 '*' 0.05 '.' 0.1 ' ' 1
```

###### Testing the effect of interest

The base model is now tested against the same models without the effects of time:

```r
model_null <- lmer(FSQ ~  (1 + Timecen | Subject),
              data = dat, REML = FALSE)
```

```
## Warning in checkConv(attr(opt, "derivs"), opt$par, ctrl =
## control$checkConv, : Model failed to converge with max|grad| = 0.00274881
## (tol = 0.002, component 1)
```

These are the results, which support rejection of null hypothesis:

```r
anova(model2, model_null)
```

```
## Data: dat
## Models:
## model_null: FSQ ~ (1 + Timecen | Subject)
## model2: FSQ ~ Timefac + (1 + Timecen | Subject)
##            Df    AIC    BIC  logLik deviance Chisq Chi Df Pr(>Chisq)
## model_null  5 1328.5 1343.6 -659.24   1318.5
## model2      8 1211.7 1235.9 -597.84   1195.7 122.8      3  < 2.2e-16 ***
## ---
## Signif. codes:  0 '***' 0.001 '**' 0.01 '*' 0.05 '.' 0.1 ' ' 1
```

This is the final model (re-estimated with REML=TRUE):

```r
model <- lmer(FSQ ~ Timefac + (1 + Timecen | Subject),
              data = dat, REML = TRUE)

summary(model)
```

```
## Linear mixed model fit by REML ['lmerMod']
## Formula: FSQ ~ Timefac + (1 + Timecen | Subject)
##    Data: dat
##
## REML criterion at convergence: 1182.7
##
## Scaled residuals:
##      Min       1Q   Median       3Q      Max
## -1.96496 -0.50693  0.01822  0.51844  2.70490
##
## Random effects:
##  Groups   Name        Variance Std.Dev. Corr
```

```
##  Subject  (Intercept) 170.056  13.041
##           Timecen       5.388   2.321   0.62
##  Residual              74.365   8.624
## Number of obs: 152, groups:  Subject, 38
##
## Fixed effects:
##             Estimate Std. Error t value
## (Intercept)   69.079      2.180   31.69
## Timefac2     -29.921      2.117  -14.13
## Timefac3     -30.947      2.486  -12.45
## Timefac4     -40.408      3.003  -13.46
##
## Correlation of Fixed Effects:
##          (Intr) Timfc2 Timfc3
## Timefac2 -0.395
## Timefac3 -0.311  0.587
## Timefac4 -0.237  0.575  0.718
```

**Assumptions**

Here are the residuals plotted against fitted values to investigate underlying heteroskedasticity and linearity assumptions:

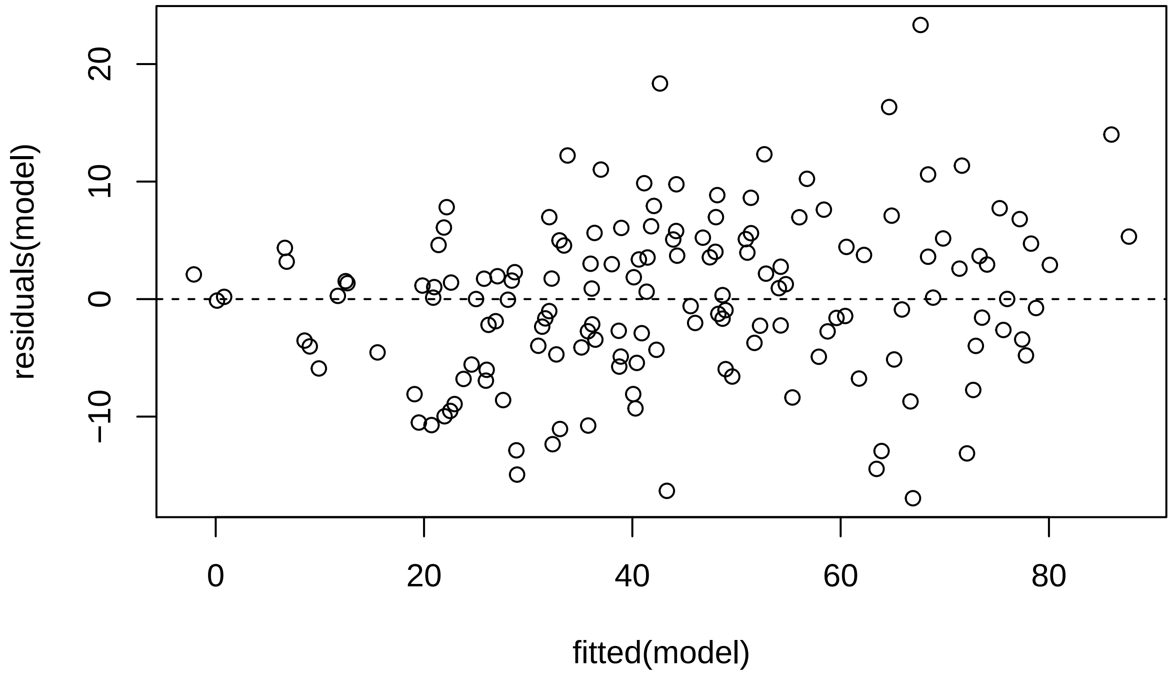

And the assumption of normality of residuals with histogram and qqplot:

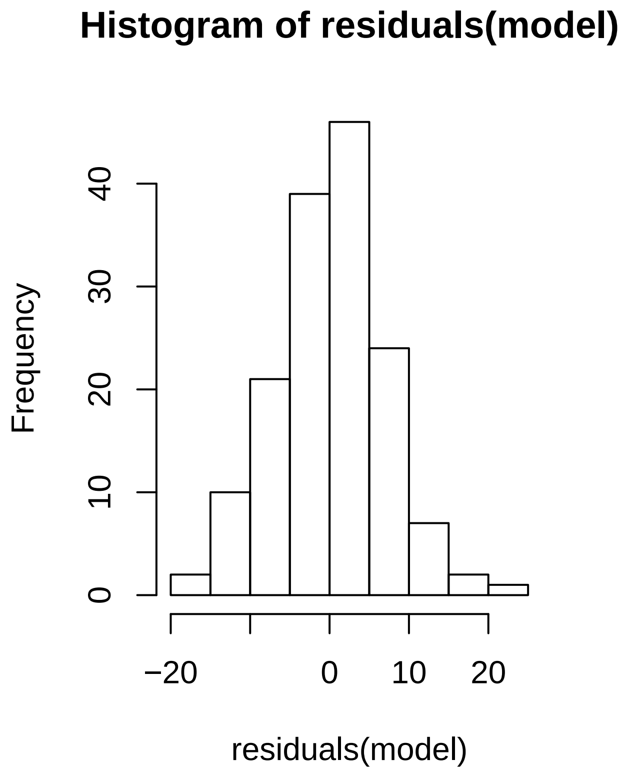

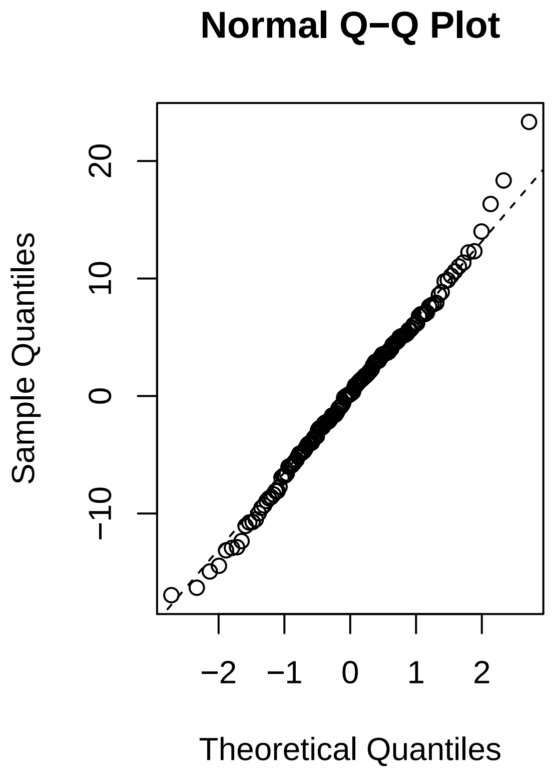

Using *Influence.ME*, outlying subjects are checked:

```
outlier <- influence(model, group = "Subject")
cks.d <- cooks.distance(outlier)
dfbetas <- dfbetas(outlier)
```

Now the Cook's distance values will be plotted. Outlier Subjects are highlighted in red:

```
plot(outlier,
     which = "cook",
     sort = TRUE,
     cutoff = 4/length(unique(dat$Subject)),
     xlab = "Cooks Distance")
```

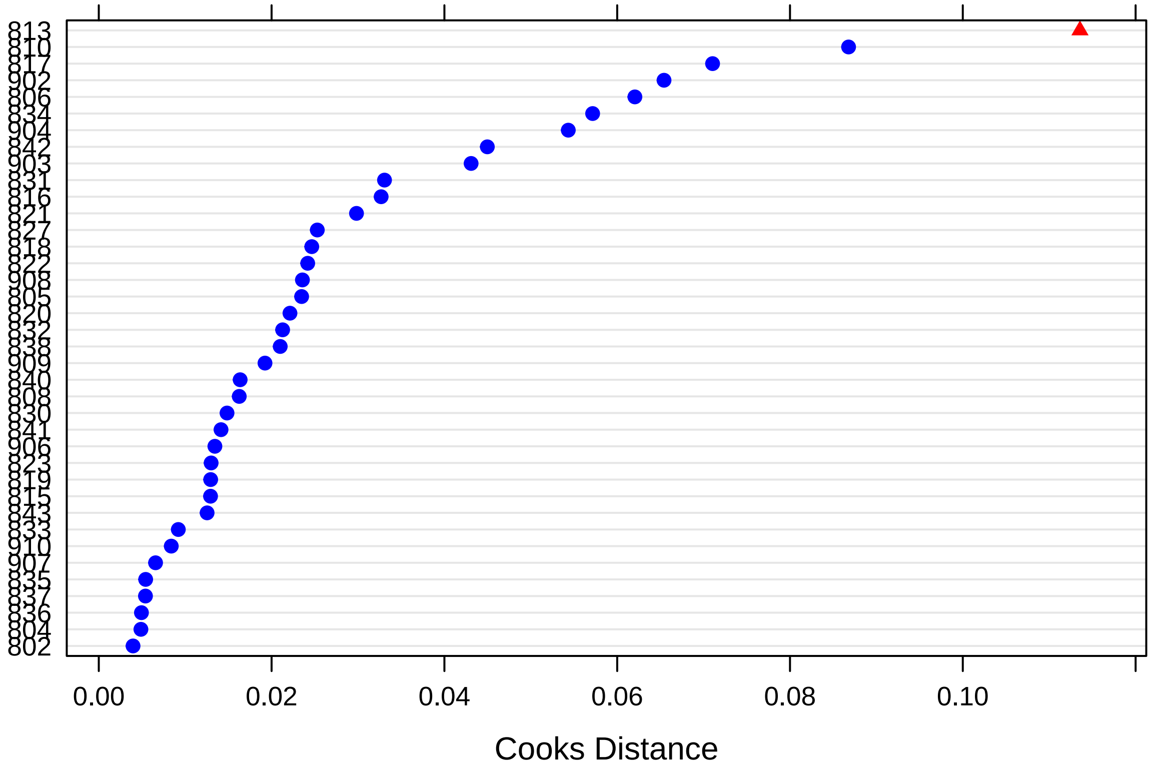

There is one outlying subject based on Cook's distance (i.e., *813*).

Finally, these are the p-values for dfbeta values, that indicate whether exclusion of a specific subject suffices to change significance values:

```
sigtest(outlier, test = -1.96)
```

```
## $Intercept
##     Altered.Teststat Altered.Sig Changed.Sig
## 802         30.84385       FALSE       FALSE
## 804         30.80640       FALSE       FALSE
## 805         30.81392       FALSE       FALSE
## 806         31.52406       FALSE       FALSE
## 808         30.85381       FALSE       FALSE
## 810         35.56010       FALSE       FALSE
## 813         30.96778       FALSE       FALSE
## 815         30.97004       FALSE       FALSE
## 816         32.55321       FALSE       FALSE
## 817         31.26173       FALSE       FALSE
## 818         31.18639       FALSE       FALSE
## 819         30.94831       FALSE       FALSE
## 820         31.21889       FALSE       FALSE
## 821         30.95046       FALSE       FALSE
## 822         30.98552       FALSE       FALSE
## 823         30.90279       FALSE       FALSE
## 827         31.11568       FALSE       FALSE
## 830         30.99877       FALSE       FALSE
```

```
## 831          31.15832       FALSE       FALSE
## 832          31.44349       FALSE       FALSE
## 833          31.24420       FALSE       FALSE
## 834          32.60219       FALSE       FALSE
## 835          31.08427       FALSE       FALSE
## 836          30.82595       FALSE       FALSE
## 837          30.85280       FALSE       FALSE
## 838          30.90242       FALSE       FALSE
## 840          30.97865       FALSE       FALSE
## 841          31.31375       FALSE       FALSE
## 842          31.72688       FALSE       FALSE
## 843          30.96254       FALSE       FALSE
## 902          30.87846       FALSE       FALSE
## 903          31.43621       FALSE       FALSE
## 904          31.39484       FALSE       FALSE
## 906          31.51035       FALSE       FALSE
## 907          30.92345       FALSE       FALSE
## 908          31.09854       FALSE       FALSE
## 909          31.08949       FALSE       FALSE
## 910          31.36275       FALSE       FALSE
##
## $Timefac2
##     Altered.Teststat Altered.Sig Changed.Sig
## 802        -13.75309        TRUE       FALSE
## 804        -13.69358        TRUE       FALSE
## 805        -13.69800        TRUE       FALSE
## 806        -14.02067        TRUE       FALSE
## 808        -13.70707        TRUE       FALSE
## 810        -14.03558        TRUE       FALSE
## 813        -14.12623        TRUE       FALSE
## 815        -13.98523        TRUE       FALSE
## 816        -13.86171        TRUE       FALSE
## 817        -14.93253        TRUE       FALSE
## 818        -13.76692        TRUE       FALSE
## 819        -13.75365        TRUE       FALSE
## 820        -13.75333        TRUE       FALSE
## 821        -14.16552        TRUE       FALSE
## 822        -14.07754        TRUE       FALSE
## 823        -13.93645        TRUE       FALSE
## 827        -13.90445        TRUE       FALSE
## 830        -13.98412        TRUE       FALSE
## 831        -13.68145        TRUE       FALSE
## 832        -13.81807        TRUE       FALSE
## 833        -13.77605        TRUE       FALSE
## 834        -13.74714        TRUE       FALSE
## 835        -13.90802        TRUE       FALSE
## 836        -13.73597        TRUE       FALSE
## 837        -13.80737        TRUE       FALSE
## 838        -13.95281        TRUE       FALSE
## 840        -13.80367        TRUE       FALSE
## 841        -13.81639        TRUE       FALSE
## 842        -14.19031        TRUE       FALSE
## 843        -13.96572        TRUE       FALSE
## 902        -13.93740        TRUE       FALSE
```

```
## 903        -14.54337        TRUE       FALSE
## 904        -14.69516        TRUE       FALSE
## 906        -13.81221        TRUE       FALSE
## 907        -13.88631        TRUE       FALSE
## 908        -13.79357        TRUE       FALSE
## 909        -14.21831        TRUE       FALSE
## 910        -13.90043        TRUE       FALSE
##
## $Timefac3
##     Altered.Teststat Altered.Sig Changed.Sig
## 802        -12.10513        TRUE       FALSE
## 804        -12.03081        TRUE       FALSE
## 805        -12.08565        TRUE       FALSE
## 806        -12.52257        TRUE       FALSE
## 808        -12.02398        TRUE       FALSE
## 810        -12.27626        TRUE       FALSE
## 813        -12.59574        TRUE       FALSE
## 815        -12.34467        TRUE       FALSE
## 816        -12.43256        TRUE       FALSE
## 817        -12.66079        TRUE       FALSE
## 818        -12.24510        TRUE       FALSE
## 819        -12.20238        TRUE       FALSE
## 820        -12.14219        TRUE       FALSE
## 821        -12.74480        TRUE       FALSE
## 822        -12.55790        TRUE       FALSE
## 823        -12.11089        TRUE       FALSE
## 827        -12.29592        TRUE       FALSE
## 830        -12.33155        TRUE       FALSE
## 831        -12.09558        TRUE       FALSE
## 832        -12.23770        TRUE       FALSE
## 833        -12.04274        TRUE       FALSE
## 834        -12.24353        TRUE       FALSE
## 835        -12.18535        TRUE       FALSE
## 836        -12.03040        TRUE       FALSE
## 837        -12.03930        TRUE       FALSE
## 838        -12.42666        TRUE       FALSE
## 840        -12.06109        TRUE       FALSE
## 841        -12.17403        TRUE       FALSE
## 842        -13.02735        TRUE       FALSE
## 843        -12.14070        TRUE       FALSE
## 902        -12.17656        TRUE       FALSE
## 903        -12.51974        TRUE       FALSE
## 904        -12.87072        TRUE       FALSE
## 906        -12.14024        TRUE       FALSE
## 907        -12.22188        TRUE       FALSE
## 908        -12.00932        TRUE       FALSE
## 909        -12.27835        TRUE       FALSE
## 910        -12.20580        TRUE       FALSE
##
## $Timefac4
##     Altered.Teststat Altered.Sig Changed.Sig
## 802        -13.14336        TRUE       FALSE
## 804        -13.06242        TRUE       FALSE
## 805        -13.05342        TRUE       FALSE
```

```
## 806        -13.84514       TRUE       FALSE
## 808        -13.11939       TRUE       FALSE
## 810        -13.27473       TRUE       FALSE
## 813        -13.90557       TRUE       FALSE
## 815        -13.34237       TRUE       FALSE
## 816        -13.52080       TRUE       FALSE
## 817        -13.16325       TRUE       FALSE
## 818        -13.19761       TRUE       FALSE
## 819        -13.24486       TRUE       FALSE
## 820        -13.14542       TRUE       FALSE
## 821        -13.68032       TRUE       FALSE
## 822        -13.46537       TRUE       FALSE
## 823        -13.21820       TRUE       FALSE
## 827        -13.24512       TRUE       FALSE
## 830        -13.13347       TRUE       FALSE
## 831        -13.17415       TRUE       FALSE
## 832        -13.02743       TRUE       FALSE
## 833        -13.07952       TRUE       FALSE
## 834        -13.12673       TRUE       FALSE
## 835        -13.20606       TRUE       FALSE
## 836        -13.07895       TRUE       FALSE
## 837        -13.08774       TRUE       FALSE
## 838        -13.63606       TRUE       FALSE
## 840        -13.14550       TRUE       FALSE
## 841        -13.06441       TRUE       FALSE
## 842        -14.42847       TRUE       FALSE
## 843        -13.19872       TRUE       FALSE
## 902        -13.27879       TRUE       FALSE
## 903        -13.20835       TRUE       FALSE
## 904        -13.63487       TRUE       FALSE
## 906        -13.05826       TRUE       FALSE
## 907        -13.09454       TRUE       FALSE
## 908        -13.13080       TRUE       FALSE
## 909        -13.17626       TRUE       FALSE
## 910        -13.21051       TRUE       FALSE
```

There is no indication that results are changed by an outlying subject.

#### BAT

##### Testing the Random Model part

The linear mixed-effects model is created with spider anxiety as outcome and fixed effects of time as independent predictor. Random effect is a random intercept by subject.

```
model <- lmer(BAT_speed ~ Sessionfac + (1 | Subject),
              data = dat, REML = FALSE)
```

```
## Warning in checkConv(attr(opt, "derivs"), opt$par, ctrl =
## control$checkConv, : Model failed to converge with max|grad| = 0.00267083
## (tol = 0.002, component 1)
```

Next, it is tested whether the model requires a random slope for Time (centred, linear variable) by testing the initial model against a more complex model.

```r
model2 <- lmer(BAT_speed ~ Sessionfac + (1 + Sessioncen | Subject),
               data = dat, REML = FALSE)
```

This is the result, which shows the random slope is significant, so will be retained in the model.

```r
anova(model, model2)
```

```
## Data: dat
## Models:
## model: BAT_speed ~ Sessionfac + (1 | Subject)
## model2: BAT_speed ~ Sessionfac + (1 + Sessioncen | Subject)
##        Df      AIC     BIC  logLik deviance  Chisq Chi Df Pr(>Chisq)
## model   5   1.9109 15.4585  4.0446   -8.089
## model2  7 -23.4514 -4.4847 18.7257  -37.451 29.362      2  4.208e-07 ***
## ---
## Signif. codes:  0 '***' 0.001 '**' 0.01 '*' 0.05 '.' 0.1 ' ' 1
```

##### Testing the effect of interest

The base model is now tested against the same models without the effects of time:

```r
model_null <- lmer(BAT_speed ~  (1 + Sessioncen | Subject),
               data = dat, REML = FALSE)
```

These are the results, which support rejection of null hypothesis:

```r
anova(model2, model_null)
```

```
## Data: dat
## Models:
## model_null: BAT_speed ~ (1 + Sessioncen | Subject)
## model2: BAT_speed ~ Sessionfac + (1 + Sessioncen | Subject)
##            Df      AIC     BIC  logLik deviance  Chisq Chi Df Pr(>Chisq)
## model_null  5   4.2671 17.8148  2.8664   -5.733
## model2      7 -23.4514 -4.4847 18.7257  -37.451 31.718      2  1.295e-07
##
## model_null
## model2     ***
## ---
## Signif. codes:  0 '***' 0.001 '**' 0.01 '*' 0.05 '.' 0.1 ' ' 1
```

This is the final model (re-estimated with REML=TRUE):

```r
model <- lmer(BAT_speed ~ Sessionfac + (1 + Sessioncen | Subject),
               data = dat, REML = TRUE)

summary(model)
```

```
## Linear mixed model fit by REML ['lmerMod']
## Formula: BAT_speed ~ Sessionfac + (1 + Sessioncen | Subject)
##    Data: dat
##
## REML criterion at convergence: -22.6
##
## Scaled residuals:
##     Min      1Q  Median      3Q     Max
## -2.3553 -0.4205 -0.0833  0.3858  2.4937
```

```
##
## Random effects:
##  Groups   Name        Variance Std.Dev. Corr
##  Subject  (Intercept) 0.05003  0.2237
##           Sessioncen  0.01668  0.1291   0.79
##  Residual             0.01541  0.1241
## Number of obs: 111, groups:  Subject, 38
##
## Fixed effects:
##             Estimate Std. Error t value
## (Intercept)  0.22010    0.03180   6.921
## Sessionfac2  0.15172    0.03614   4.198
## Sessionfac3  0.34909    0.05121   6.817
##
## Correlation of Fixed Effects:
##             (Intr) Sssnf2
## Sessionfac2 -0.259
## Sessionfac3 -0.082  0.724
```

**Assumptions**

Here are the residuals plotted against fitted values to investigate underlying heteroskedasticity and linearity assumptions:

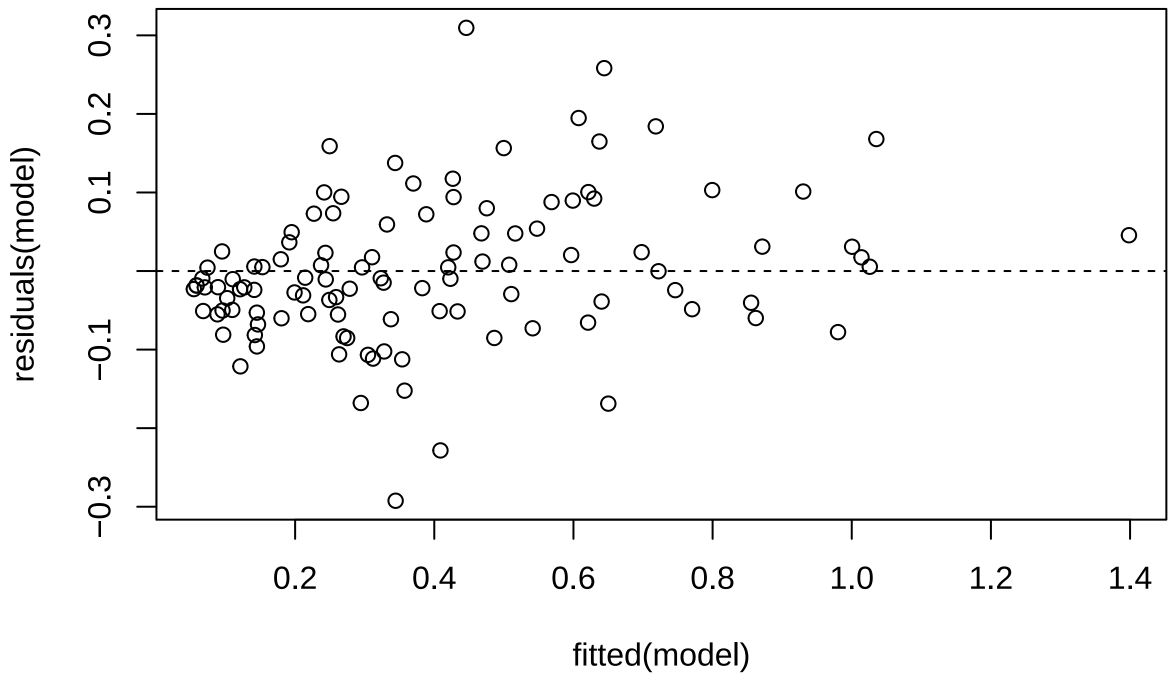

And the assumption of normality of residuals with histogram and qqplot:

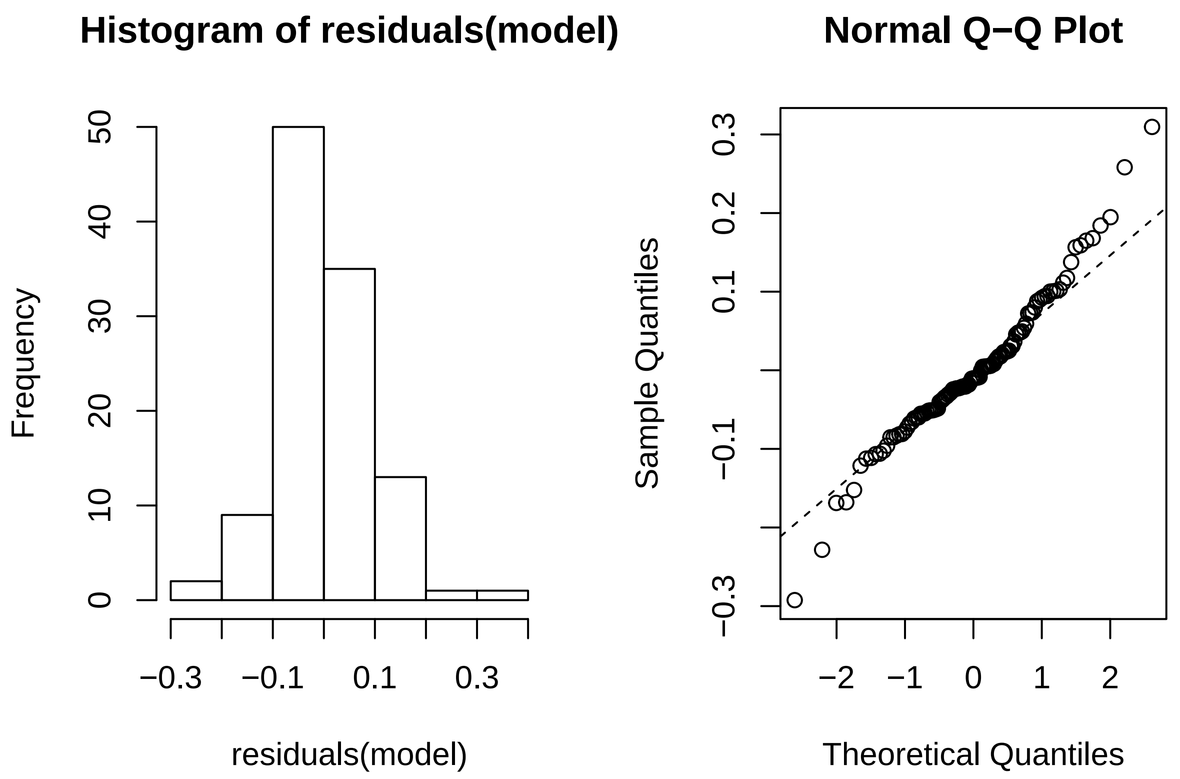

Using *Influence.ME*, outlying subjects are checked:

```
outlier <- influence(model, group = "Subject")
cks.d <- cooks.distance(outlier)
dfbetas <- dfbetas(outlier)
```

Now the Cook's distance values will be plotted. Outlier Subjects are highlighted in red:

```
plot(outlier,
     which = "cook",
     sort = TRUE,
     cutoff = 4/length(unique(dat$Subject)),
     xlab = "Cooks Distance")
```

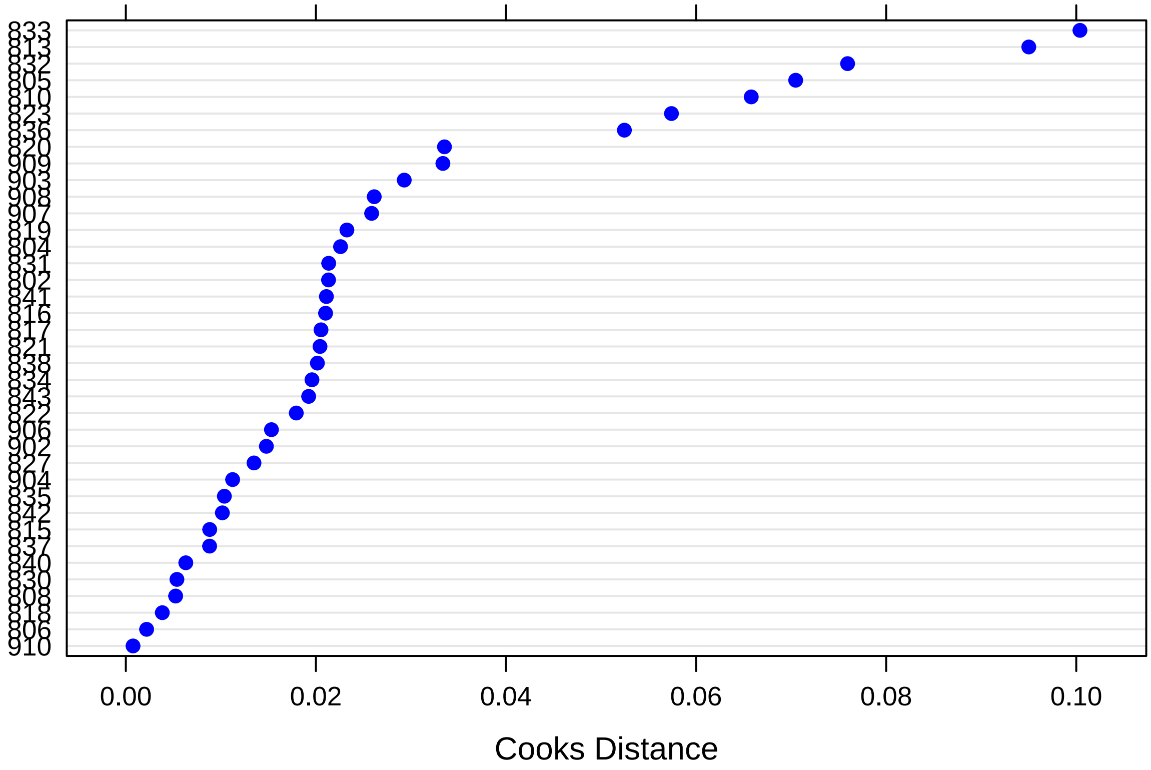

There are no outlying subjects.

Finally, these are the p-values for dfbeta values, that indicate whether exclusion of a specific subject suffices to change significance values:

```
sigtest(outlier, test = -1.96)
```

```
## $Intercept
##     Altered.Teststat Altered.Sig Changed.Sig
## 802         6.886798       FALSE       FALSE
## 804         6.761692       FALSE       FALSE
## 805         7.015242       FALSE       FALSE
## 806         6.745771       FALSE       FALSE
## 808         6.862161       FALSE       FALSE
## 810         6.738921       FALSE       FALSE
## 813         7.110712       FALSE       FALSE
## 815         6.882900       FALSE       FALSE
## 816         6.821260       FALSE       FALSE
## 817         6.929413       FALSE       FALSE
## 818         6.730066       FALSE       FALSE
## 819         6.688991       FALSE       FALSE
## 820         6.576846       FALSE       FALSE
## 821         6.726406       FALSE       FALSE
## 822         6.895703       FALSE       FALSE
## 823         6.607007       FALSE       FALSE
## 827         7.025182       FALSE       FALSE
## 830         6.801575       FALSE       FALSE
```

```
## 831          6.708176       FALSE       FALSE
## 832          6.753890       FALSE       FALSE
## 833          7.081913       FALSE       FALSE
## 834          7.000922       FALSE       FALSE
## 835          6.913964       FALSE       FALSE
## 836          6.832537       FALSE       FALSE
## 837          6.670562       FALSE       FALSE
## 838          6.682735       FALSE       FALSE
## 840          6.878763       FALSE       FALSE
## 841          7.009372       FALSE       FALSE
## 842          6.699951       FALSE       FALSE
## 843          7.025679       FALSE       FALSE
## 902          6.925004       FALSE       FALSE
## 903          6.766453       FALSE       FALSE
## 904          6.725812       FALSE       FALSE
## 906          6.774791       FALSE       FALSE
## 907          7.116785       FALSE       FALSE
## 908          6.705438       FALSE       FALSE
## 909          6.822290       FALSE       FALSE
## 910          6.741741       FALSE       FALSE
##
## $Sessionfac2
##     Altered.Teststat Altered.Sig Changed.Sig
## 802          4.270397       FALSE       FALSE
## 804          3.927487       FALSE       FALSE
## 805          4.319122       FALSE       FALSE
## 806          4.157059       FALSE       FALSE
## 808          4.125732       FALSE       FALSE
## 810          3.920576       FALSE       FALSE
## 813          4.557077       FALSE       FALSE
## 815          4.184623       FALSE       FALSE
## 816          4.073807       FALSE       FALSE
## 817          4.241678       FALSE       FALSE
## 818          3.990118       FALSE       FALSE
## 819          4.250324       FALSE       FALSE
## 820          4.466483       FALSE       FALSE
## 821          4.028644       FALSE       FALSE
## 822          4.194514       FALSE       FALSE
## 823          4.293822       FALSE       FALSE
## 827          4.283436       FALSE       FALSE
## 830          4.180578       FALSE       FALSE
## 831          4.294121       FALSE       FALSE
## 832          3.908601       FALSE       FALSE
## 833          4.505335       FALSE       FALSE
## 834          4.216583       FALSE       FALSE
## 835          4.133971       FALSE       FALSE
## 836          3.860757       FALSE       FALSE
## 837          4.069959       FALSE       FALSE
## 838          4.214859       FALSE       FALSE
## 840          4.137805       FALSE       FALSE
## 841          3.973167       FALSE       FALSE
## 842          4.217873       FALSE       FALSE
## 843          4.299462       FALSE       FALSE
## 902          3.992261       FALSE       FALSE
```

```
## 903         3.891787       FALSE       FALSE
## 904         4.050656       FALSE       FALSE
## 906         3.928314       FALSE       FALSE
## 907         4.088061       FALSE       FALSE
## 908         4.050962       FALSE       FALSE
## 909         4.059226       FALSE       FALSE
## 910         4.081398       FALSE       FALSE
##
## $Sessionfac3
##     Altered.Teststat Altered.Sig Changed.Sig
## 802         6.988305       FALSE       FALSE
## 804         6.563080       FALSE       FALSE
## 805         6.612730       FALSE       FALSE
## 806         6.715913       FALSE       FALSE
## 808         6.635135       FALSE       FALSE
## 810         6.558199       FALSE       FALSE
## 813         6.633309       FALSE       FALSE
## 815         6.724097       FALSE       FALSE
## 816         6.578554       FALSE       FALSE
## 817         6.938194       FALSE       FALSE
## 818         6.573174       FALSE       FALSE
## 819         7.050800       FALSE       FALSE
## 820         6.939198       FALSE       FALSE
## 821         6.627954       FALSE       FALSE
## 822         6.915572       FALSE       FALSE
## 823         6.644768       FALSE       FALSE
## 827         6.819596       FALSE       FALSE
## 830         6.757769       FALSE       FALSE
## 831         6.604616       FALSE       FALSE
## 832         6.639147       FALSE       FALSE
## 833         7.282934       FALSE       FALSE
## 834         6.835261       FALSE       FALSE
## 835         6.769522       FALSE       FALSE
## 836         6.645878       FALSE       FALSE
## 837         6.642843       FALSE       FALSE
## 838         6.814291       FALSE       FALSE
## 840         6.710984       FALSE       FALSE
## 841         6.615637       FALSE       FALSE
## 842         6.903763       FALSE       FALSE
## 843         7.004248       FALSE       FALSE
## 902         6.631154       FALSE       FALSE
## 903         6.605857       FALSE       FALSE
## 904         6.703821       FALSE       FALSE
## 906         6.567514       FALSE       FALSE
## 907         6.558822       FALSE       FALSE
## 908         6.568915       FALSE       FALSE
## 909         6.687406       FALSE       FALSE
## 910         6.602563       FALSE       FALSE
```

There is no indication that results are changed by.

### Improvement in Implicit Fear Evaluation and Avoidance Tendencies

#### Implicit Fear Evaluation (EAST)

##### Testing the Random Model part

The linear mixed-effects model is created with spider anxiety as outcome and fixed effects of time as independent predictor. Random effect is a random intercept by subject.

```
model <- lmer(EAST ~ Sessionfac + (1 | Subject),
              data = dat, REML = FALSE)
```

Next, it is tested whether the model requires a random slope for Time (centred, linear variable) by testing the initial model against a more complex model.

```
model2 <- lmer(EAST ~ Sessionfac + (1 + Sessioncen | Subject),
              data = dat, REML = FALSE)
```

This is the result, which shows the random slope is not significant, so will be dropped in the model.

```
anova(model, model2)
```

```
## Data: dat
## Models:
## model: EAST ~ Sessionfac + (1 | Subject)
## model2: EAST ~ Sessionfac + (1 + Sessioncen | Subject)
##        Df  AIC    BIC  logLik deviance  Chisq Chi Df Pr(>Chisq)
## model   5 1111 1124.6 -550.48     1101
## model2  7 1111 1130.1 -548.49     1097 3.9664      2     0.1376
```

##### Testing the effect of interest

The base model is now tested against the same models without the effects of time:

```
model_null <- lmer(EAST ~  (1 | Subject),
              data = dat, REML = FALSE)
```

These are the results, which **does not** support rejection of null hypothesis:

```
anova(model, model_null)
```

```
## Data: dat
## Models:
## model_null: EAST ~ (1 | Subject)
## model: EAST ~ Sessionfac + (1 | Subject)
##            Df    AIC    BIC  logLik deviance  Chisq Chi Df Pr(>Chisq)
## model_null  3 1110.3 1118.5 -552.13   1104.3
## model       5 1111.0 1124.6 -550.48   1101.0 3.3126      2     0.1908
```

This is the final model (re-estimated with REML=TRUE):

```
model <- lmer(EAST ~ Sessionfac + (1 | Subject),
              data = dat, REML = TRUE)

summary(model)
```

```
## Linear mixed model fit by REML ['lmerMod']
## Formula: EAST ~ Sessionfac + (1 | Subject)
```

```
##    Data: dat
##
## REML criterion at convergence: 1085.6
##
## Scaled residuals:
##     Min      1Q  Median      3Q     Max
## -3.8151 -0.3900  0.0517  0.4372  3.0017
##
## Random effects:
##  Groups   Name        Variance Std.Dev.
##  Subject  (Intercept) 538.7    23.21
##  Residual             682.7    26.13
## Number of obs: 113, groups:  Subject, 38
##
## Fixed effects:
##             Estimate Std. Error t value
## (Intercept)  -10.724      5.669  -1.892
## Sessionfac2    7.632      5.994   1.273
## Sessionfac3   10.606      6.047   1.754
##
## Correlation of Fixed Effects:
##             (Intr) Sssnf2
## Sessionfac2 -0.529
## Sessionfac3 -0.524  0.496
```

##### Assumptions

Here are the residuals plotted against fitted values to investigate underlying heteroskedasticity and linearity assumptions:

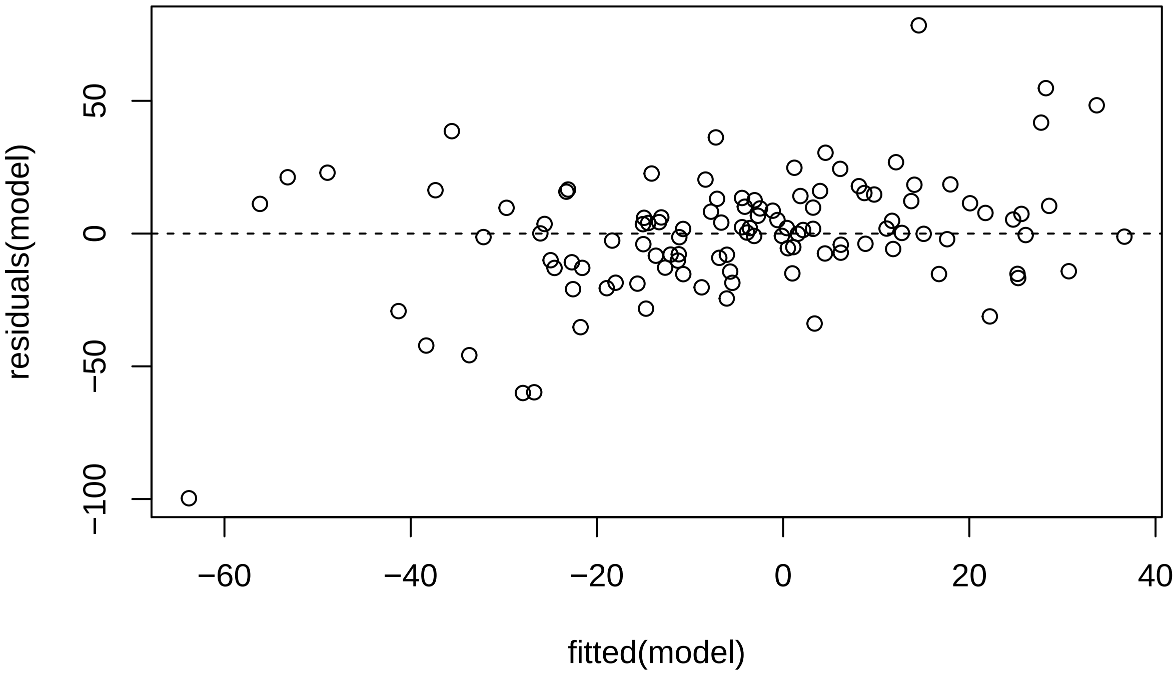

And the assumption of normality of residuals with histogram and qqplot:

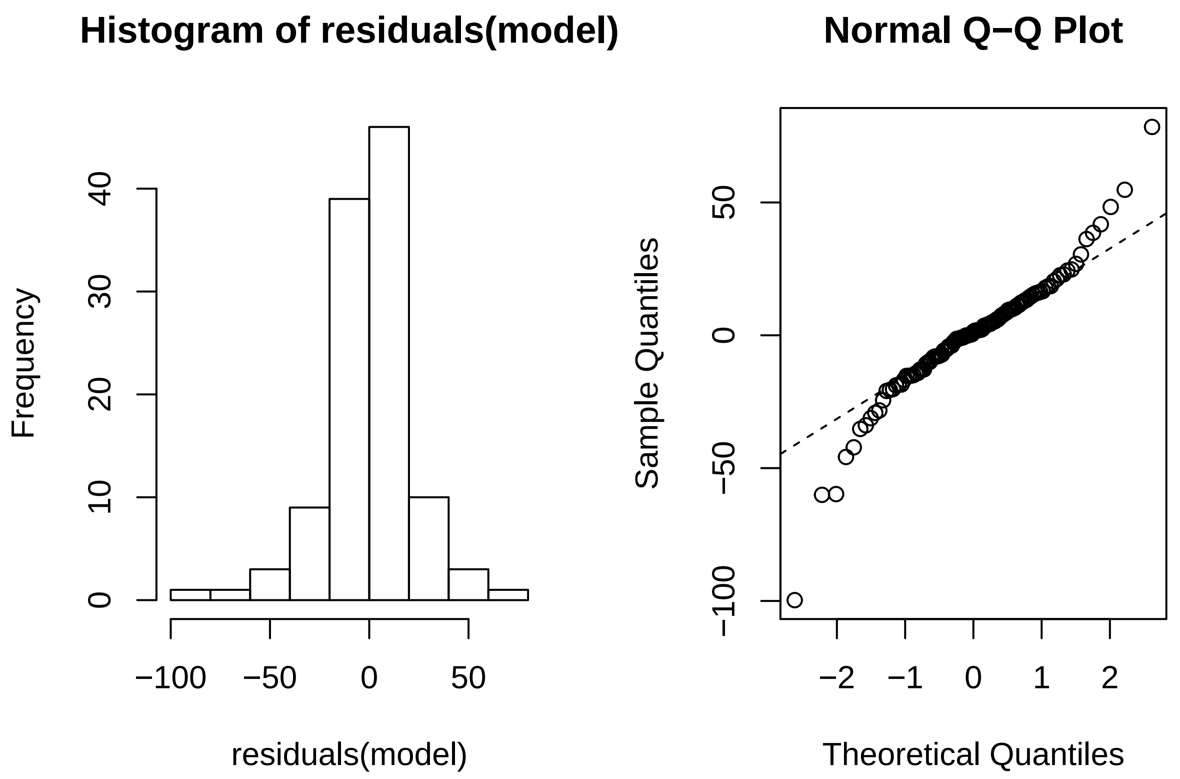

Using *Influence.ME*, outlying subjects are checked:

```
outlier <- influence(model, group = "Subject")
cks.d <- cooks.distance(outlier)
dfbetas <- dfbetas(outlier)
```

Now the Cook's distance values will be plotted. Outlier Subjects are highlighted in red:

```
plot(outlier,
     which = "cook",
     sort = TRUE,
     cutoff = 4/length(unique(dat$Subject)),
     xlab = "Cooks Distance")
```

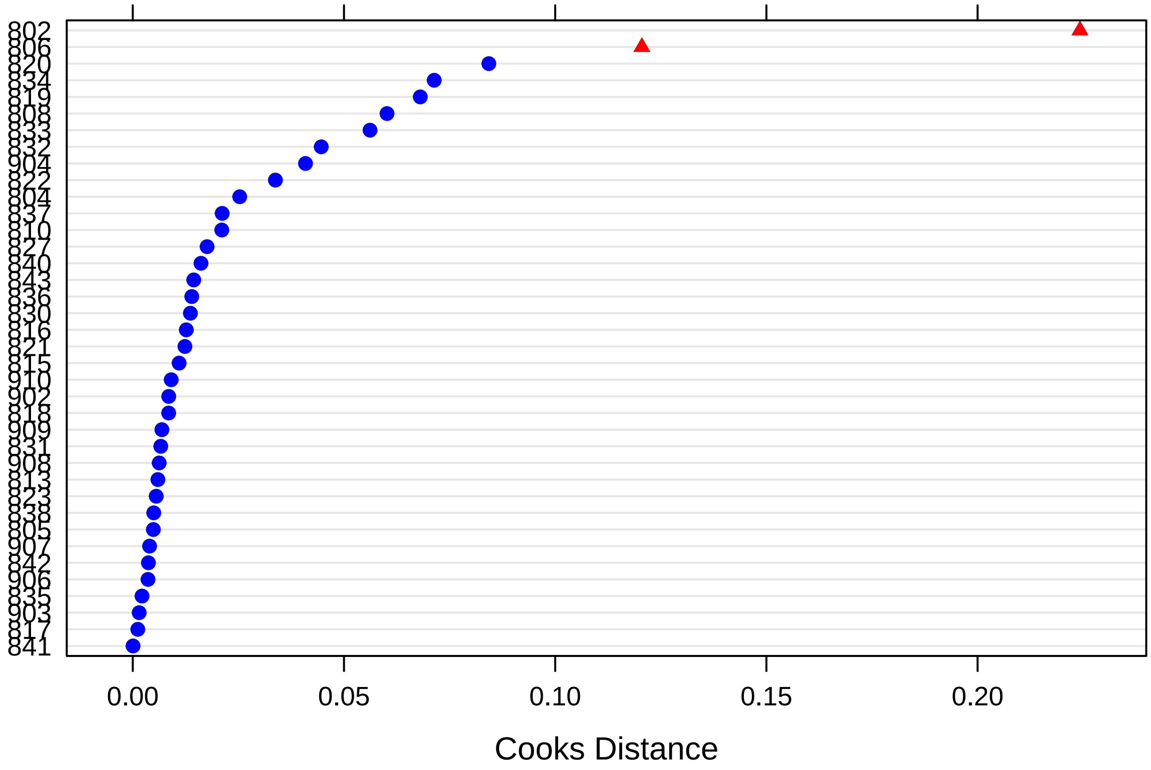

There are two outlying subjects (*802* & *806*).

Finally, these are the p-values for dfbeta values, that indicate whether exclusion of a specific subject suffices to change significance values:

```r
sigtest(outlier, test = -1.96)
```

```
## $Intercept
##     Altered.Teststat Altered.Sig Changed.Sig
## 802        -1.264867       FALSE       FALSE
## 804        -1.641525       FALSE       FALSE
## 805        -1.741811       FALSE       FALSE
## 806        -2.425498        TRUE        TRUE
## 808        -2.022775        TRUE        TRUE
## 810        -2.100744        TRUE        TRUE
## 813        -1.879636       FALSE       FALSE
## 815        -2.022552        TRUE        TRUE
## 816        -1.867667       FALSE       FALSE
## 817        -1.803113       FALSE       FALSE
## 818        -1.799597       FALSE       FALSE
## 819        -1.849915       FALSE       FALSE
## 820        -1.970582        TRUE        TRUE
## 821        -1.754654       FALSE       FALSE
## 822        -2.045395        TRUE        TRUE
## 823        -1.915184       FALSE       FALSE
## 827        -2.052292        TRUE        TRUE
## 830        -1.715921       FALSE       FALSE
```

```
## 831        -1.874970       FALSE       FALSE
## 832        -2.096219        TRUE        TRUE
## 833        -2.088543        TRUE        TRUE
## 834        -1.851313       FALSE       FALSE
## 835        -1.883840       FALSE       FALSE
## 836        -1.891646       FALSE       FALSE
## 837        -1.760117       FALSE       FALSE
## 838        -1.797507       FALSE       FALSE
## 840        -2.071847        TRUE        TRUE
## 841        -1.848937       FALSE       FALSE
## 842        -1.935823       FALSE       FALSE
## 843        -1.698677       FALSE       FALSE
## 902        -1.702699       FALSE       FALSE
## 903        -1.843192       FALSE       FALSE
## 904        -1.553413       FALSE       FALSE
## 906        -1.851030       FALSE       FALSE
## 907        -1.790837       FALSE       FALSE
## 908        -1.861706       FALSE       FALSE
## 909        -1.883233       FALSE       FALSE
## 910        -1.760801       FALSE       FALSE
##
## $Sessionfac2
##     Altered.Teststat Altered.Sig Changed.Sig
## 802        0.8337246       FALSE       FALSE
## 804        1.0026333       FALSE       FALSE
## 805        1.1772831       FALSE       FALSE
## 806        1.8591935       FALSE       FALSE
## 808        1.3435232       FALSE       FALSE
## 810        1.2887463       FALSE       FALSE
## 813        1.1520726       FALSE       FALSE
## 815        1.3728731       FALSE       FALSE
## 816        1.4007503       FALSE       FALSE
## 817        1.1968039       FALSE       FALSE
## 818        1.3395396       FALSE       FALSE
## 819        1.3168516       FALSE       FALSE
## 820        1.7621523       FALSE       FALSE
## 821        1.2911352       FALSE       FALSE
## 822        1.4250479       FALSE       FALSE
## 823        1.1818866       FALSE       FALSE
## 827        1.3032130       FALSE       FALSE
## 830        1.2048171       FALSE       FALSE
## 831        1.1397851       FALSE       FALSE
## 832        1.1239726       FALSE       FALSE
## 833        1.0419727       FALSE       FALSE
## 834        1.5046912       FALSE       FALSE
## 835        1.2078283       FALSE       FALSE
## 836        1.0955622       FALSE       FALSE
## 837        1.1502167       FALSE       FALSE
## 838        1.2981334       FALSE       FALSE
## 840        1.2869300       FALSE       FALSE
## 841        1.2505636       FALSE       FALSE
## 842        1.2928074       FALSE       FALSE
## 843        1.0692618       FALSE       FALSE
## 902        1.1325621       FALSE       FALSE
```

```
## 903        1.2978112       FALSE       FALSE
## 904        1.0608293       FALSE       FALSE
## 906        1.3243595       FALSE       FALSE
## 907        1.2410601       FALSE       FALSE
## 908        1.3536162       FALSE       FALSE
## 909        1.3297048       FALSE       FALSE
## 910        1.1202422       FALSE       FALSE
##
## $Sessionfac3
##     Altered.Teststat Altered.Sig Changed.Sig
## 802         1.305604       FALSE       FALSE
## 804         1.586335       FALSE       FALSE
## 805         1.697418       FALSE       FALSE
## 806         2.283895       FALSE       FALSE
## 808         1.495367       FALSE       FALSE
## 810         1.743229       FALSE       FALSE
## 813         1.666507       FALSE       FALSE
## 815         1.851650       FALSE       FALSE
## 816         1.845820       FALSE       FALSE
## 817         1.651157       FALSE       FALSE
## 818         1.745106       FALSE       FALSE
## 819         2.124955       FALSE       FALSE
## 820         2.021291       FALSE       FALSE
## 821         1.761654       FALSE       FALSE
## 822         2.067920       FALSE       FALSE
## 823         1.713872       FALSE       FALSE
## 827         1.897322       FALSE       FALSE
## 830         1.539531       FALSE       FALSE
## 831         1.679116       FALSE       FALSE
## 832         1.851345       FALSE       FALSE
## 833         1.740546       FALSE       FALSE
## 834         2.048843       FALSE       FALSE
## 835         1.737657       FALSE       FALSE
## 836         1.685876       FALSE       FALSE
## 837         1.485538       FALSE       FALSE
## 838         1.662591       FALSE       FALSE
## 840         1.765817       FALSE       FALSE
## 841         1.718058       FALSE       FALSE
## 842         1.801129       FALSE       FALSE
## 843         1.554479       FALSE       FALSE
## 902         1.646759       FALSE       FALSE
## 903         1.723745       FALSE       FALSE
## 904         1.464348       FALSE       FALSE
## 906         1.699567       FALSE       FALSE
## 907         1.618347       FALSE       FALSE
## 908         1.698217       FALSE       FALSE
## 909         1.660468       FALSE       FALSE
## 910         1.565391       FALSE       FALSE
```

There is no indication that results are changed.

#### Avoidance (AAT)

##### Testing the Random Model part

The linear mixed-effects model is created with spider anxiety as outcome and fixed effects of time as independent predictor. Random effect is a random intercept by subject.

```
model <- lmer(AAT ~ Sessionfac + (1 | Subject),
              data = dat, REML = FALSE)
```

Next, it is tested whether the model requires a random slope for Time (centred, linear variable) by testing the initial model against a more complex model.

```
model2 <- lmer(AAT ~ Sessionfac + (1 + Sessioncen | Subject),
              data = dat, REML = FALSE)
```

```
## singular fit
```

This is the result, which shows the random slope is significant, so will be retained in the model.

```
anova(model, model2)
```

```
## Data: dat
## Models:
## model: AAT ~ Sessionfac + (1 | Subject)
## model2: AAT ~ Sessionfac + (1 + Sessioncen | Subject)
##        Df    AIC    BIC  logLik deviance  Chisq Chi Df Pr(>Chisq)
## model   5 1270.3 1283.8 -630.13   1260.3
## model2  7 1236.6 1255.5 -611.29   1222.6 37.691      2  6.537e-09 ***
## ---
## Signif. codes:  0 '***' 0.001 '**' 0.01 '*' 0.05 '.' 0.1 ' ' 1
```

##### Testing the effect of interest

The base model is now tested against the same models without the effects of time:

```
model_null <- lmer(AAT ~  (1 + Sessioncen | Subject),
              data = dat, REML = FALSE)
```

```
## Warning in checkConv(attr(opt, "derivs"), opt$par, ctrl =
## control$checkConv, : Model failed to converge with max|grad| = 0.00380182
## (tol = 0.002, component 1)
```

These are the results, which **does not** support rejection of null hypothesis:

```
anova(model2, model_null)
```

```
## Data: dat
## Models:
## model_null: AAT ~ (1 + Sessioncen | Subject)
## model2: AAT ~ Sessionfac + (1 + Sessioncen | Subject)
##            Df    AIC    BIC  logLik deviance Chisq Chi Df Pr(>Chisq)
## model_null  5 1233.5 1247.0 -611.72   1223.5
## model2      7 1236.6 1255.5 -611.29   1222.6  0.87      2     0.6473
```

This is the final model (re-estimated with REML=TRUE):

```
model <- lmer(AAT ~ Sessionfac + (1 + Sessioncen | Subject),
              data = dat, REML = TRUE)
```

```
## singular fit
```

```r
summary(model)
```

```
## Linear mixed model fit by REML ['lmerMod']
## Formula: AAT ~ Sessionfac + (1 + Sessioncen | Subject)
##    Data: dat
##
## REML criterion at convergence: 1203.3
##
## Scaled residuals:
##      Min       1Q   Median       3Q      Max
## -2.36615 -0.48824  0.05816  0.52094  1.91536
##
## Random effects:
##  Groups   Name        Variance Std.Dev. Corr
##  Subject  (Intercept) 2592     50.91
##           Sessioncen  1775     42.13    -1.00
##  Residual             1909     43.69
## Number of obs: 111, groups:  Subject, 38
##
## Fixed effects:
##             Estimate Std. Error t value
## (Intercept)   -12.72      16.79  -0.758
## Sessionfac2     0.20      12.44   0.016
## Sessionfac3    11.50      17.07   0.674
##
## Correlation of Fixed Effects:
##             (Intr) Sssnf2
## Sessionfac2 -0.754
## Sessionfac3 -0.909  0.696
## convergence code: 0
## singular fit
```

##### Assumptions

Here are the residuals plotted against fitted values to investigate underlying heteroskedasticity and linearity assumptions:

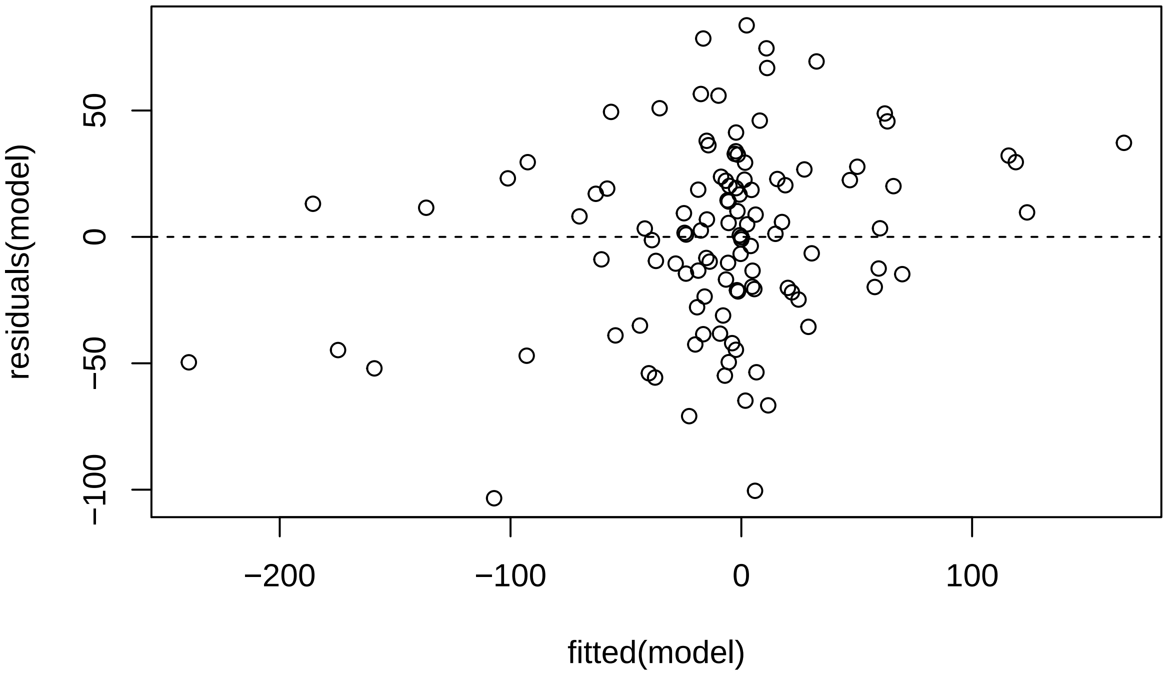

And the assumption of normality of residuals with histogram and qqplot:

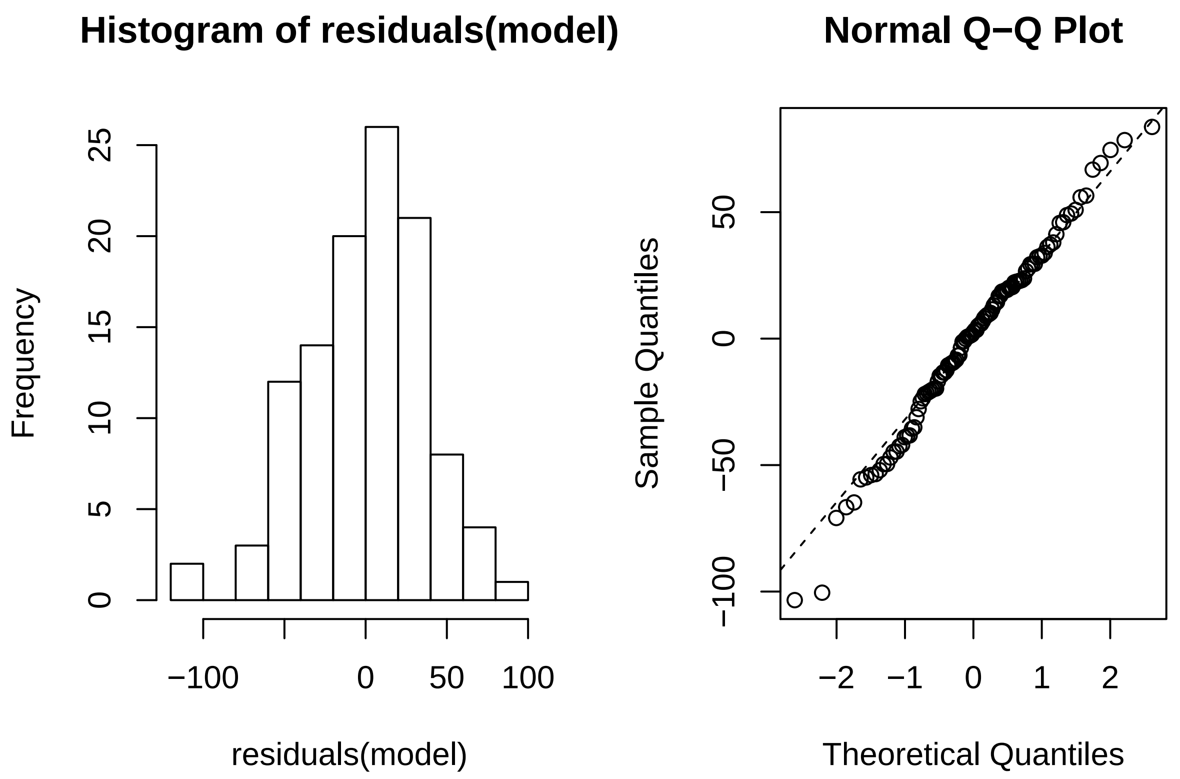

Using *Influence.ME*, outlying subjects are checked:

```
outlier <- influence(model, group = "Subject")
```

```
## singular fit
## singular fit
## singular fit
## singular fit
## singular fit
## singular fit
## singular fit
## singular fit
## singular fit
## singular fit
## singular fit
## singular fit
## singular fit
## singular fit
## singular fit
## singular fit
## singular fit
## singular fit
## singular fit
## singular fit
## singular fit
## singular fit
## singular fit
```

```
## singular fit
## singular fit
```

```r
cks.d <- cooks.distance(outlier)
dfbetas <- dfbetas(outlier)
```

Now the Cook's distance values will be plotted. Outlier Subjects are highlighted in red:

```r
plot(outlier,
     which = "cook",
     sort = TRUE,
     cutoff = 4/length(unique(dat$Subject)),
     xlab = "Cooks Distance")
```

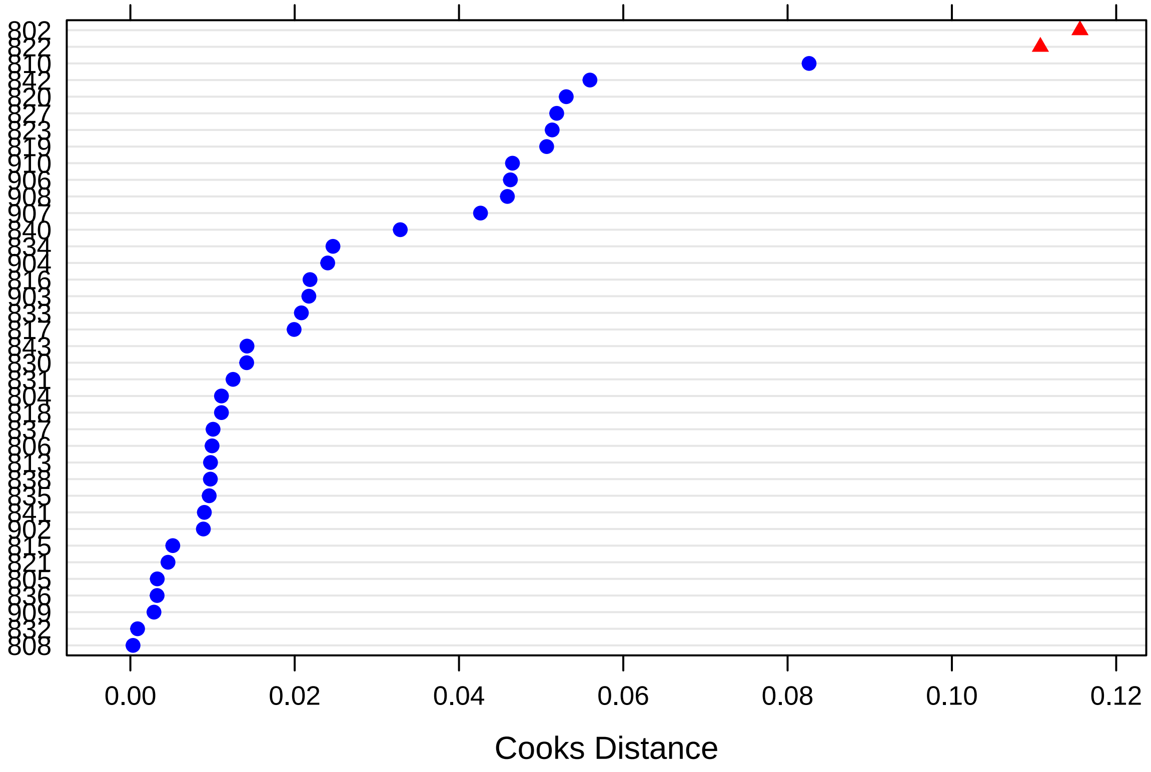

There is one outlying subject (*822*).

Finally, these are the p-values for dfbeta values, that indicate whether exclusion of a specific subject suffices to change significance values:

```r
sigtest(outlier, test = -1.96)
```

```
## $Intercept
##     Altered.Teststat Altered.Sig Changed.Sig
## 802       -0.3136201       FALSE       FALSE
## 804       -0.8657812       FALSE       FALSE
## 805       -0.7828773       FALSE       FALSE
## 806       -0.7351341       FALSE       FALSE
## 808       -0.7572758       FALSE       FALSE
## 810       -0.9581405       FALSE       FALSE
```

```
## 813       -0.8739513       FALSE       FALSE
## 815       -0.7954276       FALSE       FALSE
## 816       -0.8102588       FALSE       FALSE
## 817       -0.6909252       FALSE       FALSE
## 818       -0.6635958       FALSE       FALSE
## 819       -0.7563807       FALSE       FALSE
## 820       -1.0160940       FALSE       FALSE
## 821       -0.6957671       FALSE       FALSE
## 822       -0.5196501       FALSE       FALSE
## 823       -0.4419273       FALSE       FALSE
## 827       -0.4233620       FALSE       FALSE
## 830       -0.6936103       FALSE       FALSE
## 831       -0.8911718       FALSE       FALSE
## 832       -0.7339962       FALSE       FALSE
## 833       -0.7017460       FALSE       FALSE
## 834       -0.8271070       FALSE       FALSE
## 835       -0.7224331       FALSE       FALSE
## 836       -0.6492257       FALSE       FALSE
## 837       -0.7849922       FALSE       FALSE
## 838       -0.6977547       FALSE       FALSE
## 840       -0.5424761       FALSE       FALSE
## 841       -0.7224220       FALSE       FALSE
## 842       -1.0100705       FALSE       FALSE
## 843       -0.6769835       FALSE       FALSE
## 902       -0.9034969       FALSE       FALSE
## 903       -0.6937965       FALSE       FALSE
## 904       -0.6108154       FALSE       FALSE
## 906       -1.0176496       FALSE       FALSE
## 907       -0.6574334       FALSE       FALSE
## 908       -1.1409264       FALSE       FALSE
## 909       -0.7455786       FALSE       FALSE
## 910       -0.8563874       FALSE       FALSE
##
## $Sessionfac2
##     Altered.Teststat Altered.Sig Changed.Sig
## 802     -0.385000395       FALSE       FALSE
## 804      0.032239449       FALSE       FALSE
## 805      0.032878821       FALSE       FALSE
## 806      0.090718506       FALSE       FALSE
## 808      0.015016414       FALSE       FALSE
## 810      0.305667556       FALSE       FALSE
## 813      0.170041285       FALSE       FALSE
## 815      0.065866027       FALSE       FALSE
## 816      0.230672216       FALSE       FALSE
## 817      0.131253219       FALSE       FALSE
## 818      0.057325989       FALSE       FALSE
## 819     -0.109051656       FALSE       FALSE
## 820      0.252546843       FALSE       FALSE
## 821     -0.070147760       FALSE       FALSE
## 822      0.122315570       FALSE       FALSE
## 823     -0.319640435       FALSE       FALSE
## 827     -0.311880125       FALSE       FALSE
## 830     -0.122505977       FALSE       FALSE
## 831      0.192655061       FALSE       FALSE
```

```
## 832      0.035875432       FALSE       FALSE
## 833      0.145086505       FALSE       FALSE
## 834      0.130380623       FALSE       FALSE
## 835      0.009822969       FALSE       FALSE
## 836     -0.048875913       FALSE       FALSE
## 837      0.161372798       FALSE       FALSE
## 838      0.019195466       FALSE       FALSE
## 840     -0.282422723       FALSE       FALSE
## 841      0.039133652       FALSE       FALSE
## 842      0.064539934       FALSE       FALSE
## 843     -0.005880279       FALSE       FALSE
## 902      0.150168480       FALSE       FALSE
## 903     -0.176863607       FALSE       FALSE
## 904     -0.228236563       FALSE       FALSE
## 906      0.230877553       FALSE       FALSE
## 907     -0.266432944       FALSE       FALSE
## 908      0.274385186       FALSE       FALSE
## 909      0.051957565       FALSE       FALSE
## 910     -0.094354825       FALSE       FALSE
##
## $Sessionfac3
##     Altered.Teststat Altered.Sig Changed.Sig
## 802        0.3696658       FALSE       FALSE
## 804        0.8030864       FALSE       FALSE
## 805        0.6605718       FALSE       FALSE
## 806        0.6043900       FALSE       FALSE
## 808        0.6747939       FALSE       FALSE
## 810        1.0496882       FALSE       FALSE
## 813        0.7514041       FALSE       FALSE
## 815        0.6621190       FALSE       FALSE
## 816        0.7395753       FALSE       FALSE
## 817        0.5952183       FALSE       FALSE
## 818        0.6149015       FALSE       FALSE
## 819        0.5384495       FALSE       FALSE
## 820        0.7766977       FALSE       FALSE
## 821        0.5872945       FALSE       FALSE
## 822        0.3730856       FALSE       FALSE
## 823        0.3275215       FALSE       FALSE
## 827        0.4236086       FALSE       FALSE
## 830        0.5641395       FALSE       FALSE
## 831        0.8318167       FALSE       FALSE
## 832        0.6375466       FALSE       FALSE
## 833        0.6061658       FALSE       FALSE
## 834        0.8466087       FALSE       FALSE
## 835        0.7122696       FALSE       FALSE
## 836        0.5905260       FALSE       FALSE
## 837        0.7021009       FALSE       FALSE
## 838        0.6844976       FALSE       FALSE
## 840        0.4420366       FALSE       FALSE
## 841        0.5802722       FALSE       FALSE
## 842        1.0214452       FALSE       FALSE
## 843        0.6780215       FALSE       FALSE
## 902        0.7913598       FALSE       FALSE
## 903        0.5875289       FALSE       FALSE
```

```
## 904         0.5090522       FALSE       FALSE
## 906         0.7889518       FALSE       FALSE
## 907         0.6142172       FALSE       FALSE
## 908         0.9672931       FALSE       FALSE
## 909         0.6976151       FALSE       FALSE
## 910         0.8497077       FALSE       FALSE
```

There is no indication that results are changed.

### DCS Effect on Spider Anxiety and Bias Measures

For analysis of this hypothesis, the Group variable (centred) will be added to lme models identified in hypothesis 1.

#### SAS

The base model is now tested against the same models without the effects of time:

```
model <- lmer(SAS ~ Timefac*Groupcen + (1 + Timecen | Subject),
              data = dat, REML = FALSE)

model_null <- lmer(SAS ~ Timefac + Groupcen + (1 + Timecen | Subject),
              data = dat, REML = FALSE)
```

These are the results, which **do not** support rejection of null hypothesis:

```
anova(model, model_null)
```

```
## Data: dat
## Models:
## model_null: SAS ~ Timefac + Groupcen + (1 + Timecen | Subject)
## model: SAS ~ Timefac * Groupcen + (1 + Timecen | Subject)
##            Df    AIC    BIC  logLik deviance  Chisq Chi Df Pr(>Chisq)
## model_null  9 775.75 802.96 -378.87   757.75
## model      12 780.21 816.50 -378.10   756.21 1.5389      3     0.6733
```

This is the final model (re-estimated with REML=TRUE):

```
model <- lmer(SAS ~ Timefac*Groupcen + (1 + Timecen | Subject),
              data = dat, REML = TRUE)

summary(model)
```

```
## Linear mixed model fit by REML ['lmerMod']
## Formula: SAS ~ Timefac * Groupcen + (1 + Timecen | Subject)
##    Data: dat
##
## REML criterion at convergence: 753.3
##
## Scaled residuals:
##     Min      1Q  Median      3Q     Max
## -1.9368 -0.4989  0.1015  0.5188  2.6850
##
## Random effects:
##  Groups   Name        Variance Std.Dev. Corr
```

```
##  Subject  (Intercept) 11.0860  3.3296
##           Timecen      0.4398  0.6632   0.78
##  Residual              4.0697  2.0174
## Number of obs: 152, groups:  Subject, 38
##
## Fixed effects:
##                   Estimate Std. Error t value
## (Intercept)        19.8067     0.4841  40.916
## Timefac2           -5.7731     0.5132 -11.248
## Timefac3           -7.1499     0.6355 -11.251
## Timefac4           -8.9328     0.7987 -11.184
## Groupcen            0.3361     0.4841   0.694
## Timefac2:Groupcen   0.3445     0.5132   0.671
## Timefac3:Groupcen   0.6737     0.6355   1.060
## Timefac4:Groupcen   0.3613     0.7987   0.452
##
## Correlation of Fixed Effects:
##             (Intr) Timfc2 Timfc3 Timfc4 Gropcn Tmf2:G Tmf3:G
## Timefac2    -0.351
## Timefac3    -0.214  0.619
## Timefac4    -0.116  0.607  0.767
## Groupcen    -0.105  0.037  0.023  0.012
## Tmfc2:Grpcn  0.037 -0.105 -0.065 -0.064 -0.351
## Tmfc3:Grpcn  0.023 -0.065 -0.105 -0.081 -0.214  0.619
## Tmfc4:Grpcn  0.012 -0.064 -0.081 -0.105 -0.116  0.607  0.767
```

##### Assumptions

Here are the residuals plotted against fitted values to investigate underlying heteroskedasticity and linearity assumptions:

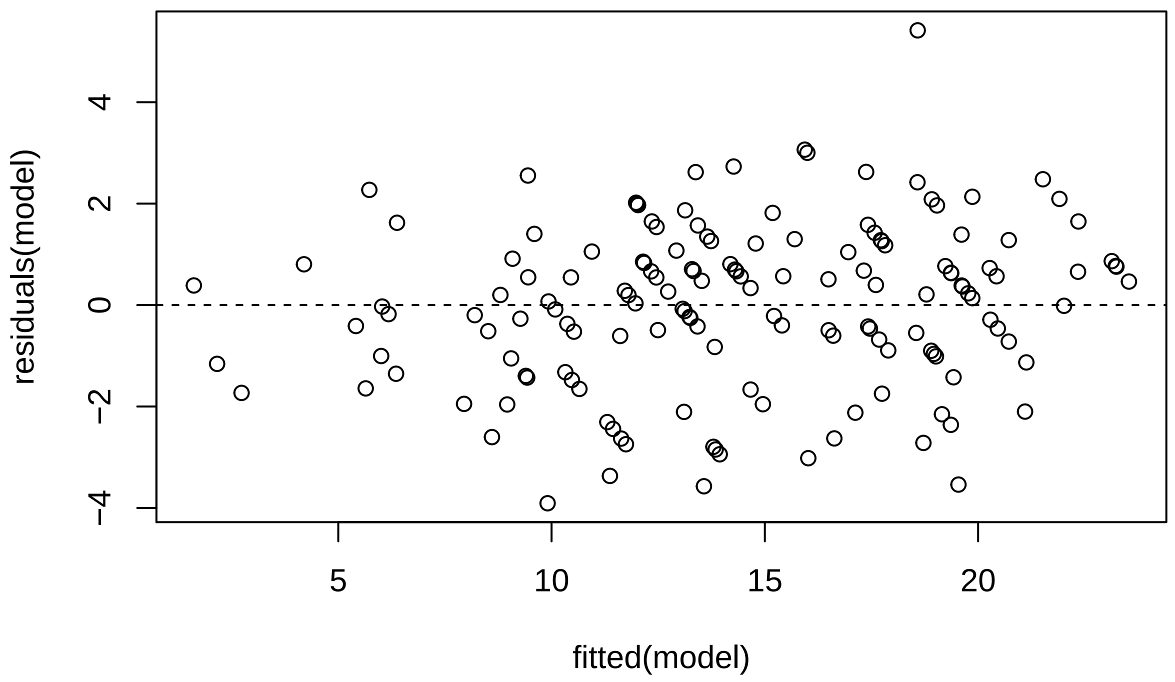

And the assumption of normality of residuals with histogram and qqplot:

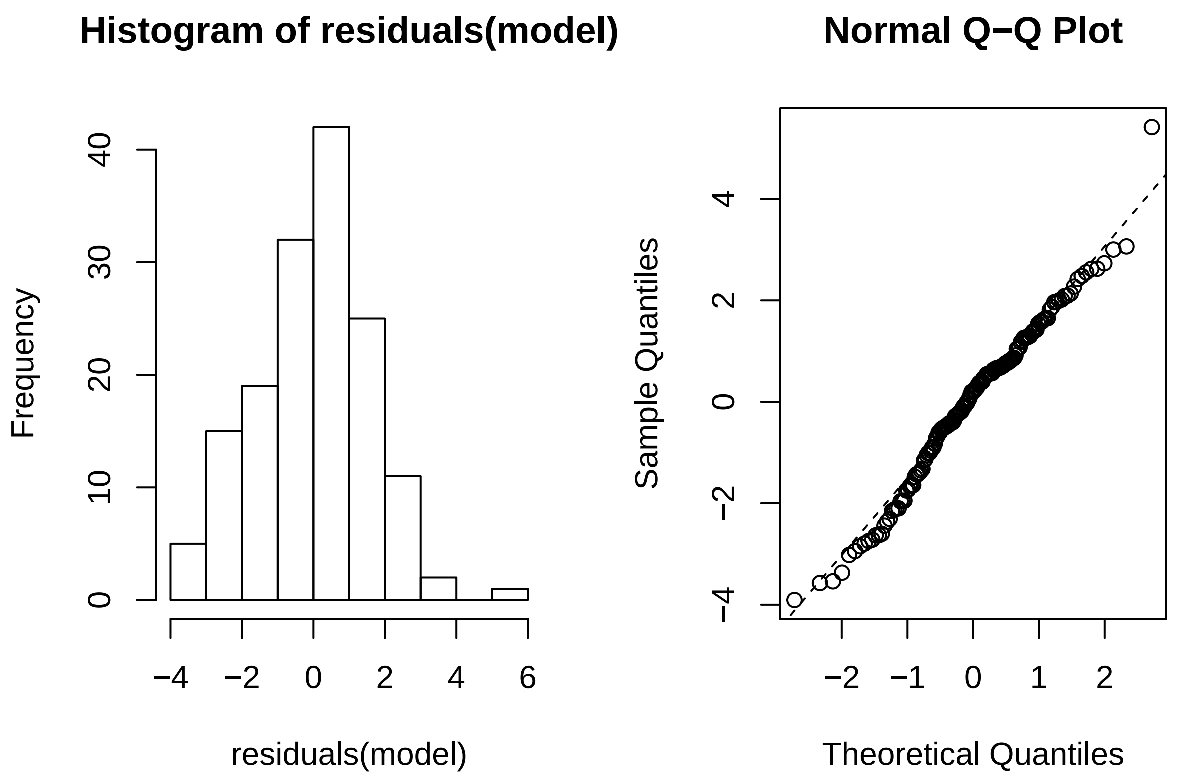

Using *Influence.ME*, outlying subjects are checked:

```
outlier <- influence(model, group = "Subject")
cks.d <- cooks.distance(outlier)
dfbetas <- dfbetas(outlier)
```

Now the Cook's distance values will be plotted. Outlier Subjects are highlighted in red:

```
plot(outlier,
     which = "cook",
     sort = TRUE,
     cutoff = 4/length(unique(dat$Subject)),
     xlab = "Cooks Distance")
```

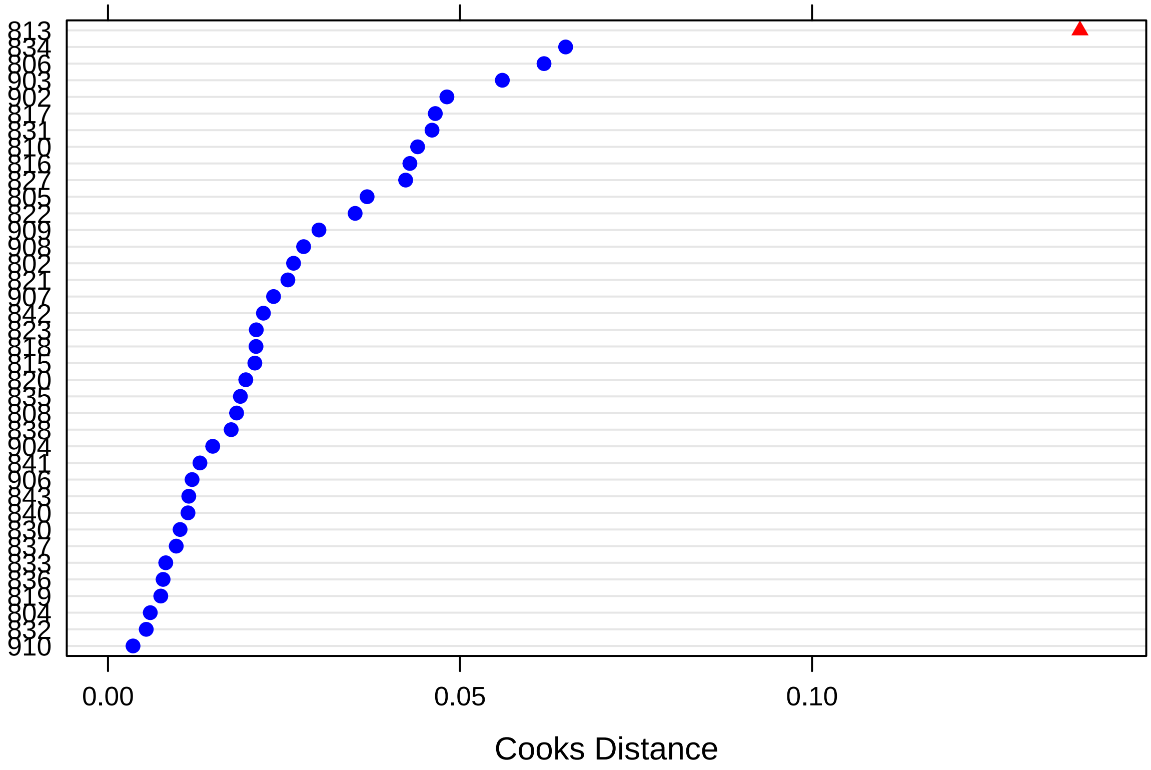

There is one outlying subject based on Cook's distance (i.e., *813*).

Finally, these are the p-values for dfbeta values, that indicate whether exclusion of a specific subject suffices to change significance values:

```
sigtest(outlier, test = -1.96)
```

```
## $Intercept
##     Altered.Teststat Altered.Sig Changed.Sig
## 802         40.93617       FALSE       FALSE
## 804         39.76386       FALSE       FALSE
## 805         40.88687       FALSE       FALSE
## 806         40.46389       FALSE       FALSE
## 808         39.67496       FALSE       FALSE
## 810         40.84758       FALSE       FALSE
## 813         39.88017       FALSE       FALSE
## 815         39.98715       FALSE       FALSE
## 816         41.09879       FALSE       FALSE
## 817         40.13600       FALSE       FALSE
## 818         40.49368       FALSE       FALSE
## 819         40.23092       FALSE       FALSE
## 820         40.05859       FALSE       FALSE
## 821         40.25877       FALSE       FALSE
## 822         41.58936       FALSE       FALSE
## 823         40.28762       FALSE       FALSE
## 827         40.79062       FALSE       FALSE
## 830         40.16801       FALSE       FALSE
```

```
## 831         39.63877       FALSE       FALSE
## 832         39.68203       FALSE       FALSE
## 833         39.99517       FALSE       FALSE
## 834         42.21988       FALSE       FALSE
## 835         40.86852       FALSE       FALSE
## 836         39.81053       FALSE       FALSE
## 837         40.30357       FALSE       FALSE
## 838         39.93558       FALSE       FALSE
## 840         39.97511       FALSE       FALSE
## 841         40.49332       FALSE       FALSE
## 842         39.93613       FALSE       FALSE
## 843         40.36494       FALSE       FALSE
## 902         40.00506       FALSE       FALSE
## 903         40.53588       FALSE       FALSE
## 904         39.97894       FALSE       FALSE
## 906         39.91385       FALSE       FALSE
## 907         39.78254       FALSE       FALSE
## 908         40.47379       FALSE       FALSE
## 909         42.43832       FALSE       FALSE
## 910         40.23901       FALSE       FALSE
##
## $Timefac2
##     Altered.Teststat Altered.Sig Changed.Sig
## 802        -11.03573        TRUE       FALSE
## 804        -11.06106        TRUE       FALSE
## 805        -10.94942        TRUE       FALSE
## 806        -11.23341        TRUE       FALSE
## 808        -10.93046        TRUE       FALSE
## 810        -10.93586        TRUE       FALSE
## 813        -11.05608        TRUE       FALSE
## 815        -11.28199        TRUE       FALSE
## 816        -10.97918        TRUE       FALSE
## 817        -11.59020        TRUE       FALSE
## 818        -10.89208        TRUE       FALSE
## 819        -11.13102        TRUE       FALSE
## 820        -11.17267        TRUE       FALSE
## 821        -11.30621        TRUE       FALSE
## 822        -11.19685        TRUE       FALSE
## 823        -11.26704        TRUE       FALSE
## 827        -11.10123        TRUE       FALSE
## 830        -11.08198        TRUE       FALSE
## 831        -10.84162        TRUE       FALSE
## 832        -10.88824        TRUE       FALSE
## 833        -10.99645        TRUE       FALSE
## 834        -11.37113        TRUE       FALSE
## 835        -11.06841        TRUE       FALSE
## 836        -11.08708        TRUE       FALSE
## 837        -10.90406        TRUE       FALSE
## 838        -11.26159        TRUE       FALSE
## 840        -11.14277        TRUE       FALSE
## 841        -10.92762        TRUE       FALSE
## 842        -11.00001        TRUE       FALSE
## 843        -11.06747        TRUE       FALSE
## 902        -11.05196        TRUE       FALSE
```

```
## 903        -11.79393        TRUE       FALSE
## 904        -11.24478        TRUE       FALSE
## 906        -10.97440        TRUE       FALSE
## 907        -10.90082        TRUE       FALSE
## 908        -10.92320        TRUE       FALSE
## 909        -11.05758        TRUE       FALSE
## 910        -10.99641        TRUE       FALSE
##
## $Timefac3
##     Altered.Teststat Altered.Sig Changed.Sig
## 802        -11.19090        TRUE       FALSE
## 804        -11.01179        TRUE       FALSE
## 805        -10.91688        TRUE       FALSE
## 806        -11.51332        TRUE       FALSE
## 808        -10.77281        TRUE       FALSE
## 810        -11.06170        TRUE       FALSE
## 813        -11.28085        TRUE       FALSE
## 815        -11.35780        TRUE       FALSE
## 816        -11.05621        TRUE       FALSE
## 817        -11.35575        TRUE       FALSE
## 818        -11.00846        TRUE       FALSE
## 819        -11.08194        TRUE       FALSE
## 820        -10.95962        TRUE       FALSE
## 821        -11.33200        TRUE       FALSE
## 822        -11.15581        TRUE       FALSE
## 823        -10.95479        TRUE       FALSE
## 827        -11.27696        TRUE       FALSE
## 830        -11.15847        TRUE       FALSE
## 831        -11.08045        TRUE       FALSE
## 832        -10.93316        TRUE       FALSE
## 833        -11.00845        TRUE       FALSE
## 834        -11.26796        TRUE       FALSE
## 835        -10.96293        TRUE       FALSE
## 836        -10.92972        TRUE       FALSE
## 837        -10.97643        TRUE       FALSE
## 838        -11.35203        TRUE       FALSE
## 840        -11.02141        TRUE       FALSE
## 841        -11.00413        TRUE       FALSE
## 842        -11.15022        TRUE       FALSE
## 843        -11.01865        TRUE       FALSE
## 902        -10.94184        TRUE       FALSE
## 903        -11.44425        TRUE       FALSE
## 904        -11.18887        TRUE       FALSE
## 906        -11.03168        TRUE       FALSE
## 907        -11.11575        TRUE       FALSE
## 908        -10.91295        TRUE       FALSE
## 909        -11.03751        TRUE       FALSE
## 910        -10.99267        TRUE       FALSE
##
## $Timefac4
##     Altered.Teststat Altered.Sig Changed.Sig
## 802        -11.22271        TRUE       FALSE
## 804        -10.89996        TRUE       FALSE
## 805        -10.86568        TRUE       FALSE
```

```
## 806        -11.78292        TRUE       FALSE
## 808        -10.83261        TRUE       FALSE
## 810        -10.75747        TRUE       FALSE
## 813        -11.38211        TRUE       FALSE
## 815        -11.24046        TRUE       FALSE
## 816        -11.14377        TRUE       FALSE
## 817        -11.03650        TRUE       FALSE
## 818        -10.93524        TRUE       FALSE
## 819        -11.07437        TRUE       FALSE
## 820        -11.00522        TRUE       FALSE
## 821        -11.14234        TRUE       FALSE
## 822        -10.93286        TRUE       FALSE
## 823        -10.93745        TRUE       FALSE
## 827        -11.35029        TRUE       FALSE
## 830        -11.04366        TRUE       FALSE
## 831        -11.24699        TRUE       FALSE
## 832        -10.77290        TRUE       FALSE
## 833        -10.89360        TRUE       FALSE
## 834        -10.91258        TRUE       FALSE
## 835        -10.85520        TRUE       FALSE
## 836        -10.82904        TRUE       FALSE
## 837        -10.89610        TRUE       FALSE
## 838        -11.46902        TRUE       FALSE
## 840        -11.06186        TRUE       FALSE
## 841        -10.89464        TRUE       FALSE
## 842        -11.02019        TRUE       FALSE
## 843        -11.01278        TRUE       FALSE
## 902        -11.10025        TRUE       FALSE
## 903        -10.98876        TRUE       FALSE
## 904        -11.10610        TRUE       FALSE
## 906        -10.98427        TRUE       FALSE
## 907        -11.01762        TRUE       FALSE
## 908        -10.94734        TRUE       FALSE
## 909        -10.92555        TRUE       FALSE
## 910        -10.89349        TRUE       FALSE
##
## $Groupcen
##     Altered.Teststat Altered.Sig Changed.Sig
## 802        0.8032643       FALSE       FALSE
## 804        0.7086273       FALSE       FALSE
## 805        0.7517752       FALSE       FALSE
## 806        0.9828913       FALSE       FALSE
## 808        0.7709098       FALSE       FALSE
## 810        0.4669864       FALSE       FALSE
## 813        0.9687124       FALSE       FALSE
## 815        0.5207127       FALSE       FALSE
## 816        0.5351885       FALSE       FALSE
## 817        0.6508541       FALSE       FALSE
## 818        0.4924622       FALSE       FALSE
## 819        0.7894257       FALSE       FALSE
## 820        0.6869225       FALSE       FALSE
## 821        0.4602549       FALSE       FALSE
## 822        0.5057872       FALSE       FALSE
## 823        0.7905383       FALSE       FALSE
```

```
## 827        0.9908278       FALSE       FALSE
## 830        0.5889036       FALSE       FALSE
## 831        0.7702065       FALSE       FALSE
## 832        0.7071690       FALSE       FALSE
## 833        0.6361652       FALSE       FALSE
## 834        0.5134552       FALSE       FALSE
## 835        0.4024653       FALSE       FALSE
## 836        0.5818944       FALSE       FALSE
## 837        0.7410502       FALSE       FALSE
## 838        0.6848131       FALSE       FALSE
## 840        0.6854911       FALSE       FALSE
## 841        0.4924578       FALSE       FALSE
## 842        0.5855041       FALSE       FALSE
## 843        0.5414076       FALSE       FALSE
## 902        0.8419258       FALSE       FALSE
## 903        0.8952085       FALSE       FALSE
## 904        0.5206059       FALSE       FALSE
## 906        0.6844407       FALSE       FALSE
## 907        0.7089602       FALSE       FALSE
## 908        0.7441800       FALSE       FALSE
## 909        1.0411819       FALSE       FALSE
## 910        0.7895845       FALSE       FALSE
##
## $`Timefac2:Groupcen`
##     Altered.Teststat Altered.Sig Changed.Sig
## 802       0.73873462       FALSE       FALSE
## 804       0.77799539       FALSE       FALSE
## 805       0.94031846       FALSE       FALSE
## 806       0.79011776       FALSE       FALSE
## 808       0.54652276       FALSE       FALSE
## 810       0.48966516       FALSE       FALSE
## 813      -0.05482354       FALSE       FALSE
## 815       0.90460742       FALSE       FALSE
## 816       0.54895896       FALSE       FALSE
## 817       1.04103604       FALSE       FALSE
## 818       0.83134868       FALSE       FALSE
## 819       0.54153959       FALSE       FALSE
## 820       0.64485024       FALSE       FALSE
## 821       0.90654972       FALSE       FALSE
## 822       0.54474229       FALSE       FALSE
## 823       0.54815690       FALSE       FALSE
## 827       0.78082040       FALSE       FALSE
## 830       0.58916870       FALSE       FALSE
## 831       0.37028551       FALSE       FALSE
## 832       0.60067202       FALSE       FALSE
## 833       0.68517186       FALSE       FALSE
## 834       0.60454118       FALSE       FALSE
## 835       0.77851220       FALSE       FALSE
## 836       0.77982488       FALSE       FALSE
## 837       0.78086801       FALSE       FALSE
## 838       0.49749989       FALSE       FALSE
## 840       0.59240035       FALSE       FALSE
## 841       0.73149752       FALSE       FALSE
## 842       0.83958711       FALSE       FALSE
```

```
## 843      0.58839730       FALSE       FALSE
## 902      0.55259797       FALSE       FALSE
## 903      0.36536338       FALSE       FALSE
## 904      0.90162361       FALSE       FALSE
## 906      0.73462897       FALSE       FALSE
## 907      0.48809649       FALSE       FALSE
## 908      0.88566518       FALSE       FALSE
## 909      0.58787127       FALSE       FALSE
## 910      0.63467734       FALSE       FALSE
##
## $`Timefac3:Groupcen`
##     Altered.Teststat Altered.Sig Changed.Sig
## 802        0.9915411       FALSE       FALSE
## 804        1.1164136       FALSE       FALSE
## 805        1.2205912       FALSE       FALSE
## 806        1.2562133       FALSE       FALSE
## 808        0.8335969       FALSE       FALSE
## 810        1.0345146       FALSE       FALSE
## 813        0.4885308       FALSE       FALSE
## 815        1.3254990       FALSE       FALSE
## 816        1.0340009       FALSE       FALSE
## 817        1.3252595       FALSE       FALSE
## 818        1.1875011       FALSE       FALSE
## 819        0.9818865       FALSE       FALSE
## 820        1.1394005       FALSE       FALSE
## 821        1.3224883       FALSE       FALSE
## 822        0.9045249       FALSE       FALSE
## 823        1.1388979       FALSE       FALSE
## 827        1.2304232       FALSE       FALSE
## 830        0.9047412       FALSE       FALSE
## 831        0.6720495       FALSE       FALSE
## 832        1.0224930       FALSE       FALSE
## 833        1.1017551       FALSE       FALSE
## 834        0.9983683       FALSE       FALSE
## 835        1.0685563       FALSE       FALSE
## 836        1.0221708       FALSE       FALSE
## 837        1.0562527       FALSE       FALSE
## 838        0.9204352       FALSE       FALSE
## 840        1.0605811       FALSE       FALSE
## 841        1.0168086       FALSE       FALSE
## 842        0.9879364       FALSE       FALSE
## 843        1.0181503       FALSE       FALSE
## 902        0.6636429       FALSE       FALSE
## 903        0.8010055       FALSE       FALSE
## 904        1.2208118       FALSE       FALSE
## 906        1.1900063       FALSE       FALSE
## 907        1.1269533       FALSE       FALSE
## 908        1.3069810       FALSE       FALSE
## 909        0.9779497       FALSE       FALSE
## 910        1.0157502       FALSE       FALSE
##
## $`Timefac4:Groupcen`
##     Altered.Teststat Altered.Sig Changed.Sig
## 802       0.30647204       FALSE       FALSE
```

```
## 804        0.48805772       FALSE       FALSE
## 805        0.51689892       FALSE       FALSE
## 806        0.83267464       FALSE       FALSE
## 808        0.41246425       FALSE       FALSE
## 810        0.22491389       FALSE       FALSE
## 813       -0.04762387       FALSE       FALSE
## 815        0.65082180       FALSE       FALSE
## 816        0.60903912       FALSE       FALSE
## 817        0.56709296       FALSE       FALSE
## 818        0.65079794       FALSE       FALSE
## 819        0.33394582       FALSE       FALSE
## 820        0.36336305       FALSE       FALSE
## 821        0.60896103       FALSE       FALSE
## 822        0.42408851       FALSE       FALSE
## 823        0.42426664       FALSE       FALSE
## 827        0.69378318       FALSE       FALSE
## 830        0.36463208       FALSE       FALSE
## 831        0.03503735       FALSE       FALSE
## 832        0.33699288       FALSE       FALSE
## 833        0.58290465       FALSE       FALSE
## 834        0.61659188       FALSE       FALSE
## 835        0.44981620       FALSE       FALSE
## 836        0.41232810       FALSE       FALSE
## 837        0.45437830       FALSE       FALSE
## 838        0.21632684       FALSE       FALSE
## 840        0.33356884       FALSE       FALSE
## 841        0.45431744       FALSE       FALSE
## 842        0.39557810       FALSE       FALSE
## 843        0.36361257       FALSE       FALSE
## 902        0.15393262       FALSE       FALSE
## 903        0.39444972       FALSE       FALSE
## 904        0.60698024       FALSE       FALSE
## 906        0.65371611       FALSE       FALSE
## 907        0.52985061       FALSE       FALSE
## 908        0.61855590       FALSE       FALSE
## 909        0.42380505       FALSE       FALSE
## 910        0.45426966       FALSE       FALSE
```

There is no indication that results are changed by an outlying subject.

#### FSQ

The base model is now tested against the same models without the effects of time:

```
model <- lmer(FSQ ~ Timefac*Groupcen + (1 + Timecen | Subject),
              data = dat, REML = FALSE)

model_null <- lmer(FSQ ~ Timefac + Groupcen + (1 + Timecen | Subject),
              data = dat, REML = FALSE)
```

These are the results, which **do not** support rejection of null hypothesis:

```
anova(model, model_null)
```

```
## Data: dat
```

```
## Models:
## model_null: FSQ ~ Timefac + Groupcen + (1 + Timecen | Subject)
## model: FSQ ~ Timefac * Groupcen + (1 + Timecen | Subject)
##            Df    AIC    BIC  logLik deviance Chisq Chi Df Pr(>Chisq)
## model_null  9 1212.2 1239.4 -597.09   1194.2
## model      12 1217.5 1253.8 -596.75   1193.5 0.683      3     0.8772
```

This is the final model (re-estimated with REML=TRUE):

```
model <- lmer(FSQ ~ Timefac*Groupcen + (1 + Timecen | Subject),
              data = dat, REML = TRUE)

summary(model)
```

```
## Linear mixed model fit by REML ['lmerMod']
## Formula: FSQ ~ Timefac * Groupcen + (1 + Timecen | Subject)
##    Data: dat
##
## REML criterion at convergence: 1167.5
##
## Scaled residuals:
##      Min       1Q   Median       3Q      Max
## -1.91326 -0.51461  0.05832  0.51931  2.68433
##
## Random effects:
##  Groups   Name        Variance Std.Dev. Corr
##  Subject  (Intercept) 165.189  12.853
##           Timecen       5.446   2.334   0.61
##  Residual              76.179   8.728
## Number of obs: 152, groups:  Subject, 38
##
## Fixed effects:
##                   Estimate Std. Error t value
## (Intercept)        68.8697     2.1943  31.386
## Timefac2          -30.0406     2.1527 -13.955
## Timefac3          -31.0364     2.5245 -12.294
## Timefac4          -40.6408     3.0449 -13.347
## Groupcen            1.9874     2.1943   0.906
## Timefac2:Groupcen   1.1359     2.1527   0.528
## Timefac3:Groupcen   0.8459     2.5245   0.335
## Timefac4:Groupcen   2.2122     3.0449   0.727
##
## Correlation of Fixed Effects:
##             (Intr) Timfc2 Timfc3 Timfc4 Gropcn Tmf2:G Tmf3:G
## Timefac2    -0.408
## Timefac3    -0.329  0.586
## Timefac4    -0.258  0.575  0.716
## Groupcen    -0.105  0.043  0.035  0.027
## Tmfc2:Grpcn  0.043 -0.105 -0.062 -0.060 -0.408
## Tmfc3:Grpcn  0.035 -0.062 -0.105 -0.075 -0.329  0.586
## Tmfc4:Grpcn  0.027 -0.060 -0.075 -0.105 -0.258  0.575  0.716
```

##### Assumptions

Here are the residuals plotted against fitted values to investigate underlying heteroskedasticity and linearity assumptions:

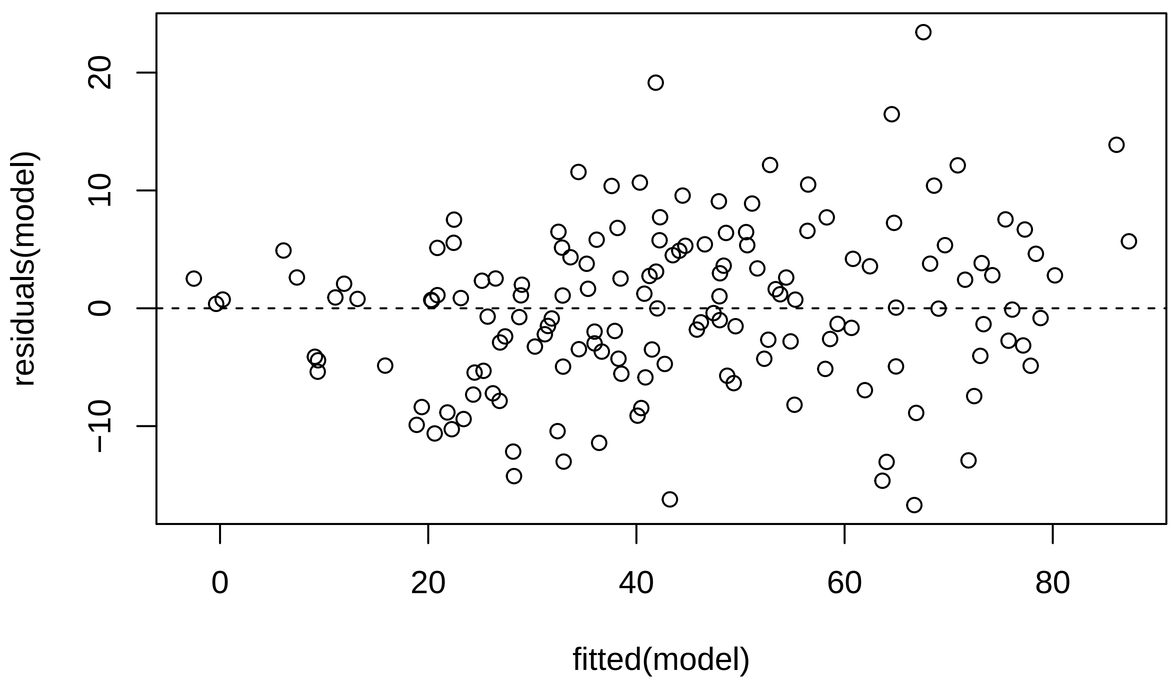

And the assumption of normality of residuals with histogram and qqplot:

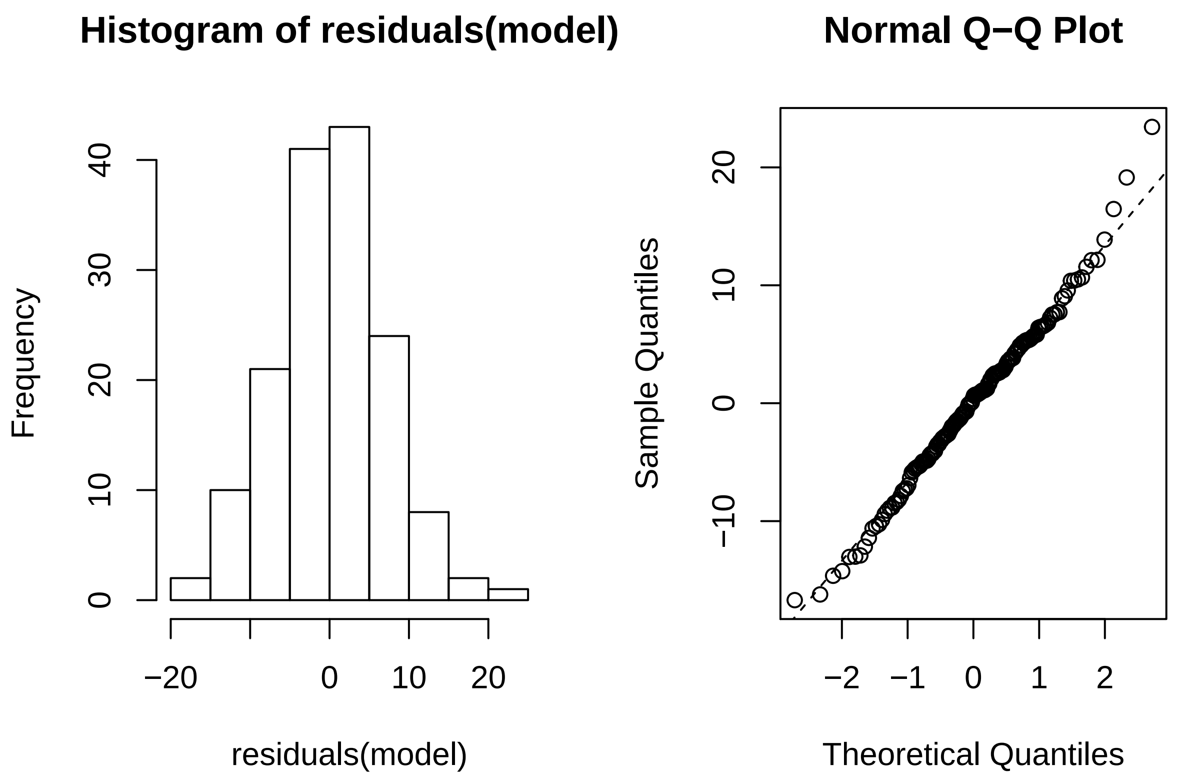

Using *Influence.ME*, outlying subjects are checked:

```
outlier <- influence(model, group = "Subject")
cks.d <- cooks.distance(outlier)
dfbetas <- dfbetas(outlier)
```

Now the Cook's distance values will be plotted. Outlier Subjects are highlighted in red:

```
plot(outlier,
     which = "cook",
     sort = TRUE,
     cutoff = 4/length(unique(dat$Subject)),
     xlab = "Cooks Distance")
```

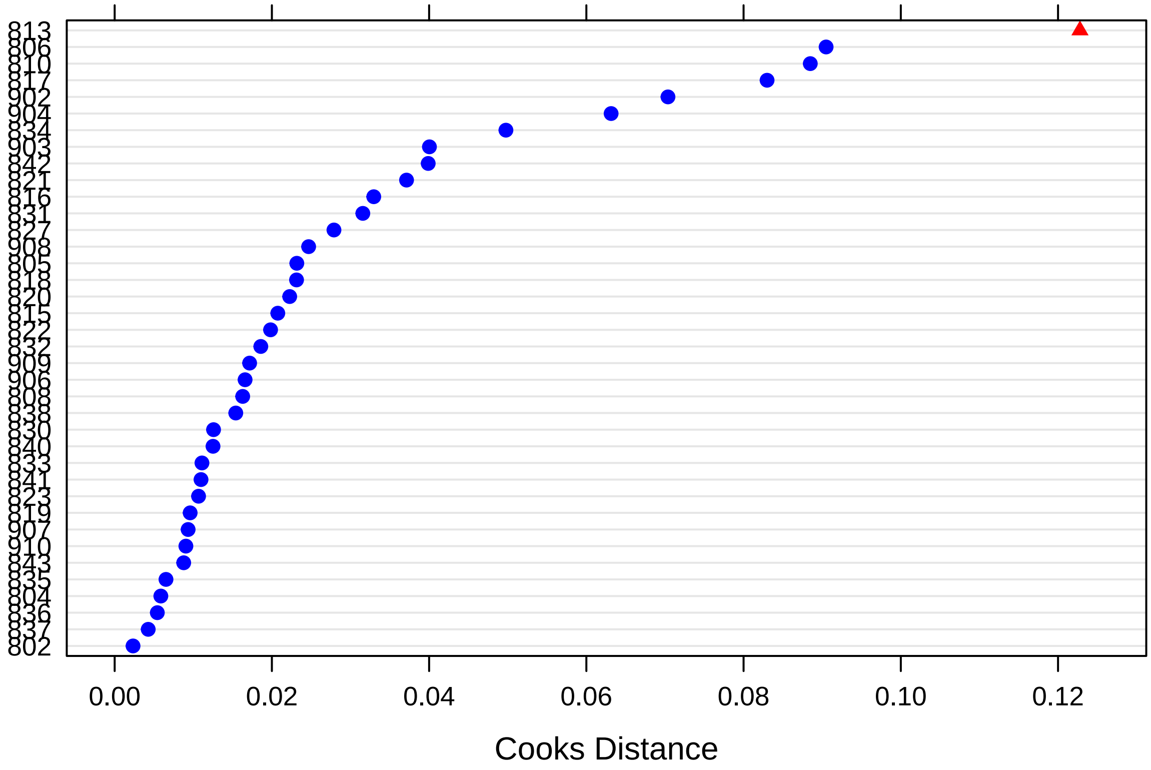

There is one outlying subject based on Cook's distance (i.e., *813*).

Finally, these are the p-values for dfbeta values, that indicate whether exclusion of a specific subject suffices to change significance values:

```
sigtest(outlier, test = -1.96)
```

```
## $Intercept
##     Altered.Teststat Altered.Sig Changed.Sig
## 802         30.58633       FALSE       FALSE
## 804         30.35367       FALSE       FALSE
## 805         30.59706       FALSE       FALSE
## 806         31.33866       FALSE       FALSE
## 808         30.39204       FALSE       FALSE
## 810         34.82221       FALSE       FALSE
## 813         30.59868       FALSE       FALSE
## 815         30.63145       FALSE       FALSE
## 816         31.92031       FALSE       FALSE
## 817         30.90671       FALSE       FALSE
## 818         30.88005       FALSE       FALSE
## 819         30.66517       FALSE       FALSE
## 820         31.12820       FALSE       FALSE
## 821         30.51864       FALSE       FALSE
## 822         30.71148       FALSE       FALSE
## 823         30.70353       FALSE       FALSE
## 827         30.82114       FALSE       FALSE
## 830         30.72543       FALSE       FALSE
```

```
## 831          30.62104       FALSE       FALSE
## 832          30.89149       FALSE       FALSE
## 833          31.16118       FALSE       FALSE
## 834          32.21924       FALSE       FALSE
## 835          30.59080       FALSE       FALSE
## 836          30.40384       FALSE       FALSE
## 837          30.60330       FALSE       FALSE
## 838          30.63021       FALSE       FALSE
## 840          30.69144       FALSE       FALSE
## 841          30.98845       FALSE       FALSE
## 842          31.66438       FALSE       FALSE
## 843          30.68458       FALSE       FALSE
## 902          30.42929       FALSE       FALSE
## 903          31.30420       FALSE       FALSE
## 904          30.90942       FALSE       FALSE
## 906          31.44241       FALSE       FALSE
## 907          30.55132       FALSE       FALSE
## 908          30.96243       FALSE       FALSE
## 909          30.90467       FALSE       FALSE
## 910          31.26417       FALSE       FALSE
##
## $Timefac2
##     Altered.Teststat Altered.Sig Changed.Sig
## 802        -13.59228        TRUE       FALSE
## 804        -13.47898        TRUE       FALSE
## 805        -13.58474        TRUE       FALSE
## 806        -13.84423        TRUE       FALSE
## 808        -13.46939        TRUE       FALSE
## 810        -13.86794        TRUE       FALSE
## 813        -13.82051        TRUE       FALSE
## 815        -13.80864        TRUE       FALSE
## 816        -13.66017        TRUE       FALSE
## 817        -14.80605        TRUE       FALSE
## 818        -13.64245        TRUE       FALSE
## 819        -13.59860        TRUE       FALSE
## 820        -13.61687        TRUE       FALSE
## 821        -13.97869        TRUE       FALSE
## 822        -13.90588        TRUE       FALSE
## 823        -13.76010        TRUE       FALSE
## 827        -13.67391        TRUE       FALSE
## 830        -13.82390        TRUE       FALSE
## 831        -13.42073        TRUE       FALSE
## 832        -13.57935        TRUE       FALSE
## 833        -13.60809        TRUE       FALSE
## 834        -13.63595        TRUE       FALSE
## 835        -13.72527        TRUE       FALSE
## 836        -13.51705        TRUE       FALSE
## 837        -13.64324        TRUE       FALSE
## 838        -13.76937        TRUE       FALSE
## 840        -13.64466        TRUE       FALSE
## 841        -13.66111        TRUE       FALSE
## 842        -14.00972        TRUE       FALSE
## 843        -13.78299        TRUE       FALSE
## 902        -13.66412        TRUE       FALSE
```

```
## 903        -14.35365        TRUE       FALSE
## 904        -14.56587        TRUE       FALSE
## 906        -13.65428        TRUE       FALSE
## 907        -13.68410        TRUE       FALSE
## 908        -13.64755        TRUE       FALSE
## 909        -14.02670        TRUE       FALSE
## 910        -13.72341        TRUE       FALSE
##
## $Timefac3
##     Altered.Teststat Altered.Sig Changed.Sig
## 802        -11.97107        TRUE       FALSE
## 804        -11.81938        TRUE       FALSE
## 805        -11.99267        TRUE       FALSE
## 806        -12.40752        TRUE       FALSE
## 808        -11.78975        TRUE       FALSE
## 810        -12.12700        TRUE       FALSE
## 813        -12.28963        TRUE       FALSE
## 815        -12.20557        TRUE       FALSE
## 816        -12.30304        TRUE       FALSE
## 817        -12.52185        TRUE       FALSE
## 818        -12.16343        TRUE       FALSE
## 819        -12.05877        TRUE       FALSE
## 820        -12.01263        TRUE       FALSE
## 821        -12.64752        TRUE       FALSE
## 822        -12.39148        TRUE       FALSE
## 823        -11.97413        TRUE       FALSE
## 827        -12.06852        TRUE       FALSE
## 830        -12.18633        TRUE       FALSE
## 831        -11.84848        TRUE       FALSE
## 832        -12.04695        TRUE       FALSE
## 833        -11.92143        TRUE       FALSE
## 834        -12.15779        TRUE       FALSE
## 835        -12.02238        TRUE       FALSE
## 836        -11.81743        TRUE       FALSE
## 837        -11.92140        TRUE       FALSE
## 838        -12.25885        TRUE       FALSE
## 840        -11.93430        TRUE       FALSE
## 841        -12.04529        TRUE       FALSE
## 842        -12.82148        TRUE       FALSE
## 843        -12.00082        TRUE       FALSE
## 902        -11.90931        TRUE       FALSE
## 903        -12.36553        TRUE       FALSE
## 904        -12.78667        TRUE       FALSE
## 906        -12.02006        TRUE       FALSE
## 907        -12.04951        TRUE       FALSE
## 908        -11.91898        TRUE       FALSE
## 909        -12.13448        TRUE       FALSE
## 910        -12.05929        TRUE       FALSE
##
## $Timefac4
##     Altered.Teststat Altered.Sig Changed.Sig
## 802        -13.03877        TRUE       FALSE
## 804        -12.88957        TRUE       FALSE
## 805        -13.00093        TRUE       FALSE
```

```
## 806        -13.84910        TRUE       FALSE
## 808        -12.93107        TRUE       FALSE
## 810        -13.18784        TRUE       FALSE
## 813        -13.63003        TRUE       FALSE
## 815        -13.26878        TRUE       FALSE
## 816        -13.47572        TRUE       FALSE
## 817        -13.03856        TRUE       FALSE
## 818        -13.18885        TRUE       FALSE
## 819        -13.12493        TRUE       FALSE
## 820        -13.03670        TRUE       FALSE
## 821        -13.65771        TRUE       FALSE
## 822        -13.33373        TRUE       FALSE
## 823        -13.10197        TRUE       FALSE
## 827        -13.03125        TRUE       FALSE
## 830        -13.03010        TRUE       FALSE
## 831        -12.96478        TRUE       FALSE
## 832        -12.86813        TRUE       FALSE
## 833        -12.99912        TRUE       FALSE
## 834        -13.10495        TRUE       FALSE
## 835        -13.10525        TRUE       FALSE
## 836        -12.91350        TRUE       FALSE
## 837        -13.00753        TRUE       FALSE
## 838        -13.48245        TRUE       FALSE
## 840        -13.04240        TRUE       FALSE
## 841        -12.98018        TRUE       FALSE
## 842        -14.21617        TRUE       FALSE
## 843        -13.08621        TRUE       FALSE
## 902        -13.05995        TRUE       FALSE
## 903        -13.09784        TRUE       FALSE
## 904        -13.60558        TRUE       FALSE
## 906        -12.98822        TRUE       FALSE
## 907        -12.96298        TRUE       FALSE
## 908        -13.08396        TRUE       FALSE
## 909        -13.07079        TRUE       FALSE
## 910        -13.09544        TRUE       FALSE
##
## $Groupcen
##     Altered.Teststat Altered.Sig Changed.Sig
## 802        0.8162552       FALSE       FALSE
## 804        0.9913819       FALSE       FALSE
## 805        0.7948564       FALSE       FALSE
## 806        1.2910443       FALSE       FALSE
## 808        0.9498130       FALSE       FALSE
## 810        0.3680433       FALSE       FALSE
## 813        1.2313271       FALSE       FALSE
## 815        0.9860549       FALSE       FALSE
## 816        0.6274969       FALSE       FALSE
## 817        0.7785568       FALSE       FALSE
## 818        0.7583411       FALSE       FALSE
## 819        0.7530643       FALSE       FALSE
## 820        0.9514823       FALSE       FALSE
## 821        0.8538895       FALSE       FALSE
## 822        0.8630324       FALSE       FALSE
## 823        0.9061089       FALSE       FALSE
```

```
## 827        1.0356672       FALSE       FALSE
## 830        0.8634244       FALSE       FALSE
## 831        0.8710319       FALSE       FALSE
## 832        0.7352744       FALSE       FALSE
## 833        1.0289225       FALSE       FALSE
## 834        0.5952103       FALSE       FALSE
## 835        0.7847851       FALSE       FALSE
## 836        0.9501817       FALSE       FALSE
## 837        0.8491830       FALSE       FALSE
## 838        0.8065758       FALSE       FALSE
## 840        0.7427898       FALSE       FALSE
## 841        0.7610031       FALSE       FALSE
## 842        1.1338195       FALSE       FALSE
## 843        0.8297389       FALSE       FALSE
## 902        1.0799538       FALSE       FALSE
## 903        1.1426634       FALSE       FALSE
## 904        0.6502013       FALSE       FALSE
## 906        1.1040033       FALSE       FALSE
## 907        0.9547908       FALSE       FALSE
## 908        0.9572881       FALSE       FALSE
## 909        1.0312503       FALSE       FALSE
## 910        1.0759690       FALSE       FALSE
##
## $`Timefac2:Groupcen`
##     Altered.Teststat Altered.Sig Changed.Sig
## 802       0.55036228       FALSE       FALSE
## 804       0.40323175       FALSE       FALSE
## 805       0.81436423       FALSE       FALSE
## 806       0.56733065       FALSE       FALSE
## 808       0.32020022       FALSE       FALSE
## 810       0.74518119       FALSE       FALSE
## 813      -0.14305760       FALSE       FALSE
## 815       0.71520450       FALSE       FALSE
## 816       0.55978762       FALSE       FALSE
## 817       1.03509087       FALSE       FALSE
## 818       0.65979198       FALSE       FALSE
## 819       0.58612351       FALSE       FALSE
## 820       0.71881252       FALSE       FALSE
## 821       0.71041042       FALSE       FALSE
## 822       0.39600021       FALSE       FALSE
## 823       0.49767701       FALSE       FALSE
## 827       0.50573812       FALSE       FALSE
## 830       0.44072287       FALSE       FALSE
## 831       0.20747740       FALSE       FALSE
## 832       0.44755215       FALSE       FALSE
## 833       0.53919872       FALSE       FALSE
## 834       0.74409986       FALSE       FALSE
## 835       0.65730066       FALSE       FALSE
## 836       0.45915080       FALSE       FALSE
## 837       0.52877662       FALSE       FALSE
## 838       0.43898452       FALSE       FALSE
## 840       0.60002384       FALSE       FALSE
## 841       0.50586818       FALSE       FALSE
## 842       0.33978545       FALSE       FALSE
```

```
## 843       0.42766021       FALSE       FALSE
## 902       0.03718222       FALSE       FALSE
## 903       0.22835976       FALSE       FALSE
## 904       1.00466282       FALSE       FALSE
## 906       0.50561517       FALSE       FALSE
## 907       0.61497021       FALSE       FALSE
## 908       0.66003875       FALSE       FALSE
## 909       0.30488418       FALSE       FALSE
## 910       0.42581153       FALSE       FALSE
##
## $`Timefac3:Groupcen`
##     Altered.Teststat Altered.Sig Changed.Sig
## 802       0.35418043       FALSE       FALSE
## 804       0.20382186       FALSE       FALSE
## 805       0.50618058       FALSE       FALSE
## 806       0.43321851       FALSE       FALSE
## 808       0.07219511       FALSE       FALSE
## 810       0.44675998       FALSE       FALSE
## 813      -0.27370827       FALSE       FALSE
## 815       0.50796488       FALSE       FALSE
## 816       0.52370644       FALSE       FALSE
## 817       0.61561700       FALSE       FALSE
## 818       0.44128100       FALSE       FALSE
## 819       0.30685933       FALSE       FALSE
## 820       0.39547973       FALSE       FALSE
## 821       0.72738224       FALSE       FALSE
## 822       0.09445079       FALSE       FALSE
## 823       0.40423814       FALSE       FALSE
## 827       0.24420301       FALSE       FALSE
## 830       0.19067894       FALSE       FALSE
## 831       0.04827568       FALSE       FALSE
## 832       0.39735404       FALSE       FALSE
## 833       0.42247126       FALSE       FALSE
## 834       0.43084744       FALSE       FALSE
## 835       0.38489497       FALSE       FALSE
## 836       0.18011224       FALSE       FALSE
## 837       0.43250049       FALSE       FALSE
## 838       0.23160350       FALSE       FALSE
## 840       0.43296867       FALSE       FALSE
## 841       0.29659230       FALSE       FALSE
## 842       0.02666819       FALSE       FALSE
## 843       0.35506064       FALSE       FALSE
## 902      -0.12516140       FALSE       FALSE
## 903       0.14367113       FALSE       FALSE
## 904       0.78222939       FALSE       FALSE
## 906       0.33567956       FALSE       FALSE
## 907       0.43232348       FALSE       FALSE
## 908       0.57454074       FALSE       FALSE
## 909       0.26889230       FALSE       FALSE
## 910       0.27711396       FALSE       FALSE
##
## $`Timefac4:Groupcen`
##     Altered.Teststat Altered.Sig Changed.Sig
## 802        0.6892263       FALSE       FALSE
```

```
## 804        0.6579068       FALSE       FALSE
## 805        0.8747285       FALSE       FALSE
## 806        1.0765011       FALSE       FALSE
## 808        0.5597835       FALSE       FALSE
## 810        0.8680257       FALSE       FALSE
## 813        0.2571118       FALSE       FALSE
## 815        0.8921578       FALSE       FALSE
## 816        0.9440935       FALSE       FALSE
## 817        0.7649609       FALSE       FALSE
## 818        0.9844461       FALSE       FALSE
## 819        0.6514717       FALSE       FALSE
## 820        0.7059995       FALSE       FALSE
## 821        0.9855926       FALSE       FALSE
## 822        0.5936087       FALSE       FALSE
## 823        0.6419160       FALSE       FALSE
## 827        0.4663662       FALSE       FALSE
## 830        0.7056418       FALSE       FALSE
## 831        0.4835601       FALSE       FALSE
## 832        0.6142594       FALSE       FALSE
## 833        0.7632094       FALSE       FALSE
## 834        0.9693596       FALSE       FALSE
## 835        0.8252750       FALSE       FALSE
## 836        0.6259403       FALSE       FALSE
## 837        0.7552026       FALSE       FALSE
## 838        0.5319107       FALSE       FALSE
## 840        0.6894183       FALSE       FALSE
## 841        0.7620977       FALSE       FALSE
## 842        0.4364612       FALSE       FALSE
## 843        0.6495498       FALSE       FALSE
## 902        0.4476150       FALSE       FALSE
## 903        0.6585508       FALSE       FALSE
## 904        1.0008516       FALSE       FALSE
## 906        0.7880958       FALSE       FALSE
## 907        0.7230489       FALSE       FALSE
## 908        0.8542793       FALSE       FALSE
## 909        0.6656070       FALSE       FALSE
## 910        0.6500076       FALSE       FALSE
```

There is no indication that results are changed by an outlying subject.

#### BAT

The base model is now tested against the same models without the effects of time:

```
model <- lmer(BAT_speed ~ Sessionfac*Groupcen + (1 + Sessioncen | Subject),
              data = dat, REML = FALSE)

model_null <- lmer(BAT_speed ~ Sessionfac + (1 + Sessioncen | Subject),
              data = dat, REML = FALSE)
```

These are the results, which **do not** support rejection of null hypothesis:

```
anova(model, model_null)
```

```
## Data: dat
```

```
## Models:
## model_null: BAT_speed ~ Sessionfac + (1 + Sessioncen | Subject)
## model: BAT_speed ~ Sessionfac * Groupcen + (1 + Sessioncen | Subject)
##            Df     AIC     BIC logLik deviance  Chisq Chi Df Pr(>Chisq)
## model_null  7 -23.451 -4.4847 18.726  -37.451
## model      10 -21.923  5.1725 20.961  -41.923 4.4714      3     0.2149
```

This is the final model (re-estimated with REML=TRUE):

```
model <- lmer(BAT_speed ~ Sessionfac*Groupcen + (1 + Sessioncen | Subject),
              data = dat, REML = TRUE)

summary(model)
```

```
## Linear mixed model fit by REML ['lmerMod']
## Formula: BAT_speed ~ Sessionfac * Groupcen + (1 + Sessioncen | Subject)
##    Data: dat
##
## REML criterion at convergence: -12.1
##
## Scaled residuals:
##      Min       1Q   Median       3Q      Max
## -2.34594 -0.43577 -0.09124  0.41218  2.32819
##
## Random effects:
##  Groups   Name        Variance Std.Dev. Corr
##  Subject  (Intercept) 0.05179  0.2276
##           Sessioncen  0.01610  0.1269   0.81
##  Residual             0.01513  0.1230
## Number of obs: 111, groups:  Subject, 38
##
## Fixed effects:
##                      Estimate Std. Error t value
## (Intercept)           0.21328    0.03214   6.636
## Sessionfac2           0.16000    0.03599   4.445
## Sessionfac3           0.35969    0.05080   7.080
## Groupcen              0.04493    0.03214   1.398
## Sessionfac2:Groupcen -0.05873    0.03599  -1.632
## Sessionfac3:Groupcen -0.08077    0.05080  -1.590
##
## Correlation of Fixed Effects:
##             (Intr) Sssnf2 Sssnf3 Gropcn Sss2:G
## Sessionfac2 -0.237
## Sessionfac3 -0.051  0.723
## Groupcen    -0.131  0.048  0.022
## Sssnfc2:Grp  0.048 -0.126 -0.090 -0.237
## Sssnfc3:Grp  0.022 -0.090 -0.115 -0.051  0.723
```

##### Assumptions

Here are the residuals plotted against fitted values to investigate underlying heteroskedasticity and linearity assumptions:

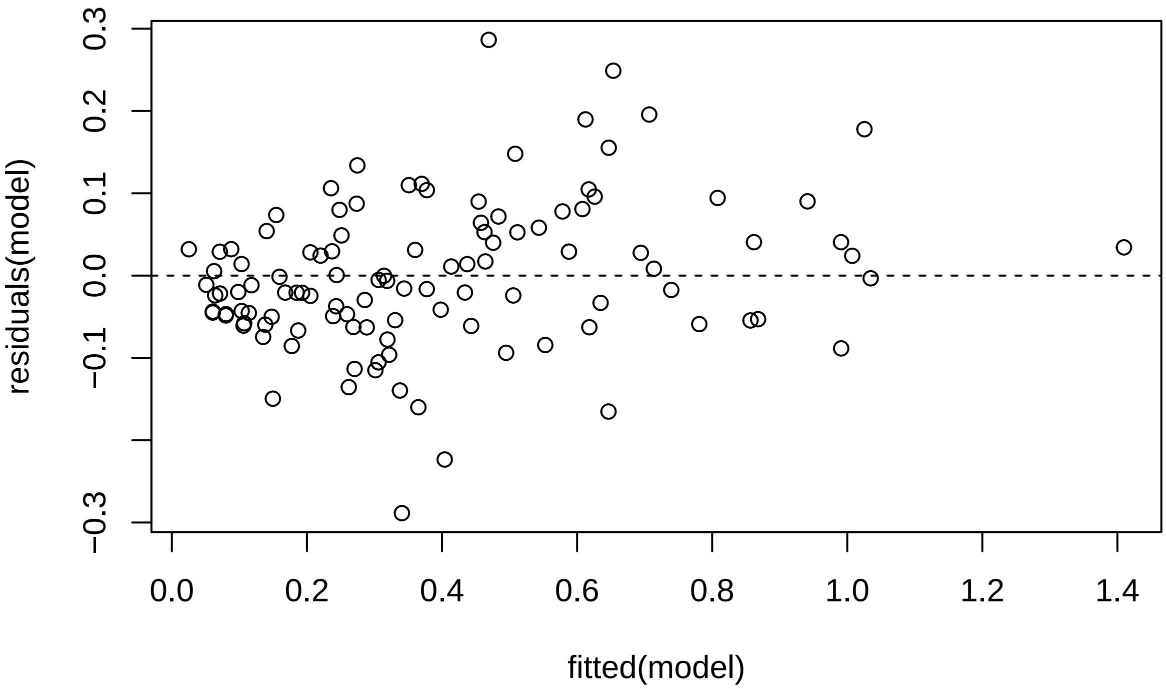

And the assumption of normality of residuals with histogram and qqplot:

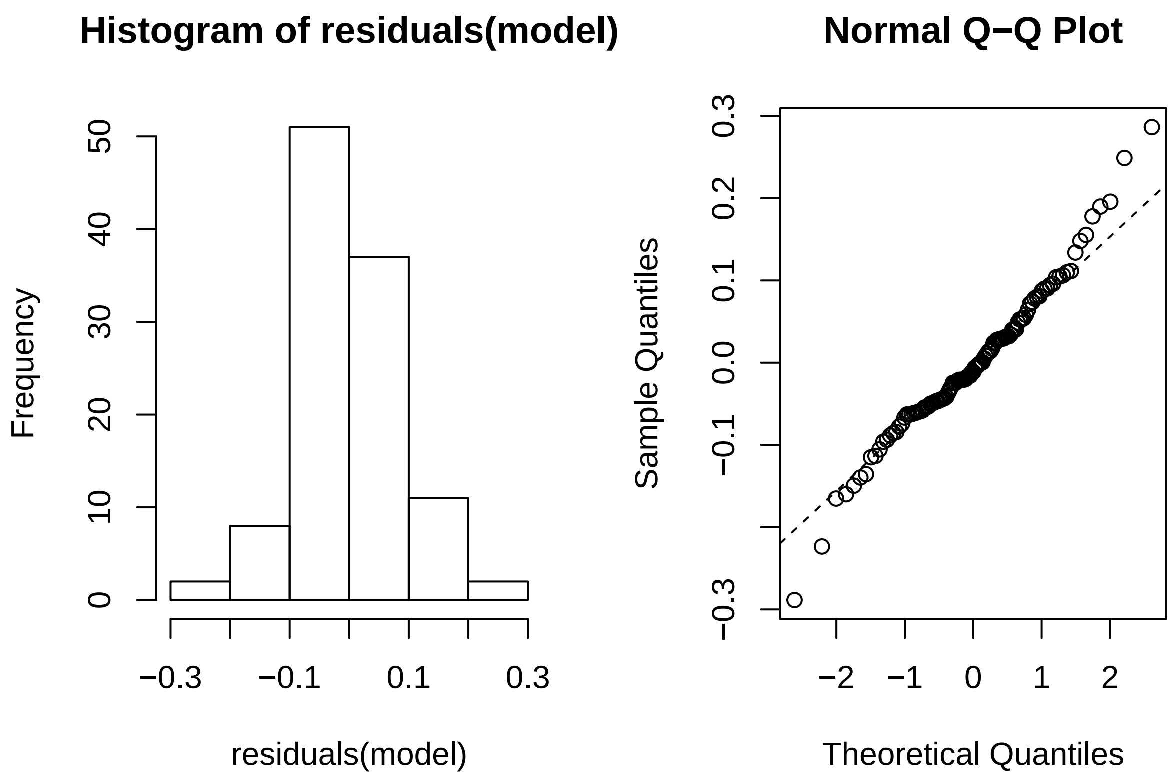

Using *Influence.ME*, outlying subjects are checked:

```
outlier <- influence(model, group = "Subject")
cks.d <- cooks.distance(outlier)
dfbetas <- dfbetas(outlier)
```

Now the Cook's distance values will be plotted. Outlier Subjects are highlighted in red:

```
plot(outlier,
     which = "cook",
     sort = TRUE,
     cutoff = 4/length(unique(dat$Subject)),
     xlab = "Cooks Distance")
```

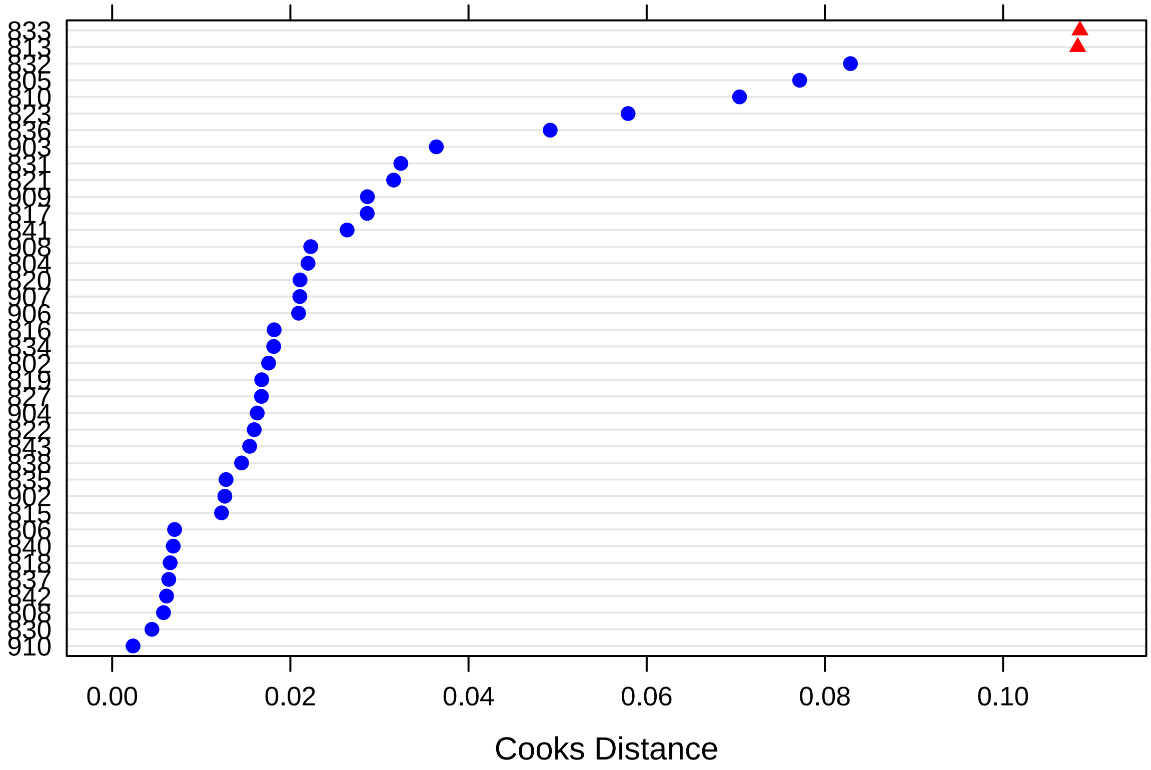

There are no outlying subjects.

Finally, these are the p-values for dfbeta values, that indicate whether exclusion of a specific subject suffices to change significance values:

```
sigtest(outlier, test = -1.96)
```

```
## $Intercept
##     Altered.Teststat Altered.Sig Changed.Sig
## 802         6.684488       FALSE       FALSE
## 804         6.424265       FALSE       FALSE
## 805         6.854502       FALSE       FALSE
## 806         6.366277       FALSE       FALSE
## 808         6.470363       FALSE       FALSE
## 810         6.428394       FALSE       FALSE
## 813         6.834550       FALSE       FALSE
## 815         6.484903       FALSE       FALSE
## 816         6.436037       FALSE       FALSE
## 817         6.529465       FALSE       FALSE
## 818         6.497136       FALSE       FALSE
## 819         6.453251       FALSE       FALSE
## 820         6.358061       FALSE       FALSE
## 821         6.405795       FALSE       FALSE
## 822         6.684425       FALSE       FALSE
## 823         6.396853       FALSE       FALSE
## 827         6.765617       FALSE       FALSE
## 830         6.591231       FALSE       FALSE
```

```
## 831          6.292783       FALSE       FALSE
## 832          6.445720       FALSE       FALSE
## 833          6.842075       FALSE       FALSE
## 834          6.807644       FALSE       FALSE
## 835          6.526421       FALSE       FALSE
## 836          6.499382       FALSE       FALSE
## 837          6.427573       FALSE       FALSE
## 838          6.442610       FALSE       FALSE
## 840          6.673670       FALSE       FALSE
## 841          6.790218       FALSE       FALSE
## 842          6.474158       FALSE       FALSE
## 843          6.734369       FALSE       FALSE
## 902          6.559275       FALSE       FALSE
## 903          6.527170       FALSE       FALSE
## 904          6.380165       FALSE       FALSE
## 906          6.544639       FALSE       FALSE
## 907          6.709557       FALSE       FALSE
## 908          6.452180       FALSE       FALSE
## 909          6.562189       FALSE       FALSE
## 910          6.523218       FALSE       FALSE
##
## $Sessionfac2
##     Altered.Teststat Altered.Sig Changed.Sig
## 802          4.460162       FALSE       FALSE
## 804          4.153355       FALSE       FALSE
## 805          4.559989       FALSE       FALSE
## 806          4.471921       FALSE       FALSE
## 808          4.428627       FALSE       FALSE
## 810          4.119432       FALSE       FALSE
## 813          4.851007       FALSE       FALSE
## 815          4.508932       FALSE       FALSE
## 816          4.341178       FALSE       FALSE
## 817          4.587129       FALSE       FALSE
## 818          4.228517       FALSE       FALSE
## 819          4.448817       FALSE       FALSE
## 820          4.637993       FALSE       FALSE
## 821          4.302764       FALSE       FALSE
## 822          4.397615       FALSE       FALSE
## 823          4.549171       FALSE       FALSE
## 827          4.546492       FALSE       FALSE
## 830          4.379663       FALSE       FALSE
## 831          4.644154       FALSE       FALSE
## 832          4.120168       FALSE       FALSE
## 833          4.650552       FALSE       FALSE
## 834          4.409629       FALSE       FALSE
## 835          4.443134       FALSE       FALSE
## 836          4.040628       FALSE       FALSE
## 837          4.286348       FALSE       FALSE
## 838          4.404574       FALSE       FALSE
## 840          4.342152       FALSE       FALSE
## 841          4.222587       FALSE       FALSE
## 842          4.414580       FALSE       FALSE
## 843          4.525235       FALSE       FALSE
## 902          4.248675       FALSE       FALSE
```

```
## 903         4.179340       FALSE       FALSE
## 904         4.327107       FALSE       FALSE
## 906         4.191683       FALSE       FALSE
## 907         4.371820       FALSE       FALSE
## 908         4.274185       FALSE       FALSE
## 909         4.271312       FALSE       FALSE
## 910         4.299180       FALSE       FALSE
##
## $Sessionfac3
##     Altered.Teststat Altered.Sig Changed.Sig
## 802         7.166600       FALSE       FALSE
## 804         6.807680       FALSE       FALSE
## 805         6.974188       FALSE       FALSE
## 806         7.051638       FALSE       FALSE
## 808         6.932641       FALSE       FALSE
## 810         6.775293       FALSE       FALSE
## 813         6.854568       FALSE       FALSE
## 815         7.065248       FALSE       FALSE
## 816         6.775879       FALSE       FALSE
## 817         7.370667       FALSE       FALSE
## 818         6.843901       FALSE       FALSE
## 819         7.225794       FALSE       FALSE
## 820         7.126536       FALSE       FALSE
## 821         6.919370       FALSE       FALSE
## 822         7.100884       FALSE       FALSE
## 823         7.033819       FALSE       FALSE
## 827         7.108405       FALSE       FALSE
## 830         6.965285       FALSE       FALSE
## 831         6.881994       FALSE       FALSE
## 832         6.798390       FALSE       FALSE
## 833         7.439619       FALSE       FALSE
## 834         7.027835       FALSE       FALSE
## 835         7.130193       FALSE       FALSE
## 836         6.808818       FALSE       FALSE
## 837         6.879642       FALSE       FALSE
## 838         7.014817       FALSE       FALSE
## 840         6.926722       FALSE       FALSE
## 841         6.857742       FALSE       FALSE
## 842         7.092661       FALSE       FALSE
## 843         7.234555       FALSE       FALSE
## 902         6.927354       FALSE       FALSE
## 903         6.976102       FALSE       FALSE
## 904         7.032671       FALSE       FALSE
## 906         6.896614       FALSE       FALSE
## 907         6.799901       FALSE       FALSE
## 908         6.841171       FALSE       FALSE
## 909         6.909646       FALSE       FALSE
## 910         6.853525       FALSE       FALSE
##
## $Groupcen
##     Altered.Teststat Altered.Sig Changed.Sig
## 802        1.5178162       FALSE       FALSE
## 804        1.3829709       FALSE       FALSE
## 805        1.5373824       FALSE       FALSE
```

```
## 806         1.3873876       FALSE       FALSE
## 808         1.3123360       FALSE       FALSE
## 810         1.4731215       FALSE       FALSE
## 813         1.3332150       FALSE       FALSE
## 815         1.3010526       FALSE       FALSE
## 816         1.3475539       FALSE       FALSE
## 817         1.2598210       FALSE       FALSE
## 818         1.3931676       FALSE       FALSE
## 819         1.3272511       FALSE       FALSE
## 820         1.2502010       FALSE       FALSE
## 821         1.7012592       FALSE       FALSE
## 822         1.5279184       FALSE       FALSE
## 823         1.3252128       FALSE       FALSE
## 827         1.2601584       FALSE       FALSE
## 830         1.4553627       FALSE       FALSE
## 831         1.4514787       FALSE       FALSE
## 832         1.7812350       FALSE       FALSE
## 833         0.9888774       FALSE       FALSE
## 834         1.5691771       FALSE       FALSE
## 835         1.2628547       FALSE       FALSE
## 836         1.3010963       FALSE       FALSE
## 837         1.2690984       FALSE       FALSE
## 838         1.1966864       FALSE       FALSE
## 840         1.5008863       FALSE       FALSE
## 841         1.6039476       FALSE       FALSE
## 842         1.3702848       FALSE       FALSE
## 843         1.5008098       FALSE       FALSE
## 902         1.2387296       FALSE       FALSE
## 903         1.4180253       FALSE       FALSE
## 904         1.4279143       FALSE       FALSE
## 906         1.4229085       FALSE       FALSE
## 907         1.2150267       FALSE       FALSE
## 908         1.1855141       FALSE       FALSE
## 909         1.1738423       FALSE       FALSE
## 910         1.4051725       FALSE       FALSE
##
## $`Sessionfac2:Groupcen`
##     Altered.Teststat Altered.Sig Changed.Sig
## 802        -1.509119       FALSE       FALSE
## 804        -1.393799       FALSE       FALSE
## 805        -1.657216       FALSE       FALSE
## 806        -1.732246       FALSE       FALSE
## 808        -1.698517       FALSE       FALSE
## 810        -1.285123       FALSE       FALSE
## 813        -1.920877       FALSE       FALSE
## 815        -1.767148       FALSE       FALSE
## 816        -1.580271       FALSE       FALSE
## 817        -1.814327       FALSE       FALSE
## 818        -1.732887       FALSE       FALSE
## 819        -1.555431       FALSE       FALSE
## 820        -1.402581       FALSE       FALSE
## 821        -1.580800       FALSE       FALSE
## 822        -1.573762       FALSE       FALSE
## 823        -1.723391       FALSE       FALSE
```

```
## 827        -1.771024       FALSE       FALSE
## 830        -1.546661       FALSE       FALSE
## 831        -1.863481       FALSE       FALSE
## 832        -1.361555       FALSE       FALSE
## 833        -1.283299       FALSE       FALSE
## 834        -1.515689       FALSE       FALSE
## 835        -1.700460       FALSE       FALSE
## 836        -1.218723       FALSE       FALSE
## 837        -1.637087       FALSE       FALSE
## 838        -1.506693       FALSE       FALSE
## 840        -1.572201       FALSE       FALSE
## 841        -1.793337       FALSE       FALSE
## 842        -1.543430       FALSE       FALSE
## 843        -1.500839       FALSE       FALSE
## 902        -1.506724       FALSE       FALSE
## 903        -1.943189       FALSE       FALSE
## 904        -1.576376       FALSE       FALSE
## 906        -1.837718       FALSE       FALSE
## 907        -1.637787       FALSE       FALSE
## 908        -1.655818       FALSE       FALSE
## 909        -1.625648       FALSE       FALSE
## 910        -1.629539       FALSE       FALSE
##
## $`Sessionfac3:Groupcen`
##     Altered.Teststat Altered.Sig Changed.Sig
## 802        -1.417366       FALSE       FALSE
## 804        -1.488617       FALSE       FALSE
## 805        -1.890247       FALSE       FALSE
## 806        -1.692839       FALSE       FALSE
## 808        -1.609194       FALSE       FALSE
## 810        -1.433696       FALSE       FALSE
## 813        -1.405287       FALSE       FALSE
## 815        -1.703332       FALSE       FALSE
## 816        -1.379605       FALSE       FALSE
## 817        -1.874428       FALSE       FALSE
## 818        -1.668906       FALSE       FALSE
## 819        -1.404673       FALSE       FALSE
## 820        -1.439627       FALSE       FALSE
## 821        -1.597825       FALSE       FALSE
## 822        -1.439039       FALSE       FALSE
## 823        -1.955720       FALSE       FALSE
## 827        -1.702057       FALSE       FALSE
## 830        -1.498261       FALSE       FALSE
## 831        -1.574442       FALSE       FALSE
## 832        -1.299433       FALSE       FALSE
## 833        -1.320905       FALSE       FALSE
## 834        -1.459391       FALSE       FALSE
## 835        -1.740482       FALSE       FALSE
## 836        -1.296399       FALSE       FALSE
## 837        -1.573205       FALSE       FALSE
## 838        -1.469159       FALSE       FALSE
## 840        -1.520086       FALSE       FALSE
## 841        -1.597366       FALSE       FALSE
## 842        -1.444039       FALSE       FALSE
```

```
## 843        -1.421250       FALSE       FALSE
## 902        -1.602734       FALSE       FALSE
## 903        -1.912859       FALSE       FALSE
## 904        -1.678197       FALSE       FALSE
## 906        -1.815345       FALSE       FALSE
## 907        -1.473356       FALSE       FALSE
## 908        -1.668418       FALSE       FALSE
## 909        -1.527843       FALSE       FALSE
## 910        -1.617685       FALSE       FALSE
```

There is no indication that results are changed by.

#### Implicit Fear Evaluation (EAST)

The base model is now tested against the same models without the effects of time:

```
model <- lmer(EAST ~ Sessionfac*Groupcen + (1 | Subject),
              data = dat, REML = FALSE)

model_null <- lmer(EAST ~ Sessionfac + Groupcen + (1 | Subject),
              data = dat, REML = FALSE)
```

These are the results, which **do not** support rejection of null hypothesis:

```
anova(model, model_null)
```

```
## Data: dat
## Models:
## model_null: EAST ~ Sessionfac + Groupcen + (1 | Subject)
## model: EAST ~ Sessionfac * Groupcen + (1 | Subject)
##            Df    AIC    BIC  logLik deviance  Chisq Chi Df Pr(>Chisq)
## model_null  6 1109.7 1126.1 -548.86   1097.7
## model       8 1113.5 1135.4 -548.78   1097.5 0.1713      2     0.9179
```

This is the final model (re-estimated with REML=TRUE):

```
model <- lmer(EAST ~ Sessionfac * Groupcen + (1 | Subject),
              data = dat, REML = TRUE)

summary(model)
```

```
## Linear mixed model fit by REML ['lmerMod']
## Formula: EAST ~ Sessionfac * Groupcen + (1 | Subject)
##    Data: dat
##
## REML criterion at convergence: 1066.9
##
## Scaled residuals:
##     Min      1Q  Median      3Q     Max
## -3.7689 -0.4063  0.0781  0.4618  2.9113
##
## Random effects:
##  Groups   Name        Variance Std.Dev.
##  Subject  (Intercept) 488.9    22.11
##  Residual             700.7    26.47
## Number of obs: 113, groups:  Subject, 38
```

```
##
## Fixed effects:
##                      Estimate Std. Error t value
## (Intercept)           -9.9433     5.6264  -1.767
## Sessionfac2            7.8235     6.1066   1.281
## Sessionfac3           10.6253     6.1743   1.721
## Groupcen              -7.4139     5.6264  -1.318
## Sessionfac2:Groupcen  -1.8235     6.1066  -0.299
## Sessionfac3:Groupcen   0.5414     6.1743   0.088
##
## Correlation of Fixed Effects:
##             (Intr) Sssnf2 Sssnf3 Gropcn Sss2:G
## Sessionfac2 -0.543
## Sessionfac3 -0.537  0.495
## Groupcen    -0.105  0.057  0.056
## Sssnfc2:Grp  0.057 -0.105 -0.052 -0.543
## Sssnfc3:Grp  0.056 -0.052 -0.125 -0.537  0.495
```

**Assumptions**

Here are the residuals plotted against fitted values to investigate underlying heteroskedasticity and linearity assumptions:

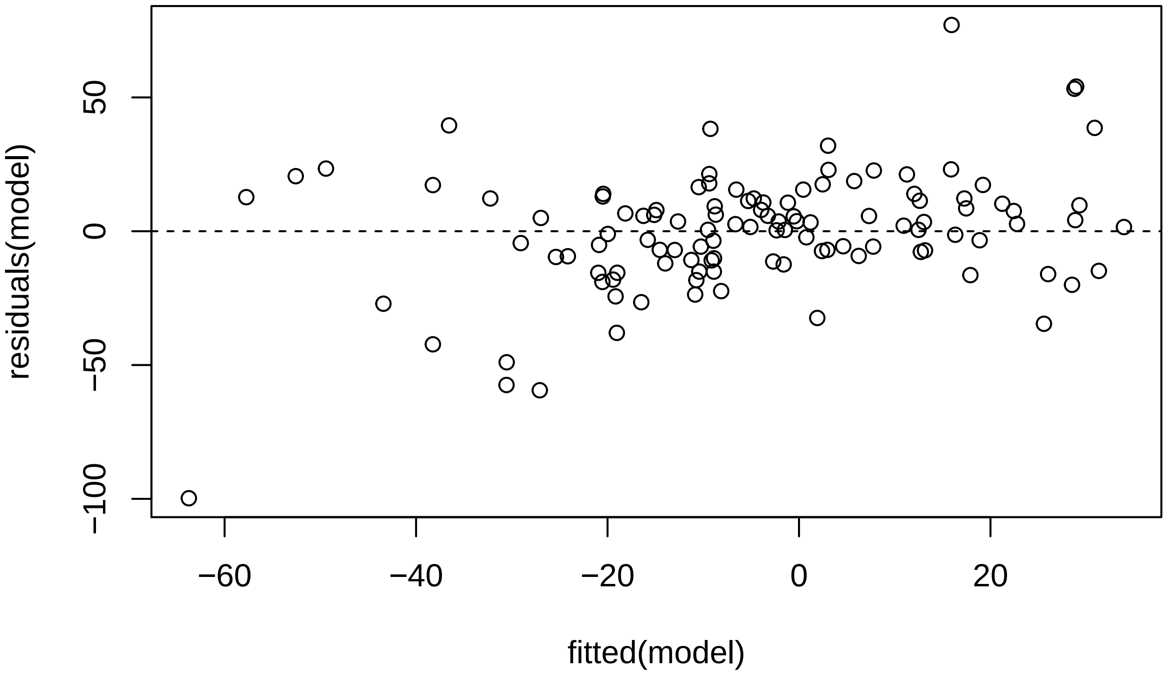

And the assumption of normality of residuals with histogram and qqplot:

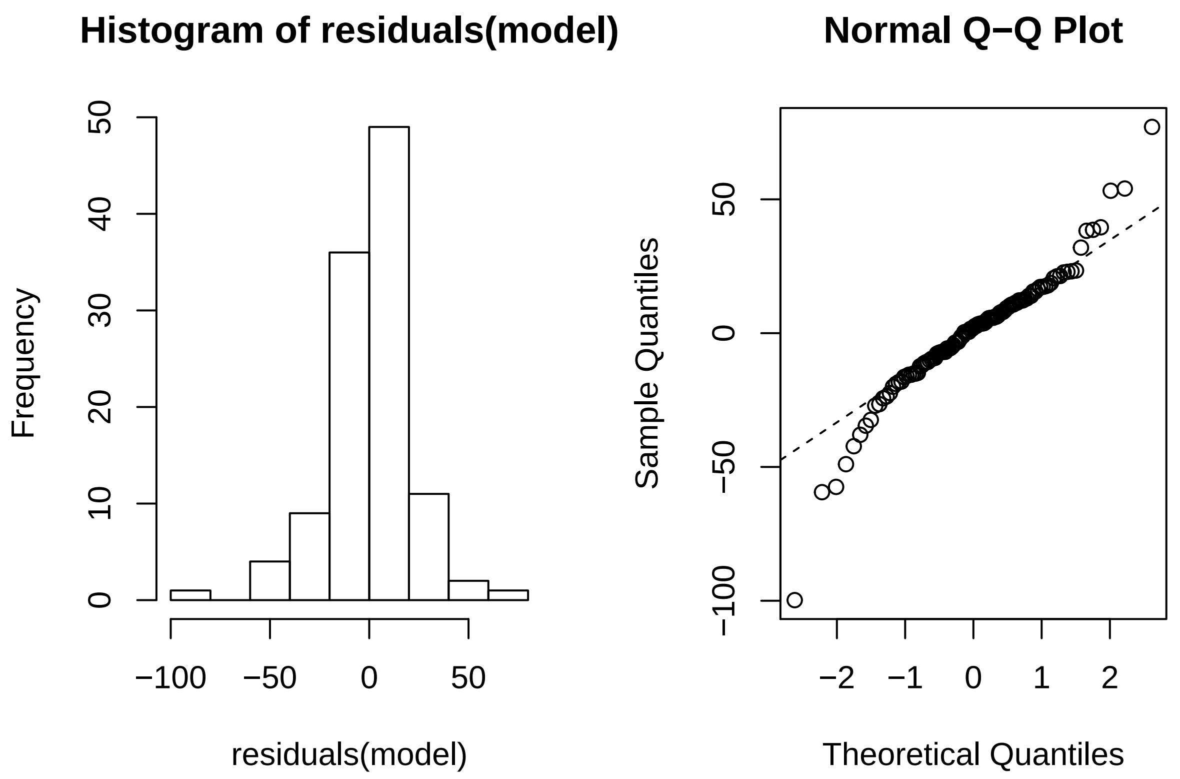

Using *Influence.ME*, outlying subjects are checked:

```
outlier <- influence(model, group = "Subject")
cks.d <- cooks.distance(outlier)
dfbetas <- dfbetas(outlier)
```

Now the Cook's distance values will be plotted. Outlier Subjects are highlighted in red:

```
plot(outlier,
     which = "cook",
     sort = TRUE,
     cutoff = 4/length(unique(dat$Subject)),
     xlab = "Cooks Distance")
```

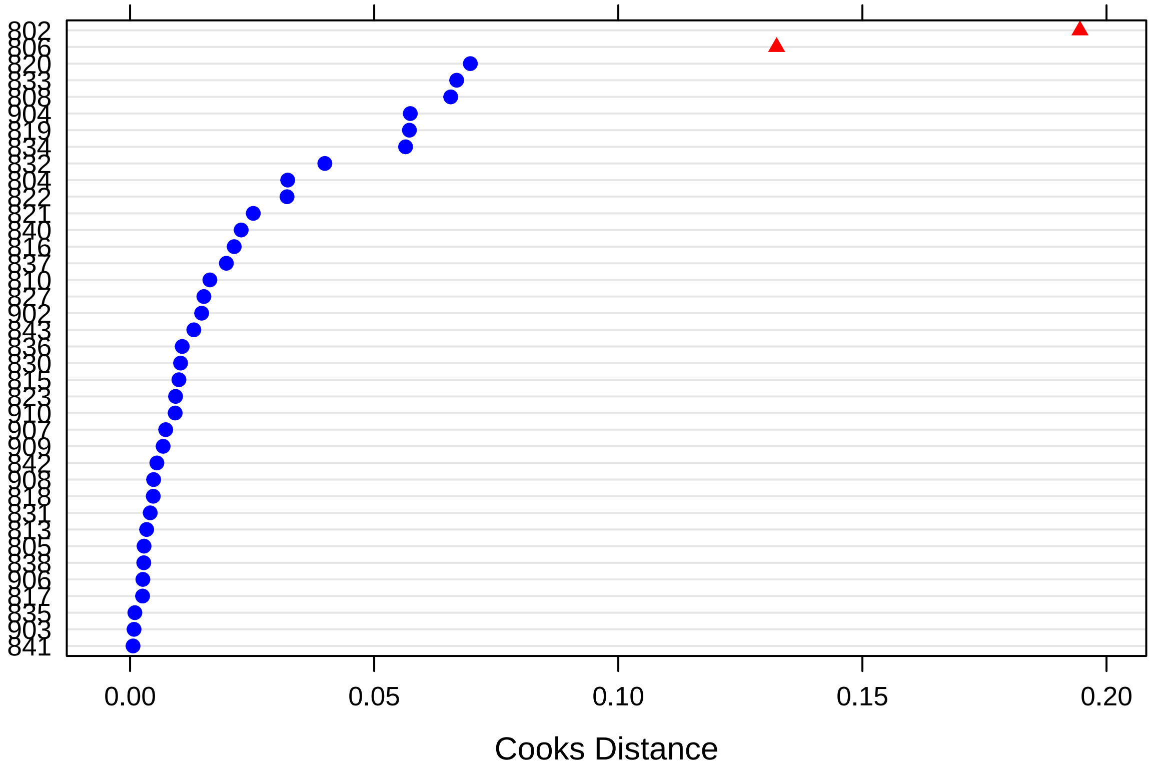

There are two outlying subjects (*802* & *806*).

Finally, these are the p-values for dfbeta values, that indicate whether exclusion of a specific subject suffices to change significance values:

```
sigtest(outlier, test = -1.96)
```

```
## $Intercept
##     Altered.Teststat Altered.Sig Changed.Sig
## 802        -1.211569       FALSE       FALSE
## 804        -1.440193       FALSE       FALSE
## 805        -1.656330       FALSE       FALSE
## 806        -2.320480        TRUE        TRUE
## 808        -1.856249       FALSE       FALSE
## 810        -1.948918       FALSE       FALSE
## 813        -1.707657       FALSE       FALSE
## 815        -1.873189       FALSE       FALSE
## 816        -1.706955       FALSE       FALSE
## 817        -1.626695       FALSE       FALSE
## 818        -1.708834       FALSE       FALSE
## 819        -1.755299       FALSE       FALSE
## 820        -1.866935       FALSE       FALSE
## 821        -1.577922       FALSE       FALSE
## 822        -1.947703       FALSE       FALSE
## 823        -1.829340       FALSE       FALSE
## 827        -1.902994       FALSE       FALSE
## 830        -1.633505       FALSE       FALSE
```

```
## 831        -1.702499       FALSE       FALSE
## 832        -1.938372       FALSE       FALSE
## 833        -2.013401        TRUE        TRUE
## 834        -1.752602       FALSE       FALSE
## 835        -1.716457       FALSE       FALSE
## 836        -1.718207       FALSE       FALSE
## 837        -1.680390       FALSE       FALSE
## 838        -1.709328       FALSE       FALSE
## 840        -1.984310        TRUE        TRUE
## 841        -1.760491       FALSE       FALSE
## 842        -1.844907       FALSE       FALSE
## 843        -1.619500       FALSE       FALSE
## 902        -1.511894       FALSE       FALSE
## 903        -1.753448       FALSE       FALSE
## 904        -1.339291       FALSE       FALSE
## 906        -1.761202       FALSE       FALSE
## 907        -1.613582       FALSE       FALSE
## 908        -1.771286       FALSE       FALSE
## 909        -1.794320       FALSE       FALSE
## 910        -1.679564       FALSE       FALSE
##
## $Sessionfac2
##     Altered.Teststat Altered.Sig Changed.Sig
## 802        0.8895049       FALSE       FALSE
## 804        0.9814339       FALSE       FALSE
## 805        1.1871260       FALSE       FALSE
## 806        1.9502347       FALSE       FALSE
## 808        1.3704189       FALSE       FALSE
## 810        1.3083090       FALSE       FALSE
## 813        1.1528613       FALSE       FALSE
## 815        1.4028655       FALSE       FALSE
## 816        1.4344812       FALSE       FALSE
## 817        1.2040236       FALSE       FALSE
## 818        1.3339411       FALSE       FALSE
## 819        1.3213764       FALSE       FALSE
## 820        1.7240806       FALSE       FALSE
## 821        1.3105172       FALSE       FALSE
## 822        1.4167545       FALSE       FALSE
## 823        1.1913408       FALSE       FALSE
## 827        1.3227498       FALSE       FALSE
## 830        1.2122296       FALSE       FALSE
## 831        1.1386845       FALSE       FALSE
## 832        1.1180792       FALSE       FALSE
## 833        1.0671606       FALSE       FALSE
## 834        1.4881592       FALSE       FALSE
## 835        1.2159587       FALSE       FALSE
## 836        1.0878828       FALSE       FALSE
## 837        1.1636110       FALSE       FALSE
## 838        1.2961454       FALSE       FALSE
## 840        1.2862640       FALSE       FALSE
## 841        1.2531810       FALSE       FALSE
## 842        1.2920867       FALSE       FALSE
## 843        1.0902405       FALSE       FALSE
## 902        1.1302949       FALSE       FALSE
```

```
## 903        1.2959524       FALSE       FALSE
## 904        1.0479555       FALSE       FALSE
## 906        1.3199855       FALSE       FALSE
## 907        1.2545575       FALSE       FALSE
## 908        1.3466437       FALSE       FALSE
## 909        1.3248801       FALSE       FALSE
## 910        1.1357402       FALSE       FALSE
##
## $Sessionfac3
##     Altered.Teststat Altered.Sig Changed.Sig
## 802         1.337534       FALSE       FALSE
## 804         1.524469       FALSE       FALSE
## 805         1.671728       FALSE       FALSE
## 806         2.305335       FALSE       FALSE
## 808         1.407810       FALSE       FALSE
## 810         1.703226       FALSE       FALSE
## 813         1.616661       FALSE       FALSE
## 815         1.827996       FALSE       FALSE
## 816         1.825054       FALSE       FALSE
## 817         1.600798       FALSE       FALSE
## 818         1.714553       FALSE       FALSE
## 819         2.060346       FALSE       FALSE
## 820         1.968922       FALSE       FALSE
## 821         1.732555       FALSE       FALSE
## 822         2.008487       FALSE       FALSE
## 823         1.689032       FALSE       FALSE
## 827         1.879743       FALSE       FALSE
## 830         1.531992       FALSE       FALSE
## 831         1.631131       FALSE       FALSE
## 832         1.824207       FALSE       FALSE
## 833         1.720992       FALSE       FALSE
## 834         1.990080       FALSE       FALSE
## 835         1.699471       FALSE       FALSE
## 836         1.637644       FALSE       FALSE
## 837         1.485775       FALSE       FALSE
## 838         1.641179       FALSE       FALSE
## 840         1.737898       FALSE       FALSE
## 841         1.690955       FALSE       FALSE
## 842         1.766313       FALSE       FALSE
## 843         1.545716       FALSE       FALSE
## 902         1.582246       FALSE       FALSE
## 903         1.695856       FALSE       FALSE
## 904         1.385018       FALSE       FALSE
## 906         1.674617       FALSE       FALSE
## 907         1.562483       FALSE       FALSE
## 908         1.673753       FALSE       FALSE
## 909         1.640701       FALSE       FALSE
## 910         1.555537       FALSE       FALSE
##
## $Groupcen
##     Altered.Teststat Altered.Sig Changed.Sig
## 802       -0.7243354       FALSE       FALSE
## 804       -1.5931063       FALSE       FALSE
## 805       -1.2171611       FALSE       FALSE
```

```
## 806       -0.7948605       FALSE       FALSE
## 808       -1.2102883       FALSE       FALSE
## 810       -1.0814332       FALSE       FALSE
## 813       -1.2870970       FALSE       FALSE
## 815       -1.1276182       FALSE       FALSE
## 816       -1.3200196       FALSE       FALSE
## 817       -1.3679027       FALSE       FALSE
## 818       -1.2689998       FALSE       FALSE
## 819       -1.3046516       FALSE       FALSE
## 820       -1.4151405       FALSE       FALSE
## 821       -1.4735383       FALSE       FALSE
## 822       -1.5039585       FALSE       FALSE
## 823       -1.3871723       FALSE       FALSE
## 827       -1.1065000       FALSE       FALSE
## 830       -1.1934690       FALSE       FALSE
## 831       -1.2926666       FALSE       FALSE
## 832       -1.1183674       FALSE       FALSE
## 833       -1.5510445       FALSE       FALSE
## 834       -1.2968648       FALSE       FALSE
## 835       -1.2748403       FALSE       FALSE
## 836       -1.2855653       FALSE       FALSE
## 837       -1.2383171       FALSE       FALSE
## 838       -1.2704825       FALSE       FALSE
## 840       -1.5357608       FALSE       FALSE
## 841       -1.3218667       FALSE       FALSE
## 842       -1.4047602       FALSE       FALSE
## 843       -1.1791334       FALSE       FALSE
## 902       -1.5173479       FALSE       FALSE
## 903       -1.3149583       FALSE       FALSE
## 904       -1.7446084       FALSE       FALSE
## 906       -1.3224003       FALSE       FALSE
## 907       -1.3867081       FALSE       FALSE
## 908       -1.3321348       FALSE       FALSE
## 909       -1.3543098       FALSE       FALSE
## 910       -1.2394252       FALSE       FALSE
##
## $`Sessionfac2:Groupcen`
##     Altered.Teststat Altered.Sig Changed.Sig
## 802      -0.82293885       FALSE       FALSE
## 804      -0.01261483       FALSE       FALSE
## 805      -0.35557340       FALSE       FALSE
## 806      -0.91585607       FALSE       FALSE
## 808      -0.38494913       FALSE       FALSE
## 810      -0.35500606       FALSE       FALSE
## 813      -0.19876919       FALSE       FALSE
## 815      -0.44432394       FALSE       FALSE
## 816      -0.47482827       FALSE       FALSE
## 817      -0.25073481       FALSE       FALSE
## 818      -0.20979244       FALSE       FALSE
## 819      -0.28236628       FALSE       FALSE
## 820       0.10118367       FALSE       FALSE
## 821      -0.35741378       FALSE       FALSE
## 822      -0.16586100       FALSE       FALSE
## 823      -0.35163958       FALSE       FALSE
```

```
## 827      -0.36256846       FALSE       FALSE
## 830      -0.34215394       FALSE       FALSE
## 831      -0.18397959       FALSE       FALSE
## 832      -0.15109179       FALSE       FALSE
## 833      -0.50194698       FALSE       FALSE
## 834      -0.09188844       FALSE       FALSE
## 835      -0.26304412       FALSE       FALSE
## 836      -0.12974749       FALSE       FALSE
## 837      -0.40122873       FALSE       FALSE
## 838      -0.24783749       FALSE       FALSE
## 840      -0.25546664       FALSE       FALSE
## 841      -0.28717264       FALSE       FALSE
## 842      -0.25182746       FALSE       FALSE
## 843      -0.46826410       FALSE       FALSE
## 902      -0.16791521       FALSE       FALSE
## 903      -0.24542149       FALSE       FALSE
## 904      -0.07358526       FALSE       FALSE
## 906      -0.22380214       FALSE       FALSE
## 907      -0.29870417       FALSE       FALSE
## 908      -0.20019270       FALSE       FALSE
## 909      -0.22228740       FALSE       FALSE
## 910      -0.41595316       FALSE       FALSE
##
## $`Sessionfac3:Groupcen`
##     Altered.Teststat Altered.Sig Changed.Sig
## 802      -0.43317140       FALSE       FALSE
## 804       0.25695446       FALSE       FALSE
## 805       0.07636198       FALSE       FALSE
## 806      -0.40373555       FALSE       FALSE
## 808       0.40417100       FALSE       FALSE
## 810       0.04981487       FALSE       FALSE
## 813       0.13770794       FALSE       FALSE
## 815      -0.06547079       FALSE       FALSE
## 816      -0.06041262       FALSE       FALSE
## 817       0.15210144       FALSE       FALSE
## 818       0.11782688       FALSE       FALSE
## 819       0.40637033       FALSE       FALSE
## 820       0.29834236       FALSE       FALSE
## 821       0.02019151       FALSE       FALSE
## 822       0.37702547       FALSE       FALSE
## 823       0.09108031       FALSE       FALSE
## 827      -0.11419146       FALSE       FALSE
## 830      -0.07355488       FALSE       FALSE
## 831       0.12435988       FALSE       FALSE
## 832      -0.04605716       FALSE       FALSE
## 833       0.08550649       FALSE       FALSE
## 834       0.35138086       FALSE       FALSE
## 835       0.05273273       FALSE       FALSE
## 836       0.12414765       FALSE       FALSE
## 837      -0.12986549       FALSE       FALSE
## 838       0.04502091       FALSE       FALSE
## 840       0.13578118       FALSE       FALSE
## 841       0.09791806       FALSE       FALSE
## 842       0.16919662       FALSE       FALSE
```

```
## 843      -0.06344825       FALSE       FALSE
## 902       0.20884976       FALSE       FALSE
## 903       0.10209599       FALSE       FALSE
## 904       0.40672052       FALSE       FALSE
## 906       0.07865128       FALSE       FALSE
## 907       0.19512377       FALSE       FALSE
## 908       0.07503635       FALSE       FALSE
## 909       0.04104901       FALSE       FALSE
## 910      -0.04789677       FALSE       FALSE
```

There is no indication that results are changed by.

#### Avoidance (AAT)

The base model is now tested against the same models without the effects of time:

```
model <- lmer(AAT ~ Sessionfac*Groupcen + (1 + Sessioncen | Subject),
              data = dat, REML = FALSE)
```

```
## singular fit
```

```
model_null <- lmer(AAT ~ Sessionfac + Groupcen + (1 + Sessioncen | Subject),
              data = dat, REML = FALSE)
```

These are the results, which **do not** support rejection of null hypothesis:

```
anova(model, model_null)
```

```
## Data: dat
## Models:
## model_null: AAT ~ Sessionfac + Groupcen + (1 + Sessioncen | Subject)
## model: AAT ~ Sessionfac * Groupcen + (1 + Sessioncen | Subject)
##            Df    AIC    BIC  logLik deviance  Chisq Chi Df Pr(>Chisq)
## model_null  8 1238.1 1259.8 -611.05   1222.1
## model      10 1241.7 1268.8 -610.85   1221.7 0.3928      2     0.8217
```

This is the final model (re-estimated with REML=TRUE):

```
model <- lmer(AAT ~ Sessionfac*Groupcen + (1 + Sessioncen | Subject),
              data = dat, REML = TRUE)

summary(model)
```

```
## Linear mixed model fit by REML ['lmerMod']
## Formula: AAT ~ Sessionfac * Groupcen + (1 + Sessioncen | Subject)
##    Data: dat
##
## REML criterion at convergence: 1183.2
##
## Scaled residuals:
##      Min       1Q   Median       3Q      Max
## -2.37637 -0.49432  0.07119  0.56931  1.82550
##
## Random effects:
##  Groups   Name        Variance Std.Dev. Corr
##  Subject  (Intercept) 2686     51.83
##           Sessioncen  1815     42.60    -1.00
```

```
##  Residual              1939     44.04
## Number of obs: 111, groups:  Subject, 38
##
## Fixed effects:
##                      Estimate Std. Error t value
## (Intercept)          -12.3533    17.0950  -0.723
## Sessionfac2           -0.1865    12.5964  -0.015
## Sessionfac3           10.5515    17.3135   0.609
## Groupcen              -5.0886    17.0950  -0.298
## Sessionfac2:Groupcen   5.2846    12.5964   0.420
## Sessionfac3:Groupcen  10.5809    17.3135   0.611
##
## Correlation of Fixed Effects:
##             (Intr) Sssnf2 Sssnf3 Gropcn Sss2:G
## Sessionfac2 -0.753
## Sessionfac3 -0.909  0.695
## Groupcen    -0.093  0.062  0.083
## Sssnfc2:Grp  0.062 -0.083 -0.056 -0.753
## Sssnfc3:Grp  0.083 -0.056 -0.093 -0.909  0.695
```

**Assumptions**

Here are the residuals plotted against fitted values to investigate underlying heteroskedasticity and linearity assumptions:

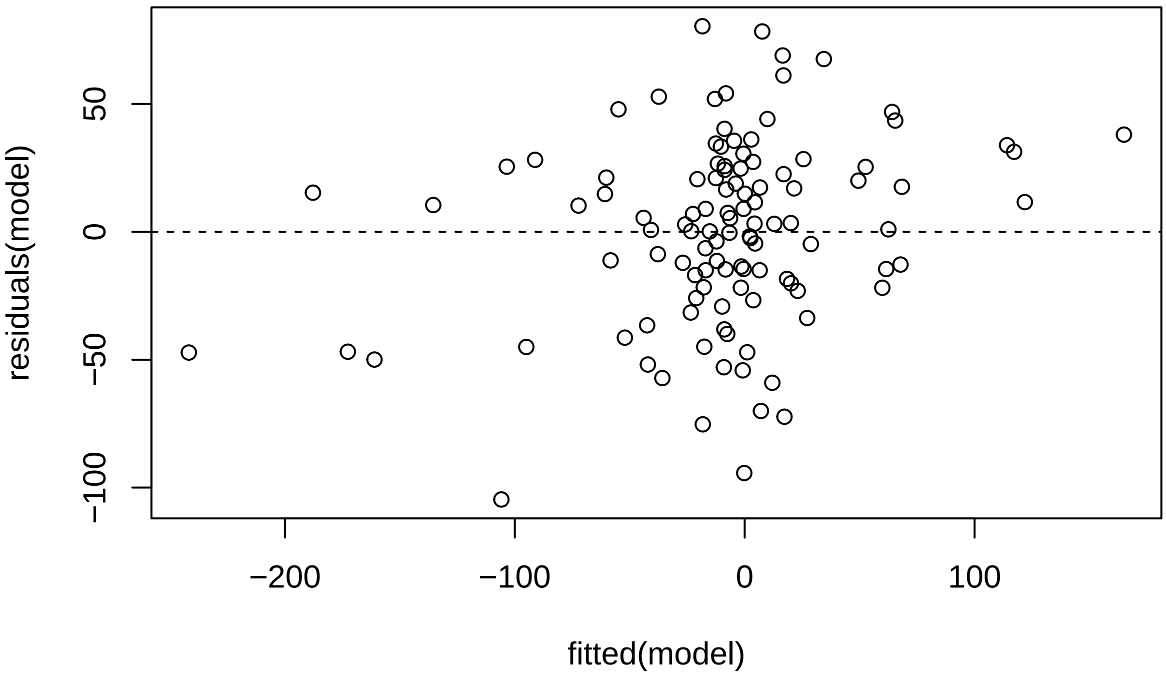

And the assumption of normality of residuals with histogram and qqplot:

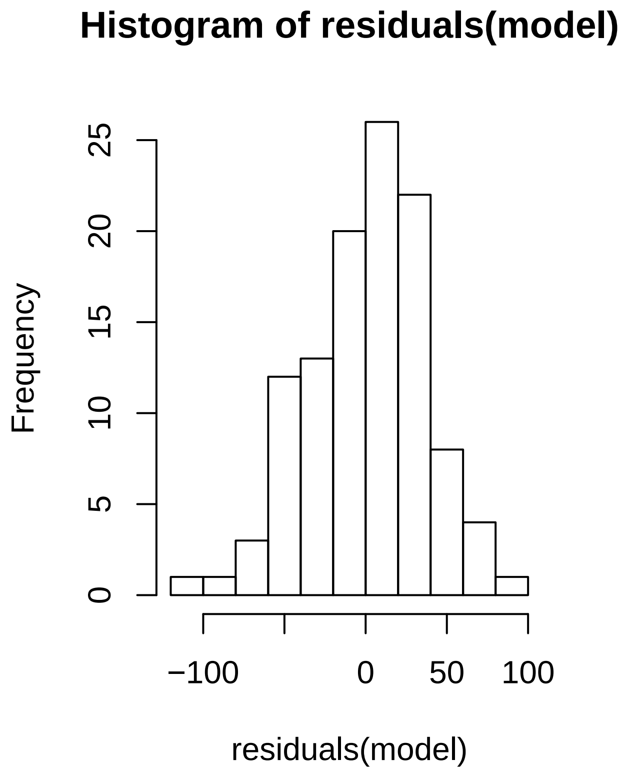

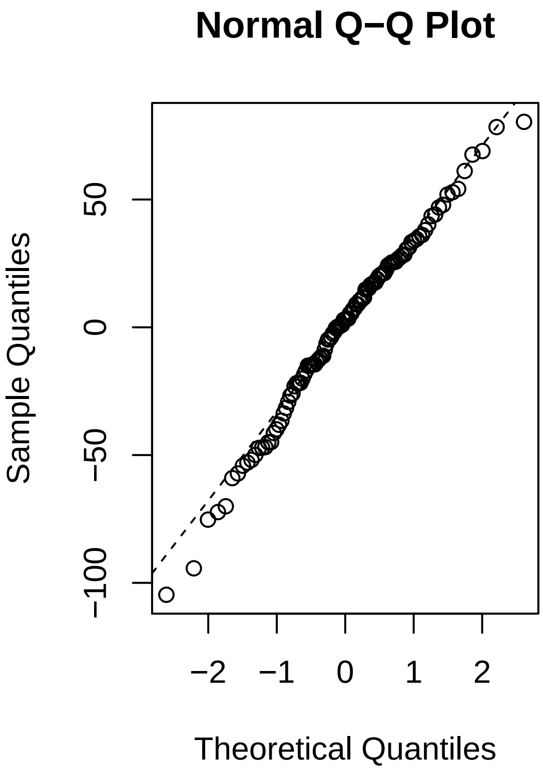

Using *Influence.ME*, outlying subjects are checked:

```
outlier <- influence(model, group = "Subject")
```

```
## singular fit
## singular fit
## singular fit
## singular fit
## singular fit
## singular fit
## singular fit
## singular fit
## singular fit
## singular fit
## singular fit
## singular fit
## singular fit
## singular fit
## singular fit
## singular fit
## singular fit
## singular fit
## singular fit
## singular fit
## singular fit
## singular fit
## singular fit
```

```
cks.d <- cooks.distance(outlier)
dfbetas <- dfbetas(outlier)
```

Now the Cook's distance values will be plotted. Outlier Subjects are highlighted in red:

```
plot(outlier,
     which = "cook",
     sort = TRUE,
     cutoff = 4/length(unique(dat$Subject)),
     xlab = "Cooks Distance")
```

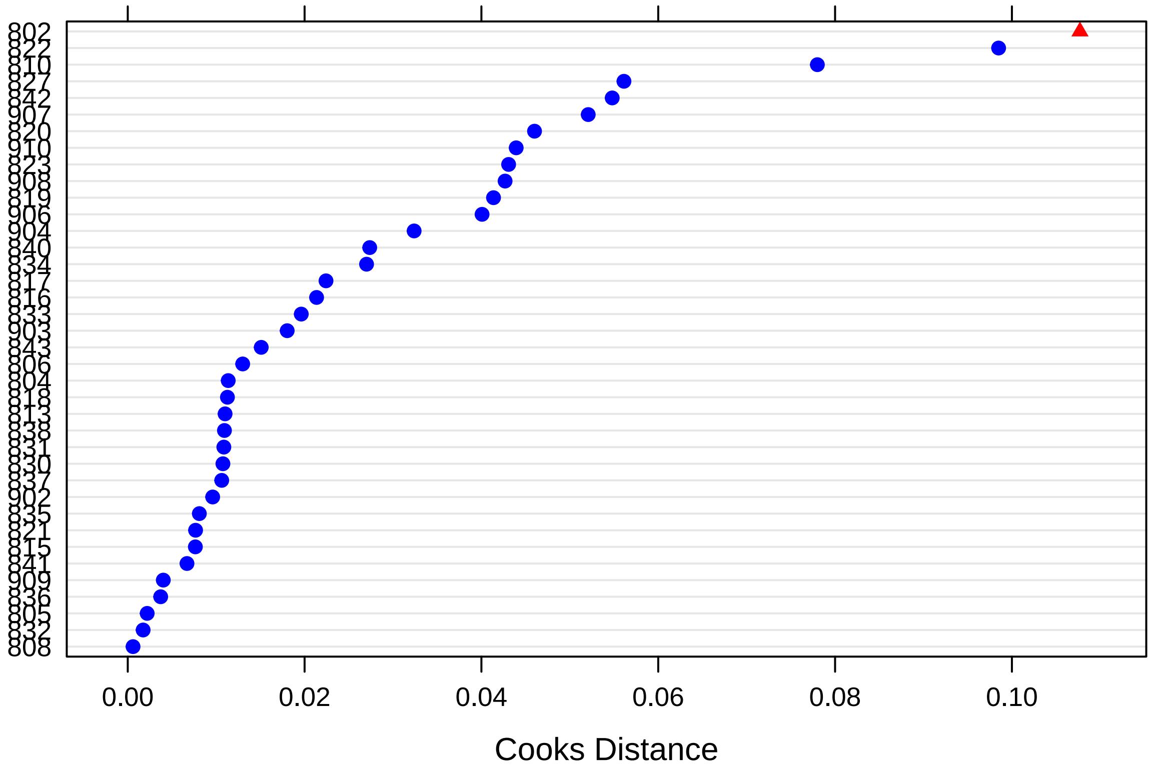

There is one outlying subject (*822*).

Finally, these are the p-values for dfbeta values, that indicate whether exclusion of a specific subject suffices to change significance values:

```
sigtest(outlier, test = -1.96)
```

```
## $Intercept
##     Altered.Teststat Altered.Sig Changed.Sig
## 802       -0.3216146       FALSE       FALSE
## 804       -0.8335326       FALSE       FALSE
## 805       -0.7533283       FALSE       FALSE
## 806       -0.6874671       FALSE       FALSE
## 808       -0.7134626       FALSE       FALSE
## 810       -0.9288099       FALSE       FALSE
## 813       -0.8409578       FALSE       FALSE
## 815       -0.7556753       FALSE       FALSE
```

```
## 816        -0.7683070       FALSE        FALSE
## 817        -0.6360365       FALSE        FALSE
## 818        -0.6450910       FALSE        FALSE
## 819        -0.7260977       FALSE        FALSE
## 820        -0.9714089       FALSE        FALSE
## 821        -0.6463459       FALSE        FALSE
## 822        -0.5171777       FALSE        FALSE
## 823        -0.4384604       FALSE        FALSE
## 827        -0.3382670       FALSE        FALSE
## 830        -0.6691560       FALSE        FALSE
## 831        -0.8589639       FALSE        FALSE
## 832        -0.6870873       FALSE        FALSE
## 833        -0.6809732       FALSE        FALSE
## 834        -0.7966942       FALSE        FALSE
## 835        -0.6737341       FALSE        FALSE
## 836        -0.5927694       FALSE        FALSE
## 837        -0.7574403       FALSE        FALSE
## 838        -0.6759127       FALSE        FALSE
## 840        -0.5294462       FALSE        FALSE
## 841        -0.6978260       FALSE        FALSE
## 842        -0.9669170       FALSE        FALSE
## 843        -0.6471474       FALSE        FALSE
## 902        -0.8738949       FALSE        FALSE
## 903        -0.6686332       FALSE        FALSE
## 904        -0.5533259       FALSE        FALSE
## 906        -0.9728387       FALSE        FALSE
## 907        -0.6063341       FALSE        FALSE
## 908        -1.0900161       FALSE        FALSE
## 909        -0.7199195       FALSE        FALSE
## 910        -0.8209369       FALSE        FALSE
##
## $Sessionfac2
##     Altered.Teststat Altered.Sig Changed.Sig
## 802    -3.818330e-01       FALSE       FALSE
## 804    -1.123263e-02       FALSE       FALSE
## 805     1.171709e-02       FALSE       FALSE
## 806     5.547846e-02       FALSE       FALSE
## 808    -2.908556e-02       FALSE       FALSE
## 810     2.831668e-01       FALSE       FALSE
## 813     1.428809e-01       FALSE       FALSE
## 815     2.795143e-02       FALSE       FALSE
## 816     2.087163e-01       FALSE       FALSE
## 817     1.002111e-01       FALSE       FALSE
## 818     3.515471e-02       FALSE       FALSE
## 819    -1.227331e-01       FALSE       FALSE
## 820     2.168106e-01       FALSE       FALSE
## 821    -1.234664e-01       FALSE       FALSE
## 822     9.401812e-02       FALSE       FALSE
## 823    -3.193499e-01       FALSE       FALSE
## 827    -3.910804e-01       FALSE       FALSE
## 830    -1.342184e-01       FALSE       FALSE
## 831     1.661134e-01       FALSE       FALSE
## 832    -5.600085e-03       FALSE       FALSE
## 833     1.170309e-01       FALSE       FALSE
```

```
## 834     1.053666e-01       FALSE       FALSE
## 835    -3.043945e-02       FALSE       FALSE
## 836    -9.964685e-02       FALSE       FALSE
## 837     1.328799e-01       FALSE       FALSE
## 838    -6.376977e-05       FALSE       FALSE
## 840    -2.838913e-01       FALSE       FALSE
## 841     1.710984e-02       FALSE       FALSE
## 842     4.379675e-02       FALSE       FALSE
## 843    -3.769733e-02       FALSE       FALSE
## 902     1.206757e-01       FALSE       FALSE
## 903    -1.849184e-01       FALSE       FALSE
## 904    -3.007890e-01       FALSE       FALSE
## 906     1.966915e-01       FALSE       FALSE
## 907    -3.447110e-01       FALSE       FALSE
## 908     2.323000e-01       FALSE       FALSE
## 909     3.045386e-02       FALSE       FALSE
## 910    -1.065478e-01       FALSE       FALSE
##
## $Sessionfac3
##     Altered.Teststat Altered.Sig Changed.Sig
## 802        0.3382559       FALSE       FALSE
## 804        0.7343559       FALSE       FALSE
## 805        0.6127732       FALSE       FALSE
## 806        0.5111573       FALSE       FALSE
## 808        0.5923999       FALSE       FALSE
## 810        0.9971220       FALSE       FALSE
## 813        0.6741233       FALSE       FALSE
## 815        0.5772302       FALSE       FALSE
## 816        0.6591909       FALSE       FALSE
## 817        0.4991952       FALSE       FALSE
## 818        0.5713984       FALSE       FALSE
## 819        0.4972555       FALSE       FALSE
## 820        0.7236952       FALSE       FALSE
## 821        0.4957508       FALSE       FALSE
## 822        0.3520372       FALSE       FALSE
## 823        0.3019435       FALSE       FALSE
## 827        0.3114809       FALSE       FALSE
## 830        0.5212715       FALSE       FALSE
## 831        0.7623673       FALSE       FALSE
## 832        0.5500493       FALSE       FALSE
## 833        0.5648633       FALSE       FALSE
## 834        0.7875777       FALSE       FALSE
## 835        0.6339766       FALSE       FALSE
## 836        0.4986418       FALSE       FALSE
## 837        0.6537043       FALSE       FALSE
## 838        0.6350381       FALSE       FALSE
## 840        0.4070091       FALSE       FALSE
## 841        0.5388967       FALSE       FALSE
## 842        0.9510641       FALSE       FALSE
## 843        0.6185823       FALSE       FALSE
## 902        0.7189417       FALSE       FALSE
## 903        0.5419145       FALSE       FALSE
## 904        0.4094118       FALSE       FALSE
## 906        0.7348380       FALSE       FALSE
```

```
## 907        0.5293827       FALSE       FALSE
## 908        0.9020712       FALSE       FALSE
## 909        0.6476172       FALSE       FALSE
## 910        0.7867115       FALSE       FALSE
##
## $Groupcen
##     Altered.Teststat Altered.Sig Changed.Sig
## 802      0.133603447       FALSE       FALSE
## 804     -0.164397877       FALSE       FALSE
## 805     -0.338611433       FALSE       FALSE
## 806     -0.304231058       FALSE       FALSE
## 808     -0.275685663       FALSE       FALSE
## 810     -0.089510247       FALSE       FALSE
## 813     -0.153001576       FALSE       FALSE
## 815     -0.234160166       FALSE       FALSE
## 816     -0.223749822       FALSE       FALSE
## 817     -0.359727807       FALSE       FALSE
## 818     -0.229352110       FALSE       FALSE
## 819     -0.310891166       FALSE       FALSE
## 820     -0.545362100       FALSE       FALSE
## 821     -0.344779908       FALSE       FALSE
## 822     -0.077828855       FALSE       FALSE
## 823     -0.004014801       FALSE       FALSE
## 827     -0.697167432       FALSE       FALSE
## 830     -0.254517186       FALSE       FALSE
## 831     -0.138953589       FALSE       FALSE
## 832     -0.302433622       FALSE       FALSE
## 833     -0.265791840       FALSE       FALSE
## 834     -0.380624826       FALSE       FALSE
## 835     -0.317206002       FALSE       FALSE
## 836     -0.400440039       FALSE       FALSE
## 837     -0.343367754       FALSE       FALSE
## 838     -0.261556181       FALSE       FALSE
## 840     -0.108599536       FALSE       FALSE
## 841     -0.283524295       FALSE       FALSE
## 842     -0.533645417       FALSE       FALSE
## 843     -0.227192320       FALSE       FALSE
## 902     -0.123636542       FALSE       FALSE
## 903     -0.253799706       FALSE       FALSE
## 904     -0.446307800       FALSE       FALSE
## 906     -0.546029111       FALSE       FALSE
## 907     -0.389719635       FALSE       FALSE
## 908     -0.648564567       FALSE       FALSE
## 909     -0.305828776       FALSE       FALSE
## 910     -0.399283010       FALSE       FALSE
##
## $`Sessionfac2:Groupcen`
##     Altered.Teststat Altered.Sig Changed.Sig
## 802       0.04781476       FALSE       FALSE
## 804       0.40412707       FALSE       FALSE
## 805       0.43464195       FALSE       FALSE
## 806       0.34307661       FALSE       FALSE
## 808       0.42102640       FALSE       FALSE
## 810       0.15991302       FALSE       FALSE
```

```
## 813        0.24868749       FALSE       FALSE
## 815        0.36430605       FALSE       FALSE
## 816        0.19256953       FALSE       FALSE
## 817        0.29792544       FALSE       FALSE
## 818        0.45906559       FALSE       FALSE
## 819        0.30630110       FALSE       FALSE
## 820        0.64440169       FALSE       FALSE
## 821        0.52013743       FALSE       FALSE
## 822        0.54365130       FALSE       FALSE
## 823        0.12213725       FALSE       FALSE
## 827        0.77267625       FALSE       FALSE
## 830        0.29122204       FALSE       FALSE
## 831        0.23132319       FALSE       FALSE
## 832        0.39854361       FALSE       FALSE
## 833        0.54430760       FALSE       FALSE
## 834        0.53812548       FALSE       FALSE
## 835        0.43067751       FALSE       FALSE
## 836        0.49058991       FALSE       FALSE
## 837        0.55882495       FALSE       FALSE
## 838        0.42442000       FALSE       FALSE
## 840        0.14934316       FALSE       FALSE
## 841        0.44209938       FALSE       FALSE
## 842        0.48917679       FALSE       FALSE
## 843        0.39483395       FALSE       FALSE
## 902        0.26804138       FALSE       FALSE
## 903        0.24185858       FALSE       FALSE
## 904        0.71173996       FALSE       FALSE
## 906        0.62312967       FALSE       FALSE
## 907        0.75949763       FALSE       FALSE
## 908        0.66797992       FALSE       FALSE
## 909        0.45449374       FALSE       FALSE
## 910        0.33015368       FALSE       FALSE
##
## $`Sessionfac3:Groupcen`
##     Altered.Teststat Altered.Sig Changed.Sig
## 802        0.3399830       FALSE       FALSE
## 804        0.4589292       FALSE       FALSE
## 805        0.6144286       FALSE       FALSE
## 806        0.6780192       FALSE       FALSE
## 808        0.5906503       FALSE       FALSE
## 810        0.2531369       FALSE       FALSE
## 813        0.5125700       FALSE       FALSE
## 815        0.6052126       FALSE       FALSE
## 816        0.5309330       FALSE       FALSE
## 817        0.6929938       FALSE       FALSE
## 818        0.5730541       FALSE       FALSE
## 819        0.4989090       FALSE       FALSE
## 820        0.7253689       FALSE       FALSE
## 821        0.6923600       FALSE       FALSE
## 822        0.3538020       FALSE       FALSE
## 823        0.3037030       FALSE       FALSE
## 827        0.9125511       FALSE       FALSE
## 830        0.5229309       FALSE       FALSE
## 831        0.4349978       FALSE       FALSE
```

```
## 832        0.6340894       FALSE       FALSE
## 833        0.5665258       FALSE       FALSE
## 834        0.7892592       FALSE       FALSE
## 835        0.5516372       FALSE       FALSE
## 836        0.6883267       FALSE       FALSE
## 837        0.6553647       FALSE       FALSE
## 838        0.6366923       FALSE       FALSE
## 840        0.4087059       FALSE       FALSE
## 841        0.5405561       FALSE       FALSE
## 842        0.9528258       FALSE       FALSE
## 843        0.6202575       FALSE       FALSE
## 902        0.4705420       FALSE       FALSE
## 903        0.5435749       FALSE       FALSE
## 904        0.7970056       FALSE       FALSE
## 906        0.7365135       FALSE       FALSE
## 907        0.6707669       FALSE       FALSE
## 908        0.9038092       FALSE       FALSE
## 909        0.6492741       FALSE       FALSE
## 910        0.7884117       FALSE       FALSE
```

There is no indication that results are changed.

### Prediction of Spider Anxiety Improvements Bias Measures

To test the predictive value of bias change following treatment, linear regression will be applied. Specifically, linear regression of 1-month spider anxiety measures (SAS, FSQ, & BAT) on 1-day follow-up bias measures of AAT and EAST will be run. If these measures are significant, it will further be evaluated whether inclusion of spider anxiety covariates at 1-day follow-up will adjust effects of bias measures.

#### SAS

##### Unadjusted

This is the linear model:

```
model <- lm(SAS_4 ~ AAT_3 + EAST_3, data = dat[dat$WideData == 1,])
```

These are the results:

```
summary(model)
```

```
##
## Call:
## lm(formula = SAS_4 ~ AAT_3 + EAST_3, data = dat[dat$WideData ==
##     1, ])
##
## Residuals:
##     Min      1Q  Median      3Q     Max
## -8.7185 -2.2934 -0.1165  2.7371 11.8256
##
## Coefficients:
##             Estimate Std. Error t value Pr(>|t|)
## (Intercept) 10.80966    0.85896  12.585 3.82e-14 ***
```

```
## AAT_3       -0.02100    0.01338  -1.570    0.126
## EAST_3      -0.04197    0.02519  -1.666    0.105
## ---
## Signif. codes:  0 '***' 0.001 '**' 0.01 '*' 0.05 '.' 0.1 ' ' 1
##
## Residual standard error: 4.975 on 33 degrees of freedom
##   (2 observations deleted due to missingness)
## Multiple R-squared:  0.1224, Adjusted R-squared:  0.06925
## F-statistic: 2.302 on 2 and 33 DF,  p-value: 0.1159
```

There is no significant bias effect.

##### Assumptions

Here are the residuals plotted against fitted values to investigate underlying heteroskedasticity and linearity assumptions:

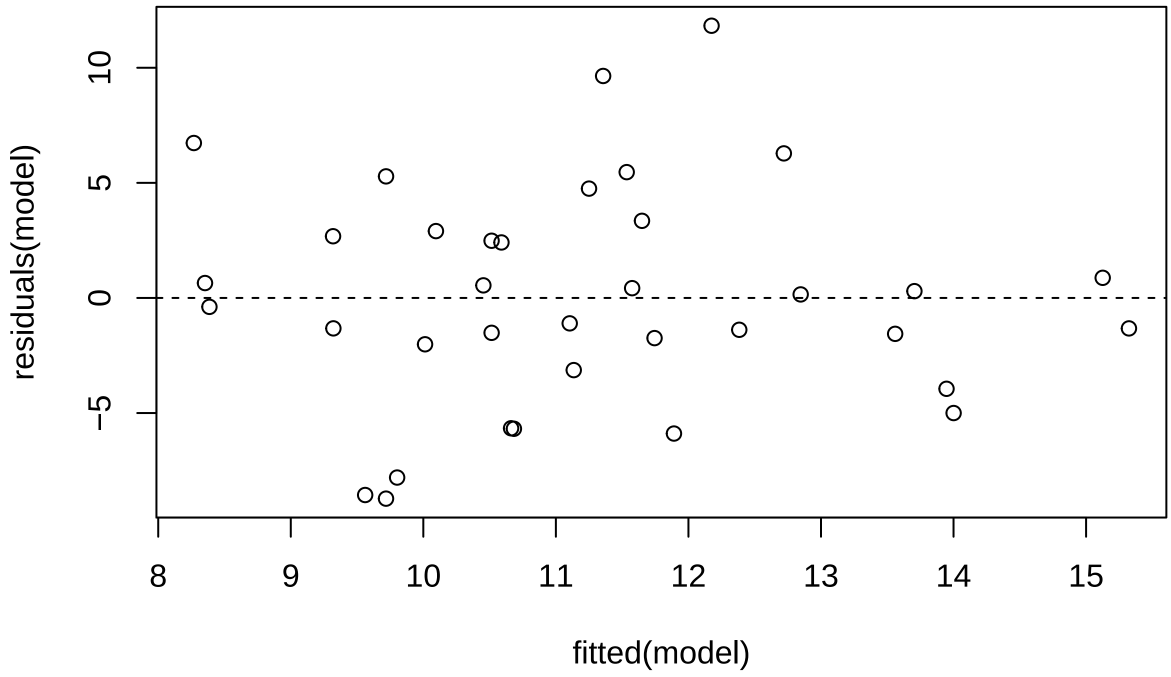

And the assumption of normality of residuals with histogram and qqplot:

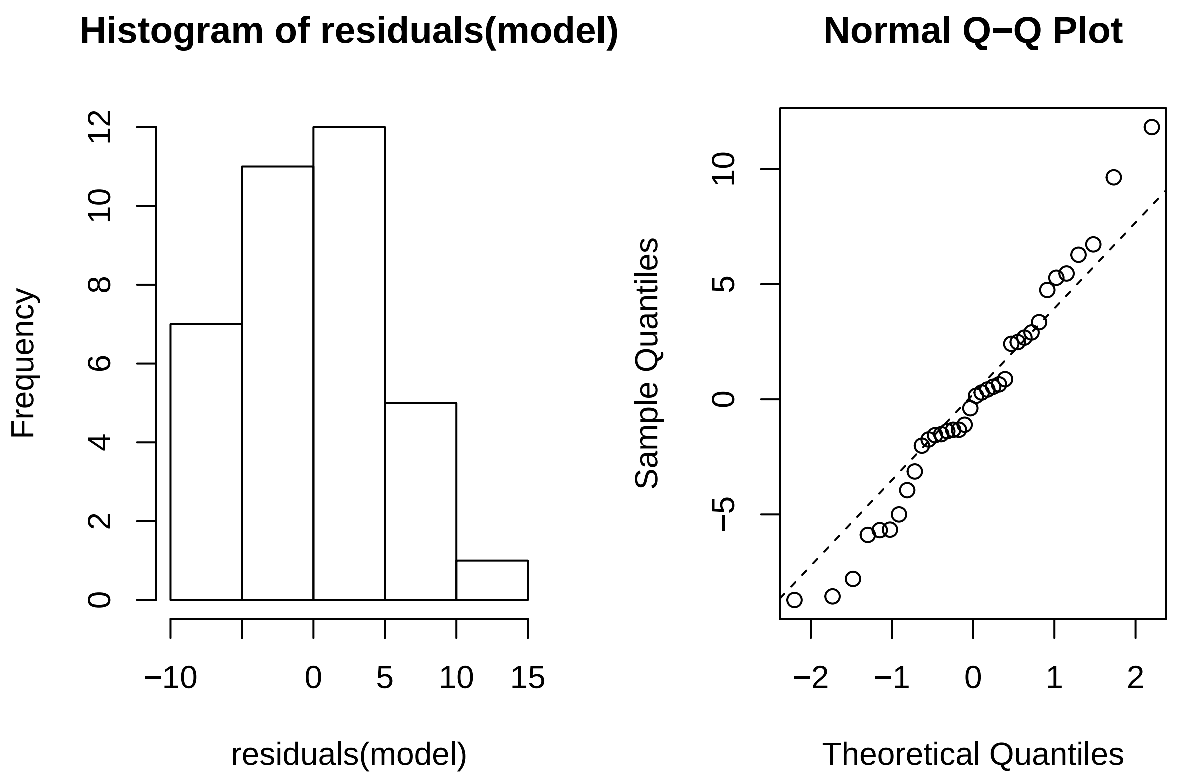

###### Adjusted

This is the linear model:

```
model <- lm(SAS_4 ~ AAT_3 + EAST_3 + SAS_3, data = dat[dat$WideData == 1,])
```

These are the results:

```
summary(model)
```

```
##
## Call:
## lm(formula = SAS_4 ~ AAT_3 + EAST_3 + SAS_3, data = dat[dat$WideData ==
##     1, ])
##
## Residuals:
##    Min     1Q Median     3Q    Max
## -7.653 -1.803  0.142  1.909  6.222
##
## Coefficients:
##              Estimate Std. Error t value Pr(>|t|)
## (Intercept) -0.813375   1.721835  -0.472    0.640
## AAT_3       -0.003458   0.008809  -0.393    0.697
## EAST_3      -0.015011   0.016364  -0.917    0.366
## SAS_3        0.914668   0.128586   7.113 4.52e-08 ***
## ---
```

```
## Signif. codes:  0 '***' 0.001 '**' 0.01 '*' 0.05 '.' 0.1 ' ' 1
##
## Residual standard error: 3.145 on 32 degrees of freedom
##   (2 observations deleted due to missingness)
## Multiple R-squared:   0.66,  Adjusted R-squared:  0.6281
## F-statistic: 20.71 on 3 and 32 DF,  p-value: 1.213e-07
```

There is no significant bias effect.

###### Assumptions

Here are the residuals plotted against fitted values to investigate underlying heteroskedasticity and linearity assumptions:

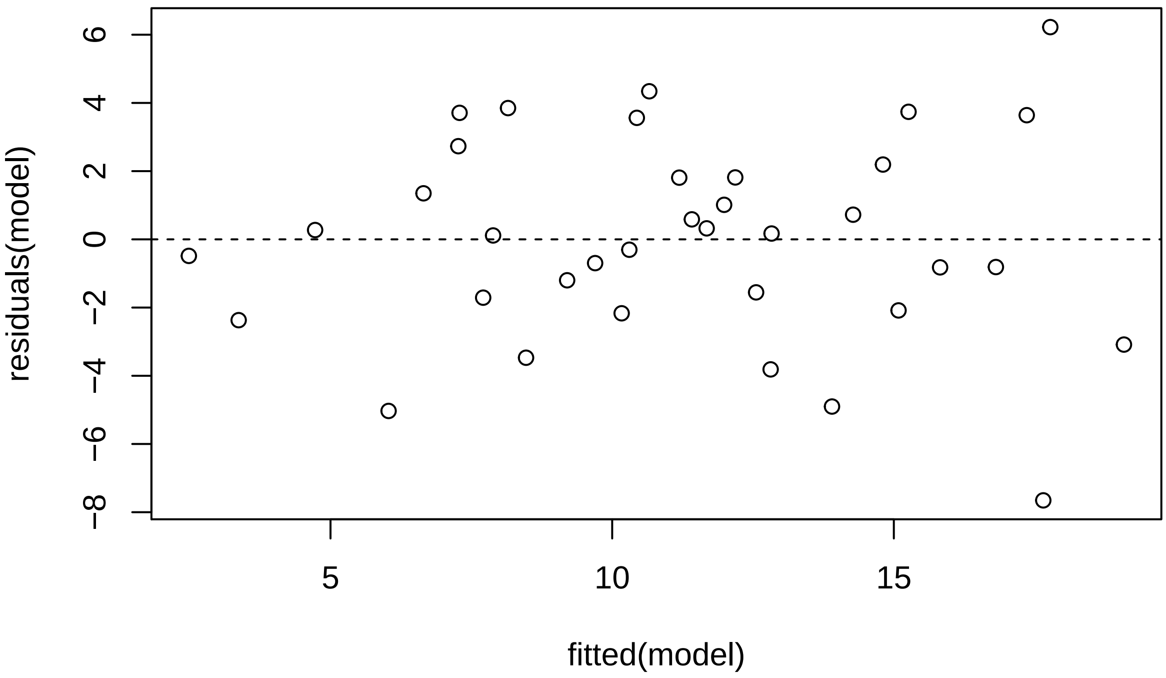

And the assumption of normality of residuals with histogram and qqplot:

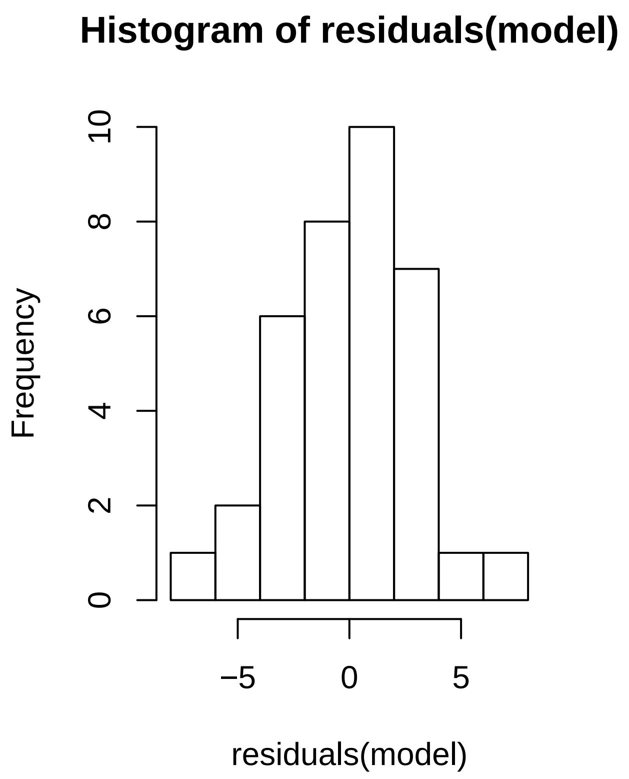

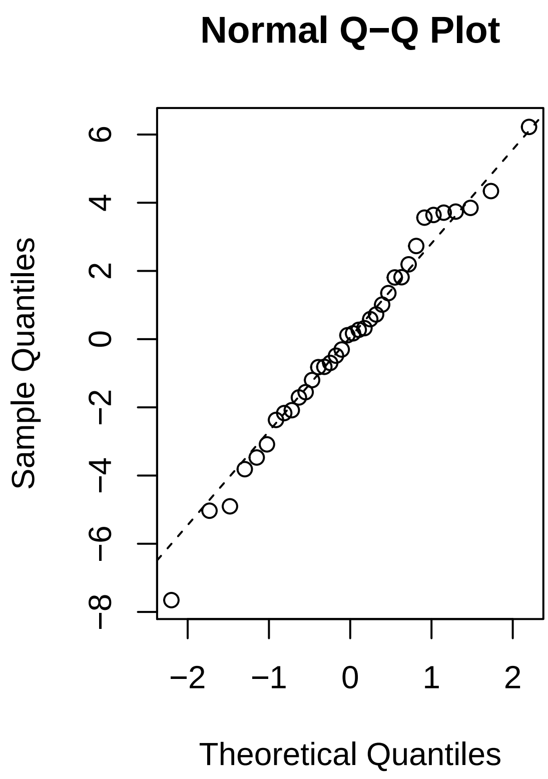

##### FSQ

###### Unadjusted

This is the linear model:

```
model <- lm(FSQ_4 ~ AAT_3 + EAST_3, data = dat[dat$WideData == 1,])
```

These are the results:

```
summary(model)
```

```
##
## Call:
## lm(formula = FSQ_4 ~ AAT_3 + EAST_3, data = dat[dat$WideData ==
##     1, ])
##
## Residuals:
##     Min      1Q  Median      3Q     Max
## -24.650 -12.046  -1.163  12.206  51.274
##
## Coefficients:
##             Estimate Std. Error t value Pr(>|t|)
## (Intercept) 27.95989    3.17399   8.809 3.51e-10 ***
## AAT_3       -0.04410    0.04944  -0.892   0.3788
## EAST_3      -0.18816    0.09306  -2.022   0.0514 .
```

```
## ---
## Signif. codes:  0 '***' 0.001 '**' 0.01 '*' 0.05 '.' 0.1 ' ' 1
##
## Residual standard error: 18.38 on 33 degrees of freedom
##   (2 observations deleted due to missingness)
## Multiple R-squared:  0.1193, Adjusted R-squared:  0.06592
## F-statistic: 2.235 on 2 and 33 DF,  p-value: 0.1229
```

There is no significant effect of biases.

##### Assumptions

Here are the residuals plotted against fitted values to investigate underlying heteroskedasticity and linearity assumptions:

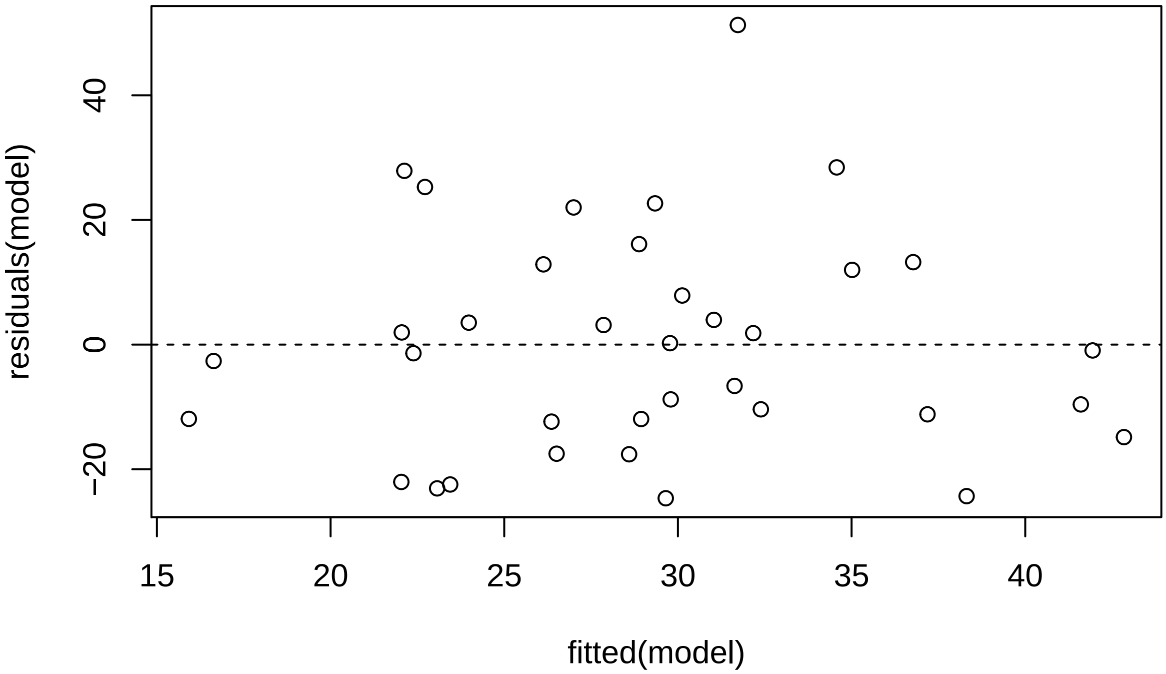

And the assumption of normality of residuals with histogram and qqplot:

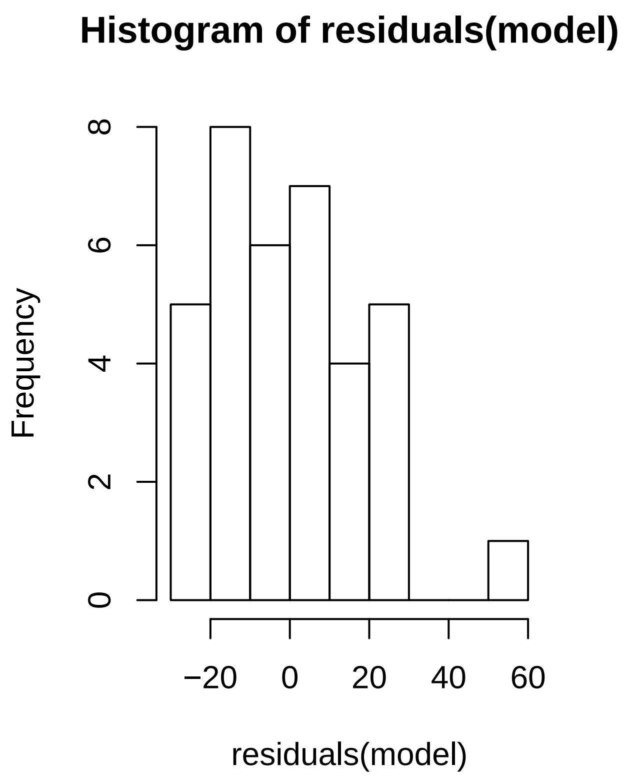

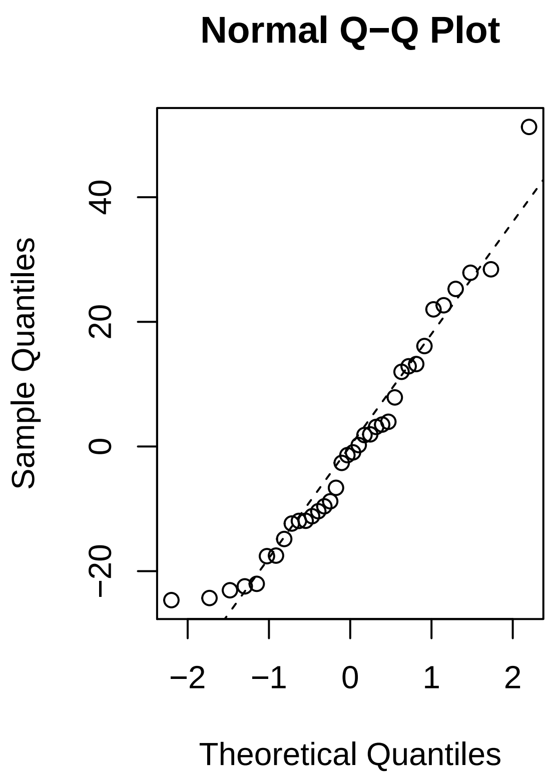

###### Adjusted

This is the linear model:

```
model <- lm(FSQ_4 ~ AAT_3 + EAST_3 + FSQ_3, data = dat[dat$WideData == 1,])
```

These are the results:

```
summary(model)
```

```
##
## Call:
## lm(formula = FSQ_4 ~ AAT_3 + EAST_3 + FSQ_3, data = dat[dat$WideData ==
##     1, ])
##
## Residuals:
##     Min      1Q  Median      3Q     Max
## -19.033  -8.331   1.550   6.829  26.344
##
## Coefficients:
##             Estimate Std. Error t value Pr(>|t|)
## (Intercept) -7.10675    5.15934  -1.377    0.178
## AAT_3        0.01285    0.03158   0.407    0.687
## EAST_3       0.02242    0.06436   0.348    0.730
## FSQ_3        0.93577    0.12730   7.351 2.33e-08 ***
## ---
```

```
## Signif. codes:  0 '***' 0.001 '**' 0.01 '*' 0.05 '.' 0.1 ' ' 1
##
## Residual standard error: 11.39 on 32 degrees of freedom
##   (2 observations deleted due to missingness)
## Multiple R-squared:  0.6724, Adjusted R-squared:  0.6417
## F-statistic:  21.9 on 3 and 32 DF,  p-value: 6.744e-08
```

There is no significant effect of biases.

###### Assumptions

Here are the residuals plotted against fitted values to investigate underlying heteroskedasticity and linearity assumptions:

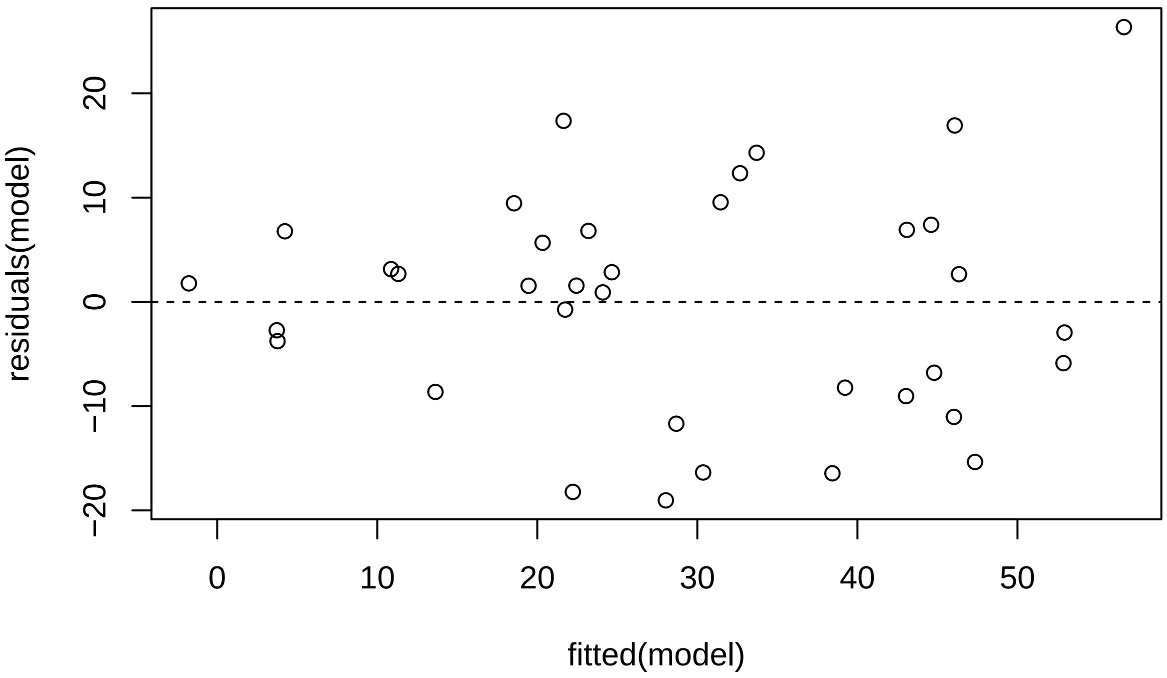

And the assumption of normality of residuals with histogram and qqplot:

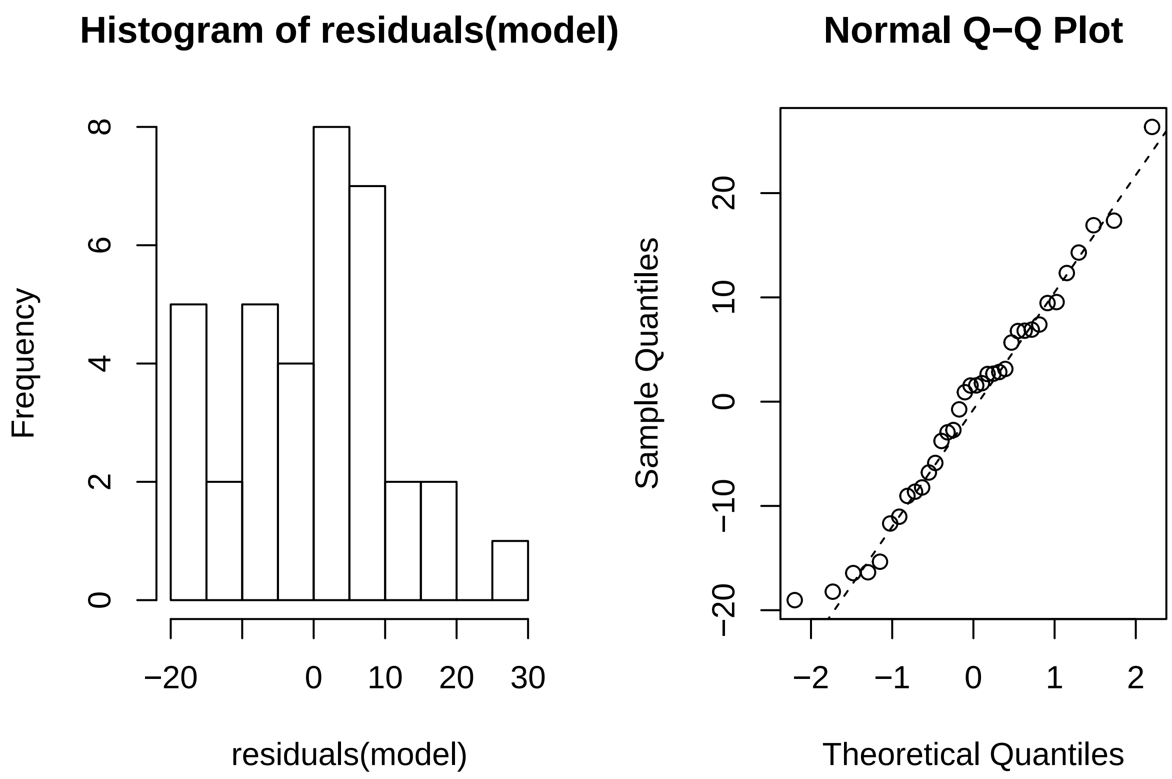

#### BAT

##### Unadjusted

This is the linear model:

```
model <- lm(BAT_speed_4 ~ AAT_3 + EAST_3, data = dat[dat$WideData == 1,])
```

These are the results:

```
summary(model)
```

```
##
## Call:
## lm(formula = BAT_speed_4 ~ AAT_3 + EAST_3, data = dat[dat$WideData ==
##     1, ])
##
## Residuals:
##     Min      1Q  Median      3Q     Max
## -0.4981 -0.2357 -0.0617  0.2946  0.6263
##
## Coefficients:
##              Estimate Std. Error t value Pr(>|t|)
## (Intercept) 0.5963036  0.0567812  10.502 4.68e-12 ***
## AAT_3       0.0012840  0.0008845   1.452   0.1560
## EAST_3      0.0036202  0.0016648   2.174   0.0369 *
```

```
## ---
## Signif. codes:  0 '***' 0.001 '**' 0.01 '*' 0.05 '.' 0.1 ' ' 1
##
## Residual standard error: 0.3289 on 33 degrees of freedom
##   (2 observations deleted due to missingness)
## Multiple R-squared:  0.1555, Adjusted R-squared:  0.1044
## F-statistic: 3.039 on 2 and 33 DF,  p-value: 0.06146
```

There is a significant effect of 1-day EAST scores on the BAT.

###### Assumptions

Here are the residuals plotted against fitted values to investigate underlying heteroskedasticity and linearity assumptions:

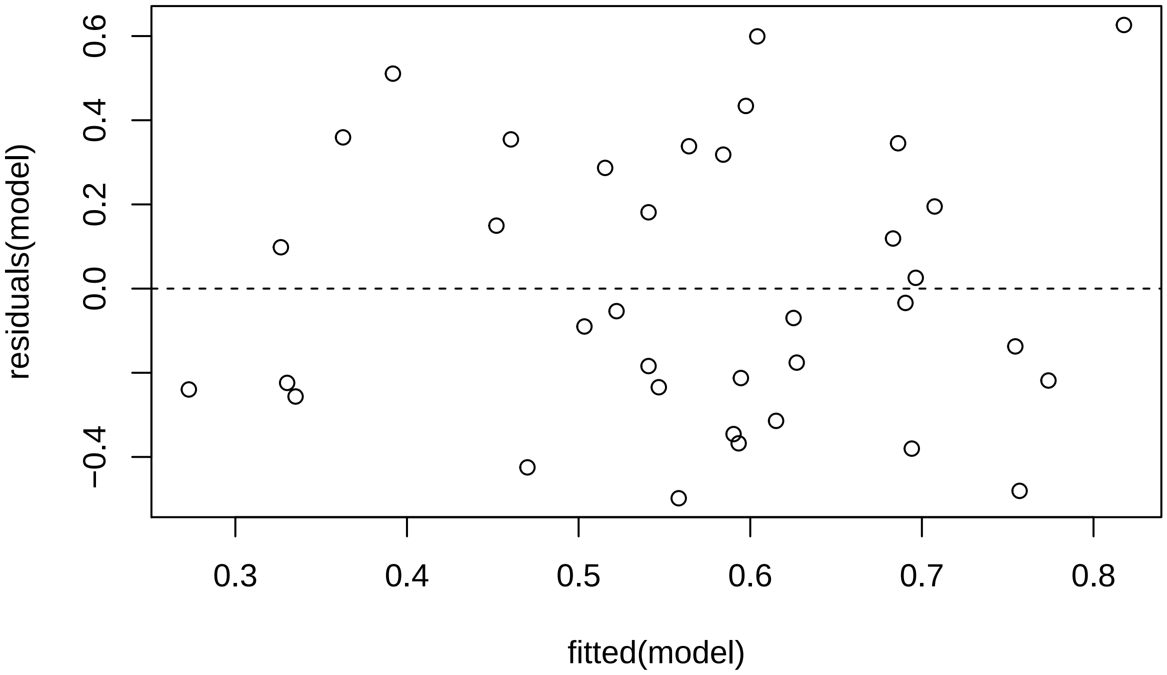

And the assumption of normality of residuals with histogram and qqplot:

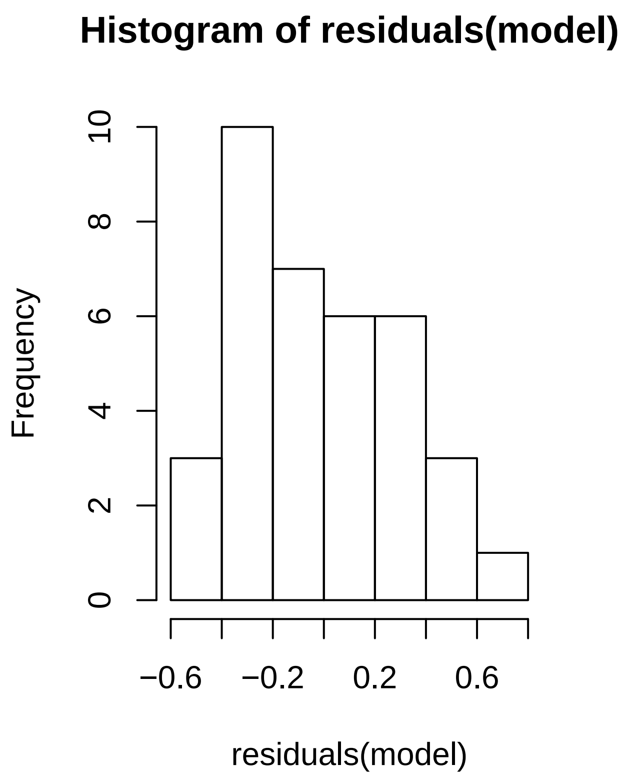

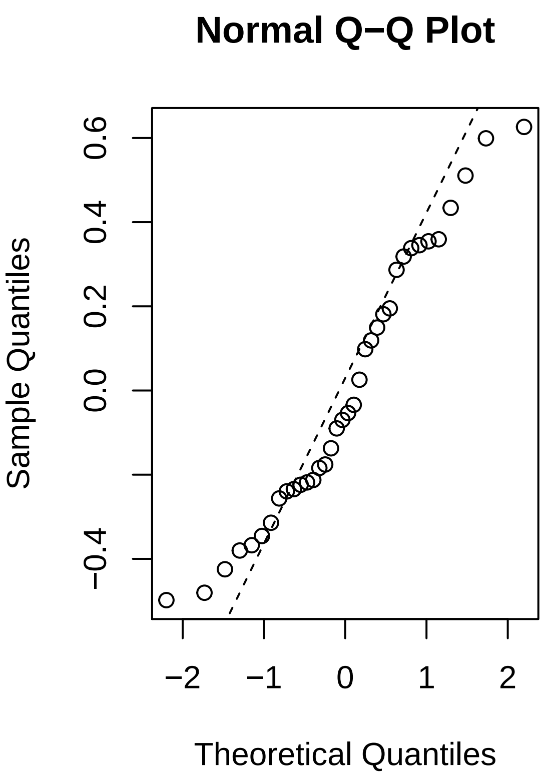

###### Adjusted

This is the linear model:

```
model <- lm(BAT_speed_4 ~ AAT_3 + EAST_3 + BAT_speed, data = dat[dat$WideData == 1,])
```

These are the results:

```
summary(model)
```

```
##
## Call:
## lm(formula = BAT_speed_4 ~ AAT_3 + EAST_3 + BAT_speed, data = dat[dat$WideData ==
##     1, ])
##
## Residuals:
##      Min       1Q   Median       3Q      Max
## -0.55967 -0.20456 -0.06366  0.18937  0.51484
##
## Coefficients:
##             Estimate Std. Error t value Pr(>|t|)
## (Intercept) 0.4599742  0.0930766   4.942 2.98e-05 ***
## AAT_3       0.0010163  0.0008415   1.208   0.2369
## EAST_3      0.0029723  0.0016491   1.802   0.0819 .
## BAT_speed   0.6695242  0.3211585   2.085   0.0460 *
## ---
```

```
## Signif. codes:  0 '***' 0.001 '**' 0.01 '*' 0.05 '.' 0.1 ' ' 1
##
## Residual standard error: 0.3077 on 29 degrees of freedom
##   (5 observations deleted due to missingness)
## Multiple R-squared:  0.2778, Adjusted R-squared:  0.203
## F-statistic: 3.718 on 3 and 29 DF,  p-value: 0.02237
```

There is no significant bias effect.

###### Assumptions

Here are the residuals plotted against fitted values to investigate underlying heteroskedasticity and linearity assumptions:

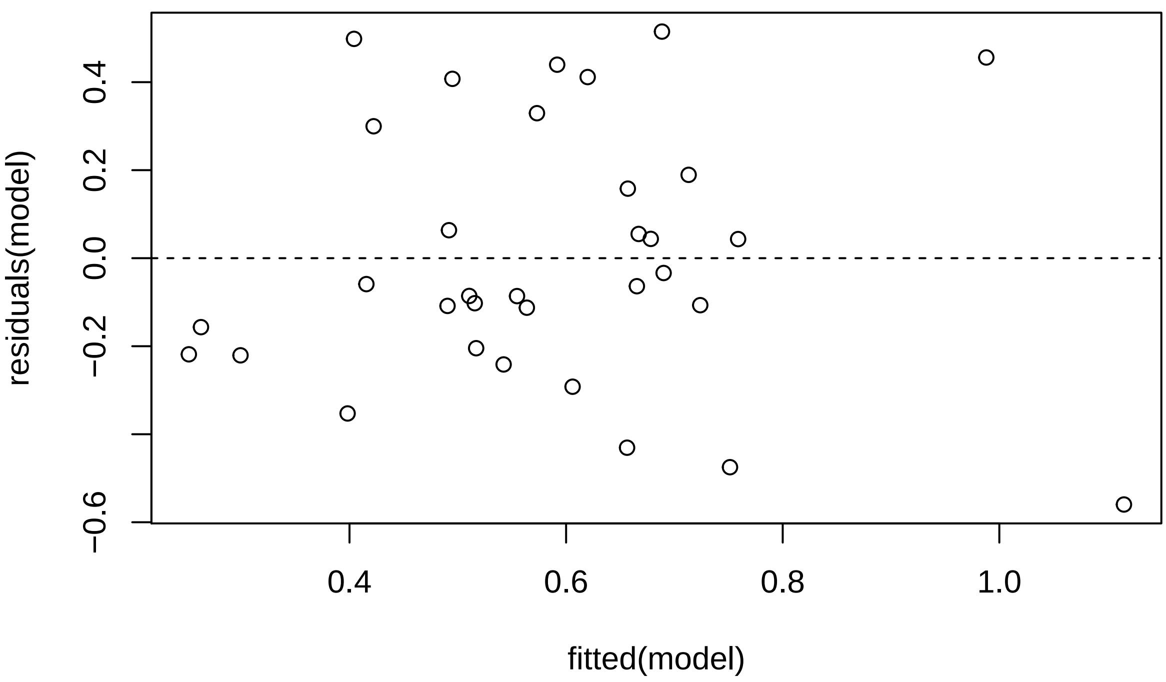

And the assumption of normality of residuals with histogram and qqplot:

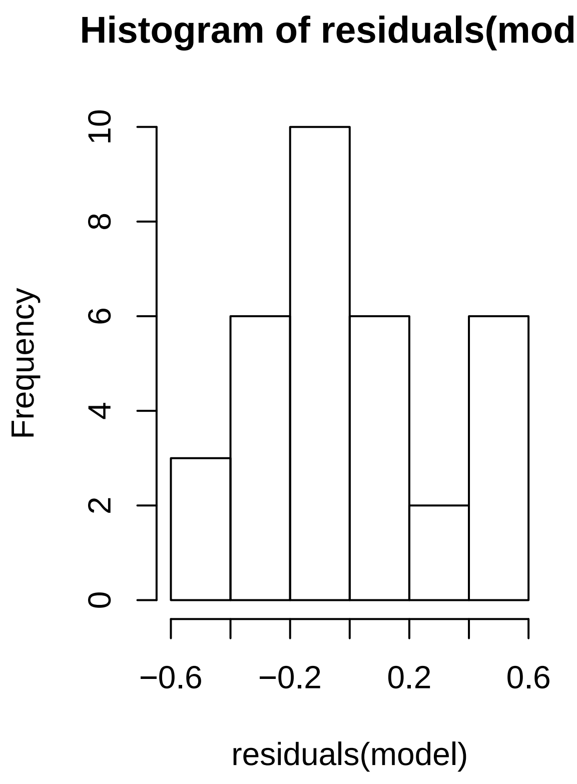

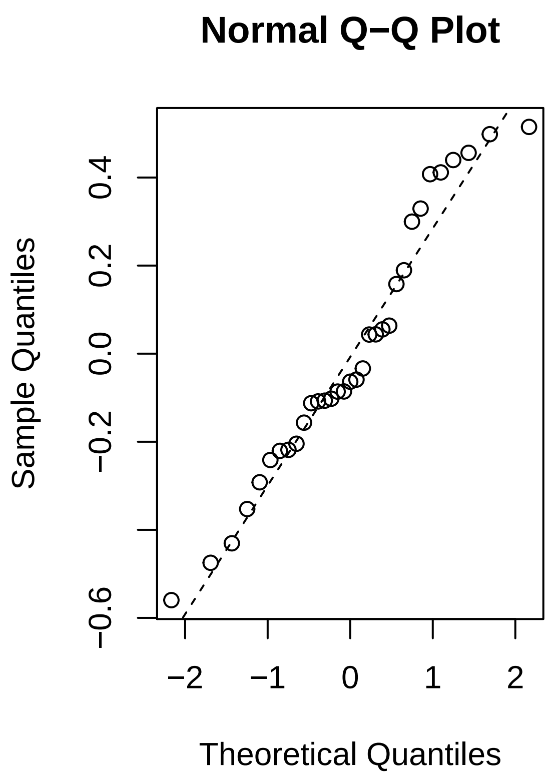

##### Correlation Matrix

Lastly, a correlation matrix is created including spider anxiety measures at 1-day follow-up as well as at 1-month follow-up. This will hopefully aid exploratory insight into potential processes of clinical change:

```r
round(cor(dat[, c("SAS_3", "FSQ_3", "BAT_speed_3", "EAST_3", "AAT_3",
                  "SAS_4", "FSQ_4", "BAT_speed_4", "EAST_4", "AAT_4")],
          use = "pairwise.complete.obs", method = "pearson"), 2)
```

```
##             SAS_3 FSQ_3 BAT_speed_3 EAST_3 AAT_3 SAS_4 FSQ_4 BAT_speed_4
## SAS_3        1.00  0.83       -0.28  -0.17 -0.25  0.82  0.60       -0.52
## FSQ_3        0.83  1.00       -0.31  -0.37 -0.16  0.76  0.82       -0.49
## BAT_speed_3 -0.28 -0.31        1.00   0.24  0.15 -0.40 -0.29        0.79
## EAST_3      -0.17 -0.37        0.24   1.00 -0.14 -0.22 -0.29        0.27
## AAT_3       -0.25 -0.16        0.15  -0.14  1.00 -0.22 -0.10        0.19
## SAS_4        0.82  0.76       -0.40  -0.22 -0.22  1.00  0.76       -0.52
## FSQ_4        0.60  0.82       -0.29  -0.29 -0.10  0.76  1.00       -0.45
## BAT_speed_4 -0.52 -0.49        0.79   0.27  0.19 -0.52 -0.45        1.00
## EAST_4      -0.29 -0.49        0.21   0.62  0.01 -0.20 -0.31        0.27
## AAT_4       -0.08  0.07        0.05  -0.14  0.11  0.02  0.08        0.16
##             EAST_4 AAT_4
## SAS_3        -0.29 -0.08
## FSQ_3        -0.49  0.07
## BAT_speed_3   0.21  0.05
## EAST_3        0.62 -0.14
## AAT_3         0.01  0.11
```

```
## SAS_4        -0.20  0.02
## FSQ_4        -0.31  0.08
## BAT_speed_4   0.27  0.16
## EAST_4        1.00 -0.14
## AAT_4        -0.14  1.00
```

This correlation matrix is also saved in apa format using the *apaTables* package:

```r
apa.cor.table(dat[, c("SAS_3", "FSQ_3", "BAT_speed_3", "EAST_3", "AAT_3",
                      "SAS_4", "FSQ_4", "BAT_speed_4", "EAST_4", "AAT_4")],
              filename = "SpiderFear_CorMatrix.doc", table.number = 1)
```

```
##
##
## Table 1
##
## Means, standard deviations, and correlations with confidence intervals
##
##
##   Variable      M      SD    1            2            3
##   1. SAS_3      12.76  4.44
##
##   2. FSQ_3      38.13  17.15 .83**
##                              [.78, .88]
##
##   3. BAT_speed_3 0.37  0.27  -.28**       -.31**
##                              [-.42, -.13] [-.45, -.16]
##
##   4. EAST_3     -3.09  32.75 -.17*        -.37**       .24**
##                              [-.32, -.01] [-.50, -.22] [.08, .38]
##
##   5. AAT_3      -15.06 62.79 -.25**       -.16*        .15
##                              [-.39, -.09] [-.32, -.00] [-.02, .30]
##
##   6. SAS_4      10.95  5.13  .82**        .76**        -.40**
##                              [.76, .87]   [.69, .82]   [-.53, -.26]
##
##   7. FSQ_4      28.67  18.57 .60**        .82**        -.29**
##                              [.49, .69]   [.76, .87]   [-.43, -.14]
##
##   8. BAT_speed_4 0.57  0.35  -.52**       -.49**       .79**
##                              [-.63, -.39] [-.61, -.36] [.73, .85]
##
##   9. EAST_4     0.20   30.42 -.29**       -.49**       .21*
##                              [-.43, -.14] [-.60, -.35] [.05, .36]
##
##   10. AAT_4     -1.22  43.33 -.08         .07          .05
##                              [-.23, .09]  [-.09, .23]  [-.11, .21]
##
##   4            5            6            7            8
##
##
##
##
##
```

```
##
##
##
##
##
##
##   -.14
##   [-.30, .03]
##
##   -.22**        -.22**
##   [-.37, -.06] [-.37, -.06]
##
##   -.29**        -.10          .76**
##   [-.43, -.14] [-.26, .06]  [.69, .82]
##
##   .27**         .19*          -.52**        -.45**
##   [.11, .41]    [.02, .34]    [-.62, -.39] [-.57, -.31]
##
##   .62**         .01           -.20*         -.31**        .27**
##   [.51, .71]    [-.16, .18]  [-.35, -.04] [-.45, -.16] [.12, .42]
##
##   -.14          .11           .02           .08           .16
##   [-.29, .02]  [-.05, .27]  [-.14, .18]  [-.08, .23]  [-.00, .31]
##
##   9
##
##
##
##
##
##
##
##
##
##
##
##
##
##
##
##
##
##
##
##
##
##
##
##
##
##
##
##   -.14
##   [-.30, .02]
##
```

```
##
## Note. M and SD are used to represent mean and standard deviation, respectively.
## Values in square brackets indicate the 95% confidence interval.
## The confidence interval is a plausible range of population correlations
## that could have caused the sample correlation (Cumming, 2014).
## * indicates p < .05. ** indicates p < .01.
##
```

### References

See below references for all packages used:

```r
citation("tidyverse")
```

```
##
## To cite package 'tidyverse' in publications use:
##
##   Hadley Wickham (2017). tidyverse: Easily Install and Load the
##   'Tidyverse'. R package version 1.2.1.
##   https://CRAN.R-project.org/package=tidyverse
##
## A BibTeX entry for LaTeX users is
##
##   @Manual{,
##     title = {tidyverse: Easily Install and Load the 'Tidyverse'},
##     author = {Hadley Wickham},
##     year = {2017},
##     note = {R package version 1.2.1},
##     url = {https://CRAN.R-project.org/package=tidyverse},
##   }
```

```r
citation("lme4")
```

```
##
## To cite lme4 in publications use:
##
##   Douglas Bates, Martin Maechler, Ben Bolker, Steve Walker (2015).
##   Fitting Linear Mixed-Effects Models Using lme4. Journal of
##   Statistical Software, 67(1), 1-48. doi:10.18637/jss.v067.i01.
##
## A BibTeX entry for LaTeX users is
##
##   @Article{,
##     title = {Fitting Linear Mixed-Effects Models Using {lme4}},
##     author = {Douglas Bates and Martin M{\"a}chler and Ben Bolker and Steve Walker},
##     journal = {Journal of Statistical Software},
##     year = {2015},
##     volume = {67},
##     number = {1},
##     pages = {1--48},
##     doi = {10.18637/jss.v067.i01},
##   }
```

```r
citation("influence.ME")
```

```
##
## To cite package `influence.ME' in publications, please use:
##
##   Rense Nieuwenhuis, Manfred te Grotenhuis and Ben Pelzer (2012).
##   influence.ME: Tools for Detecting Influential Data in Mixed
##   Effects Models. R Journal, 4(2): pp. 38-47.
##
## A BibTeX entry for LaTeX users is
##
##   @Article{,
##     title = {influence.ME: Tools for Detecting Influential Data in Mixed Effects Models},
##     author = {Rense Nieuwenhuis and Manfred {Te Grotenhuis} and Ben Pelzer},
##     year = {2012},
##     journal = {R Journal},
##     volume = {4},
##     number = {2},
##     pages = {38-47},
##   }
```

```r
citation("apaTables")
```

```
##
## To cite package 'apaTables' in publications use:
##
##   David Stanley (2018). apaTables: Create American Psychological
##   Association (APA) Style Tables. R package version 2.0.5.
##   https://CRAN.R-project.org/package=apaTables
##
## A BibTeX entry for LaTeX users is
##
##   @Manual{,
##     title = {apaTables: Create American Psychological Association (APA) Style Tables},
##     author = {David Stanley},
##     year = {2018},
##     note = {R package version 2.0.5},
##     url = {https://CRAN.R-project.org/package=apaTables},
##   }
```
